# Phylogenetic systematics and evolution of the spider infraorder Mygalomorphae using genomic scale data

**DOI:** 10.1101/531756

**Authors:** Vera Opatova, Chris A. Hamilton, Marshal Hedin, Laura Montes de Oca, Jiří Král, Jason E. Bond

## Abstract

The Infraorder Mygalomorphae is one of the three main lineages of spiders comprising over 3,000 nominal species. This ancient group has a world-wide distribution that includes among its ranks large and charismatic taxa such as tarantulas, trapdoor spiders, and highly venomous funnel web spiders. Based on past molecular studies using Sanger-sequencing approaches, numerous mygalomorph families (e.g., Hexathelidae, Ctenizidae, Cyrtaucheniidae, Dipluridae and Nemesiidae) have been identified as non-monophyletic. However, these data were unable to sufficiently resolve the higher-level (intra- and interfamilial) relationships such that the necessary changes in classification could be made with confidence. Here we present a comprehensive phylogenomic treatment of the spider infraorder Mygalomorphae. We employ 472 loci obtained through Anchored Hybrid Enrichment to reconstruct relationships among all the mygalomorph spider families and estimate the timeframe of their diversification. We sampled all currently recognized families, which has allowed us to assess their status, and as a result, propose a new classification scheme. Our generic-level sampling has also provided an evolutionary framework for revisiting questions regarding silk use in mygalomorph spiders. The first such analysis for the group within a strict phylogenetic framework shows that a sheet web is likely the plesiomorphic condition for mygalomorphs, as well as providing hints to the ancestral foraging behavior for all spiders. Our divergence time estimates, concomitant with detailed biogeographic analysis, suggest that both ancient continental-level vicariance and more recent dispersal events have played an important role in shaping modern day distributional patterns. Based on our results, we relimit the generic composition of the Ctenizidae, Cyrtaucheniidae, Dipluridae and Nemesiidae. We also elevate five subfamilies to family rank: Anamidae (NEW RANK), Euagridae (NEW RANK), Ischnothelidae (NEW RANK), Pycnothelidae (NEW RANK), and Bemmeridae (NEW RANK). The three families Entypesidae (NEW FAMILY), Microhexuridae (NEW FAMILY), and Stasimopidae (NEW FAMILY) are newly proposed. Such a major rearrangement in classification, recognizing eight newly established family-level rank taxa, is the largest the group has seen in over three decades since Raven’s (1985) taxonomic treatment.

---

Spiders placed in the infraorder Mygalomorphae are a charismatic assemblage of taxa that includes among its ranks the tarantulas, trapdoor spiders, and also some of the most venomous species, like the Sydney funnel web spider and its close relatives (Hedin et al. 2018). Today, the infraorder comprises 20 families, divided into 354 genera, with 3,006 species (World Spider Catalog 2018; Table 1). The group is ancient, known to occur in the fossil record since the Middle Triassic (Selden and Gall 1992), but with origins estimated further back into the Carboniferous over 300 million years ago (Ayoub et al. 2007; Starrett et al. 2013; Garrison et al. 2016). Their ancient origins and concomitant sedentary nature provide a rich biogeographical context for studying geological-scale continental vicariant events (Raven 1980; Hedin and Bond 2006; Opatova et al. 2013; Opatova and Arnedo 2014a). Not surprisingly for such an ancient lineage, mygalomorph spider morphology is complicated; they have retained a number of features considered primitive for spiders, such as a simple silk spinning system, two pairs of book lungs and paraxial cheliceral arrangement, yet are also relatively homogenous when compared to the overall morphological diversity observed among their araneomorph counterparts (Hendrixson and Bond 2009). Although such striking homogeneity presents significant challenges for morphology-based taxonomy, mygalomorphs have become a noteworthy system for studying allopatric speciation and species crypsis (Bond et al. 2001; Bond and Stockman 2008; Satler et al. 2013; Hamilton et al. 2014; Opatova and Arnedo 2014b; Hedin et al. 2015; Leavitt et al. 2015; Castalanelli et al. 2017; Rix et al. 2017b; Starrett et al. 2018).

**Table 1.**
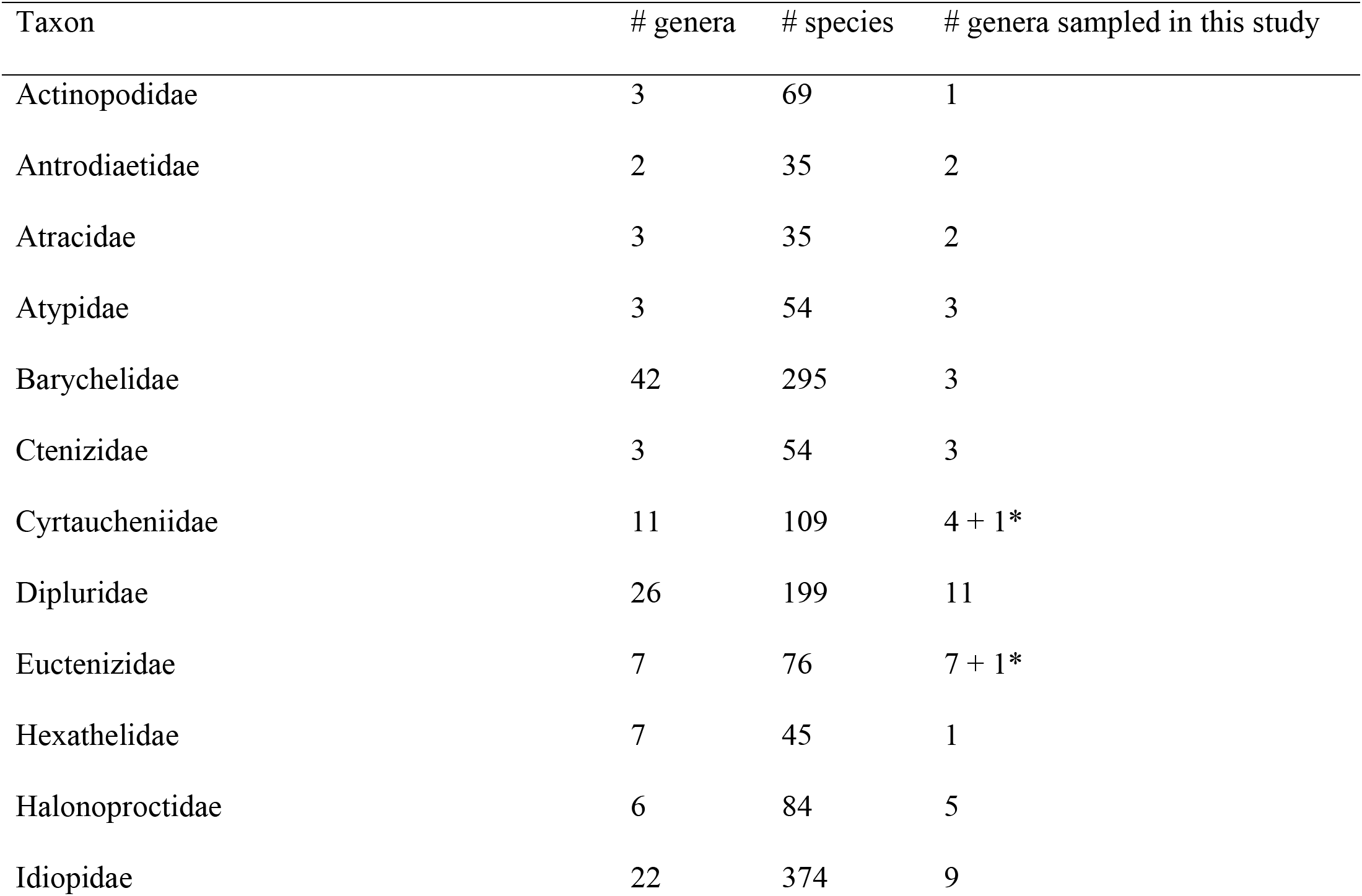
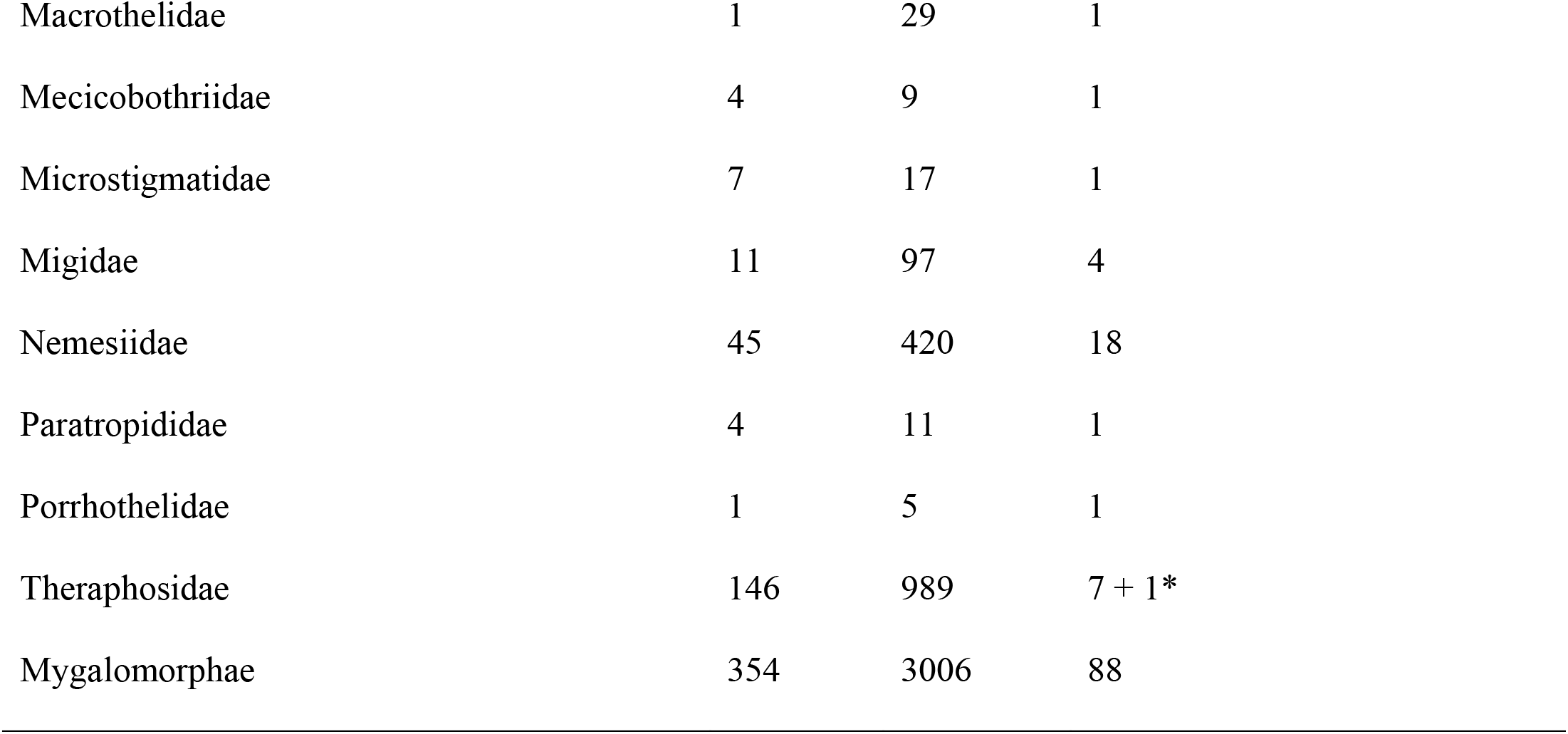
Taxon counts obtained from the World Spider Catalog (2018), undescribed and undetermined genera marked with *.

For less than two decades, since the first application of molecular systematics to questions regarding mygalomorph phylogeny (Hedin and Bond 2006), it has been generally acknowledged that the current system of classification is replete with problems that include numerous instances of family-level para and/or polyphyly. In short, much of today’s classification scheme dates back to Raven (1985) Fig. 1a, which first applied cladistic thinking towards evaluating family-level relationships on the basis of morphological character argumentation. Post Raven (1985), a number of authors continued to explore morphological characters as evidence for broad scale familial-level relationships and identity (Eskov and Zonshtein 1990), further facilitated by the availability of computational phylogenetic inference (Goloboff 1993; Goloboff 1995; Bond and Opell 2002; Fig. 1b). These morphological studies were followed shortly thereafter by a number of early molecular treatments (Hedin and Bond 2006; Ayoub et al. 2007; Fig. 2a). The first major taxonomic changes to Raven’s classification followed in 2012 (Bond et al. 2012; Fig. 2b) with the elevation of the subfamily Euctenizinae to family status. A number of subsequent studies employing Sanger-sequencing based approaches continued to make progress in exploring relationships between closely related taxa (Hamilton et al. 2014; Opatova et al. 2016; Mora et al. 2017; Rix et al. 2017a; Ortiz et al. 2018; Starrett et al. 2018) and within families (Opatova et al. 2013; Opatova and Arnedo 2014a; Harrison et al. 2017; Rix et al. 2017c; Harvey et al. 2018; Lüddecke et al. 2018; Turner et al. 2018). Although Sanger approaches have been fruitful in terms of advancing our understanding of some relationships, it has been long recognized that ribosomal RNA genes (e.g., 28S and 18S), elongation factors, and mitochondrial genes, are often of limited utility in the context of deep scale divergences of mygalomorph spiders (Hedin and Bond 2006; Ayoub et al. 2007; Bond et al. 2012).

**Figure 1.**
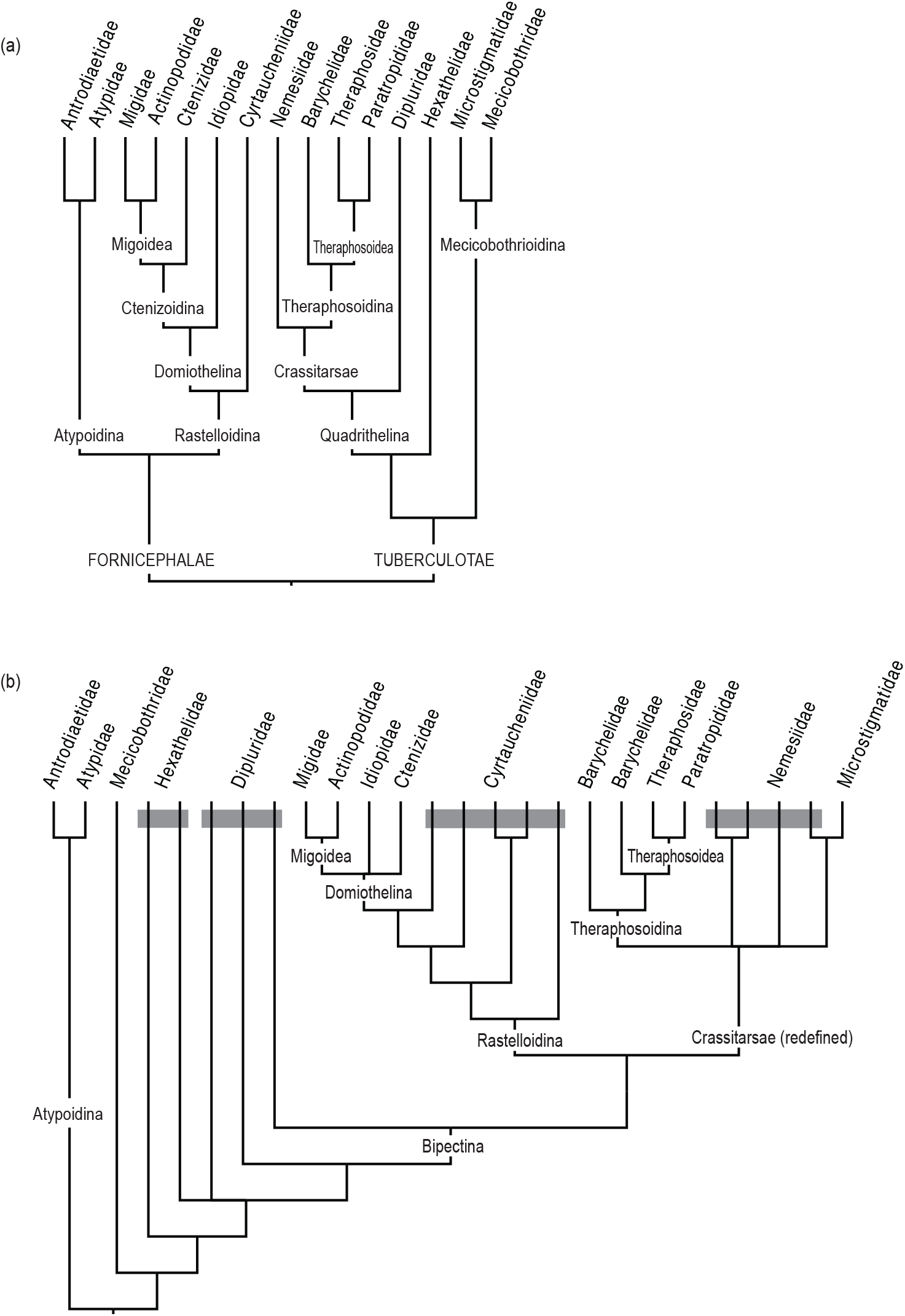
Phylogenetic relationships of mygalomorph families reconstructed from morphological characters: (a) Raven’s classification scheme (Raven 1985), (b) results of the first cladistic analyses performed by Goloboff (1993). Poly- and paraphyletic lineages are indicated by grey boxes.

**Figure 2.**
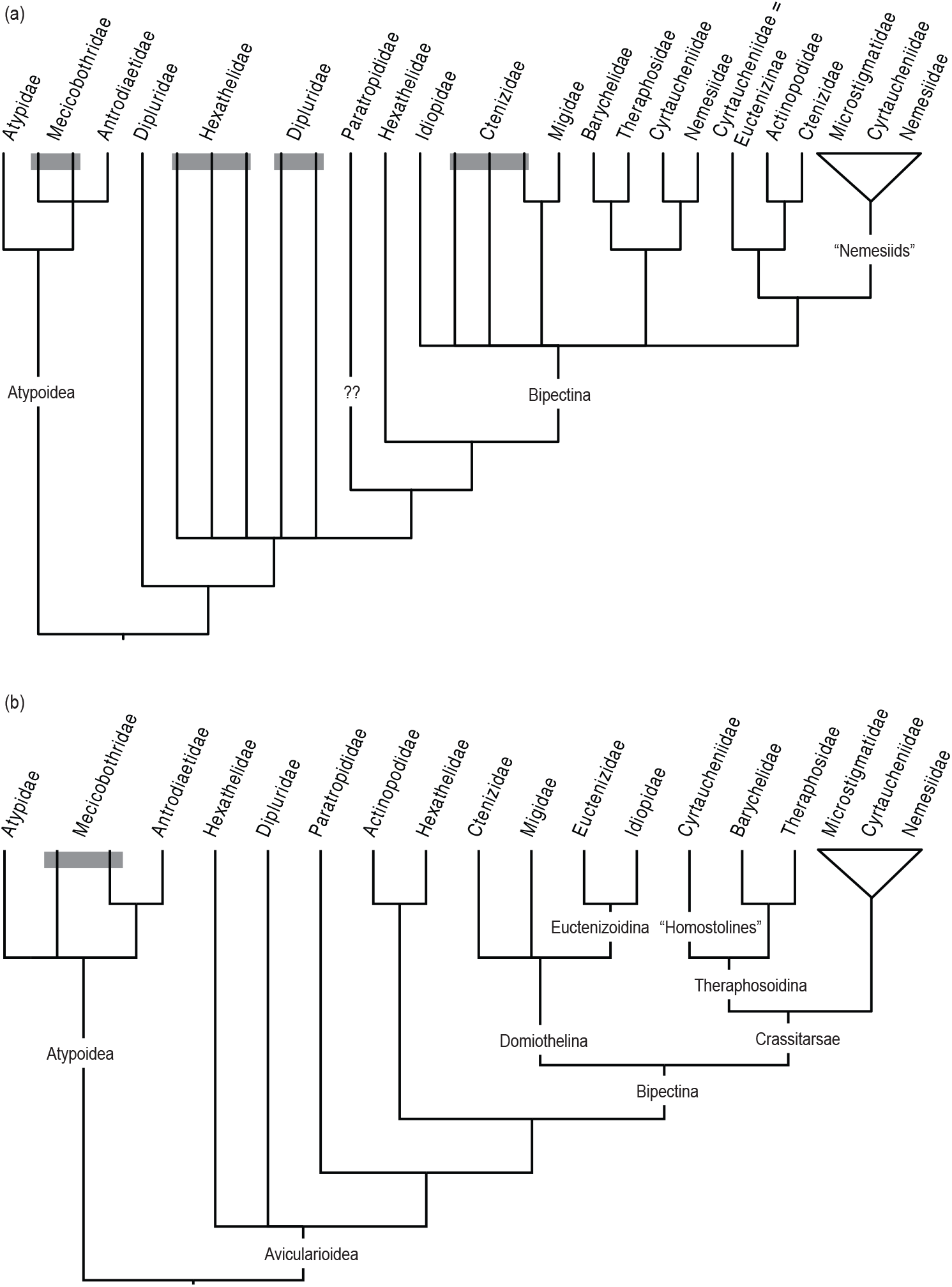
Phylogenetic relationships of mygalomorph families: (a) inferred from ribosomal data (*18S*, *28S*) in the first molecular analyses of the infraorder (Hedin and Bond 2006), (b) inferred in the “total evidence” approach combining three loci (*18S*, *28S*, *EF-1g*) and morphological data (Bond et al. 2012). Poly- and paraphyletic lineages are indicated by grey boxes.

Within the last several years, the application of phylogenomic approaches has helped to transform spider systematics (Bond et al. 2014; Fernández et al. 2014; Garrison et al. 2016; Hamilton et al. 2016b; Hedin et al. 2018; Kuntner et al. 2018; Wood et al. 2018; Hedin et al. 2019), by providing the data and impetus for making major changes in mygalomorph spider classification. Specifically, targeted capture approaches using anchored hybrid enrichment (AHE; Lemmon et al. 2012) and ultra-conserved elements (UCE; Faircloth et al. 2012) are now being applied to questions at multiple hierarchical levels across the spider tree of life (Hamilton et al. 2016a; Hamilton et al. 2016b; Maddison et al. 2017; Chamberland et al. 2018; Godwin et al. 2018; Hedin et al. 2018) and other arachnids (Starrett et al. 2017; Sharma et al. 2018). Implementation of genomic-based methods seemingly overcomes the perceived limitations of the traditional Sanger loci, enabling mygalomorph systematists to formally address a number of long-standing issues like the polyphyletic nature of ctenizids and hexathelids (Godwin et al. 2018; Hedin et al. 2018). In both cases, phylogenomic results stabilized the taxonomic position of genera that were notoriously challenging to place and established new familial ranks (Halonoproctidae (Godwin et al. 2018), Atracidae, Macrothelidae and Porrhothelidae (Hedin et al. 2018)). Although these smaller scale studies have made important advances in our understanding of mygalomorph phylogeny and classification, they have all thus far lacked the broad scale phylogenetic context instrumental to understanding relationships spanning the entire infraorder.

We present here the most comprehensive phylogenomic treatment of Mygalomorphae conducted to date. Our analysis includes broad sampling of taxa from all currently recognized families (88 genera, 111 species), allowing us to assess the monophyly of a number of key families (e.g., Cyrtaucheniidae, Nemesiidae, Dipluridae, Ctenizidae) and address some long-standing taxonomic problems. Our generic-level sampling also provides an evolutionary framework for revisiting questions regarding silk use in non-araneomorph spiders. We explore questions regarding the timing and biogeographical context of mygalomorph spider diversification, and explicitly test a number of hypotheses related to the timing and origin of major mygalomorph lineages, the use of silk in mygalomorph prey capture, and the monophyly of the major mygalomorph higher-level groups and families.

## Materials & Methods

We sampled a total of 113 taxa, 111 ingroup specimens representing 88 genera from all 20 currently recognized mygalomorph families (Table 1) and two representatives of the suborder Mesothelae (*Liphisthius* and *Vinathela*, family Liphistiidae) as outgroups to root phylogenies. Our sampling represents approximately 25% of the infraorder’s known generic diversity and about 47% if the two most diverse mygalomorph families, Barychelidae and Theraphosidae, are excluded from the calculation. Approximately 60% of AHE sequence data was newly generated for this study, additional sequences from Hamilton et al. (2016b) and Godwin et al. (2018) were included. Taxon sampling is summarized in Supplemental Table S1; all statistics related to family and generic-level species diversity are extracted from the World Spider Catalog (https://wsc.nmbe.ch/) accessed November 2018.

Whole genomic DNA was extracted using the DNeasy Blood and Tissue Kit (Qiagen), following the manufacturer’s guidelines; to ensure the complete digestion of RNA, RNase A was added to the mix after the tissue lysis step. Library preparation, enrichment, and sequencing were performed at the Center for Anchored Phylogenomics at Florida State University (http://anchoredphylogeny.com/) following the methods described in Lemmon et al. (2012), Hamilton et al. (2016b) and Godwin et al. (2018). During the development of the spider AHE loci, a mixture of spider genome sequencing and transcriptomes were utilized to build scaffolds from which probes were designed in conserved anchor regions neighboring variable flanking regions, creating a set of probes that include exons, introns, intergenic, and conserved regions of the genome. Initial probe design was based on the core arthropod orthologs and the targeted loci were then found in the spider reference genomic datasets (both spider genomes and transcriptomes). Loci that did not contain both *Aliatypus* (Antrodiaetidae) and *Aphonopelma* (Theraphosidae) were not considered for further usage. After sequencing, all reads were assembled into contigs following Prum et al. (2015), but using references derived from the *Ixodes* (a tick), *Hypochilus* (araneomorph), *Aphonopelma*, and *Aliatypus* (mygalomorphs) sequences used for probe design. Homologous nucleotide sequence sets were produced by grouping filtered consensus sequences across individuals by target locus. Orthologous groups were then determined for each target locus by a) employing a pairwise-distance measure computed for pairs of homologs; b) a neighbor-joining clustering algorithm to cluster the consensus sequences into orthologous sets; c) the identified sequences are then grouped into a cluster; d) steps A-C are repeated, treating each cluster as a sequence by using the average distance between the cluster and other clusters/sequence when assessing distance. This approach efficiently divides the homologs into two or more orthologous sets, allowing us to identify duplications occurring before the ancestor of the taxonomic group being analyzed. For duplications occurring within the taxonomic group under investigation, this approach tends to produce a single cluster containing one sequence per taxon and a second cluster containing sequences from those taxa that are descendants from the ancestor in which the duplication occurred.

Following the targeting of the 585 loci within the Spider Probe Kit v1 (Hamilton et al. 2016b), 472 loci were recovered and assembled. All loci were aligned with MAFFT 7.402 (Katoh and Standley 2013) using the L-INS-i algorithm (--localprior and --maxiterate 1000 flags). The alignments were scored for accuracy in ALISCORE (Misof and Misof 2009) and ambiguously aligned positions were removed with ALICUT (Kück 2009). Individual loci were concatenated with FASconCAT (Kück and Longo 2014) yielding a supermatrix of 93,410 nucleotides for 111 ingroup and two outgroup taxa (hereafter “All_taxa”); all alignments were visually inspected in Geneious 10.1.3 (Biomatters 2017) prior to concatenation.

Two additional matrices were constructed to assess the effect of taxon removal or sequence data addition on tree topology and support values. First, terminals representing the family Paratropididae were removed from the “All_taxa” (hereafter referred to as “No_Paratropis”). The family was recovered as a “stand-alone” lineage with unresolved and unstable placement (Hedin and Bond 2006; Bond et al. 2012), potentially causing conflict in topology and lowering the support of deeper nodes. To test the effect of data addition on node support and the unresolved placement of Stasimopidae, we combined our AHE loci with pre-existing transcriptomic data (Garrison et al. 2016). To maximize the taxon sampling, we extracted the mygalomorph terminals from “BCC-75” matrix (Garrison et al. 2016) and combined them into the “DNAAA_matrix” with corresponding terminals represented in our AHE dataset, in two instances, forming chimaeras at generic level (Supplemental Table S2). In order to maintain a relevant phylogenetic framework, we also retained additional individuals representing key lineages/clades that were only represented by AHE data. Terminals extracted from the “BCC-75” matrix (Garrison et al. 2016) were parsed into individual loci using BIOPYTHON (Hamelryck and Manderick 2003; Cock et al. 2009). Resulting 1675 transcriptomic loci were blasted (tblastn command) in Geneious (Biomatters 2017) against a custom AHE locus database comprising nucleotide sequence data prior to ALICUT treatment. Amino acid loci that received an e-value of < 1e^-10^; that is, the loci already represented in AHE dataset, were excluded from the analyses. Loci that did not contain all 20 amino acids were further removed from the dataset (per Stamatakis 2014).

Phylogenetic analyses were run on the Hopper Community Cluster at Auburn University. For the DNA matrices, partition schemes under the GTR substitution model were evaluated and selected based on AICc criterion in PartionFinder 2 (Lanfear et al. 2016) using the rcluster algorithm (Lanfear et al. 2014) with RAxML implementation (Stamatakis 2014). Partitioning by codon was not considered in the analyses because ambiguously aligned positions were removed with ALICUT. Partition schemes and models of protein evolution for the amino acid data were inferred independently using PartionFinder 2 with rcluster algorithm prior to their concatenation with the nucleotide AHE sequences into the “DNAAA_matrix”.

Maximum likelihood analyses (ML) were conducted in RAxML v 8.2.9 (Stamatakis 2014). Both “All_taxa” and “No_Paratropis” datasets were analyzed using the 191-partition scheme and an independent GTR+G model defined for each partition. The best ML tree for each matrix was selected from 1000 iterations, each starting from an independently derived parsimony-based tree. Bootstrap support was inferred from 50 replicates, determined as sufficient by the automatic bootstopping criterion (-N autoMRE flag) for both datasets (Pattengale et al. 2010). Additional ML analysis of “All_taxa” dataset was also conducted in IQ-TREE (Nguyen et al. 2014), model selection for each partition was performed by with the *ModelFinder Plus*; support values were inferred via SH-aLRT test (Anisimova et al. 2011) and ultrafast bootstrapping (both 1000 replicates).

The “DNAAA_matrix” mixed dataset combining both nucleotide and amino acid data was analyzed with 1367-partition scheme, applying an independent GTR+G model for each AHE partition and a corresponding protein model inferred by PartionFinder to each transcriptome partition. The best ML tree was selected from 1000 iterations and support was assessed from 100 bootstrap replicates (autoMRE).

Statistical significance of alternative tree topologies; that is, different, or traditional, family-level arrangements (Table 2) than those recovered in our analyses, was assessed with Approximately Unbiased (AU) topology test (Shimodaira 2002) implemented in CONSEL (Shimodaira and Hasegawa 2001). Alternative tree topologies were obtained from 1000 independent constrained searches conducted in RAxML implementing the same settings, partition scheme, and nucleotide substitution model as described above.

**Table 2.**
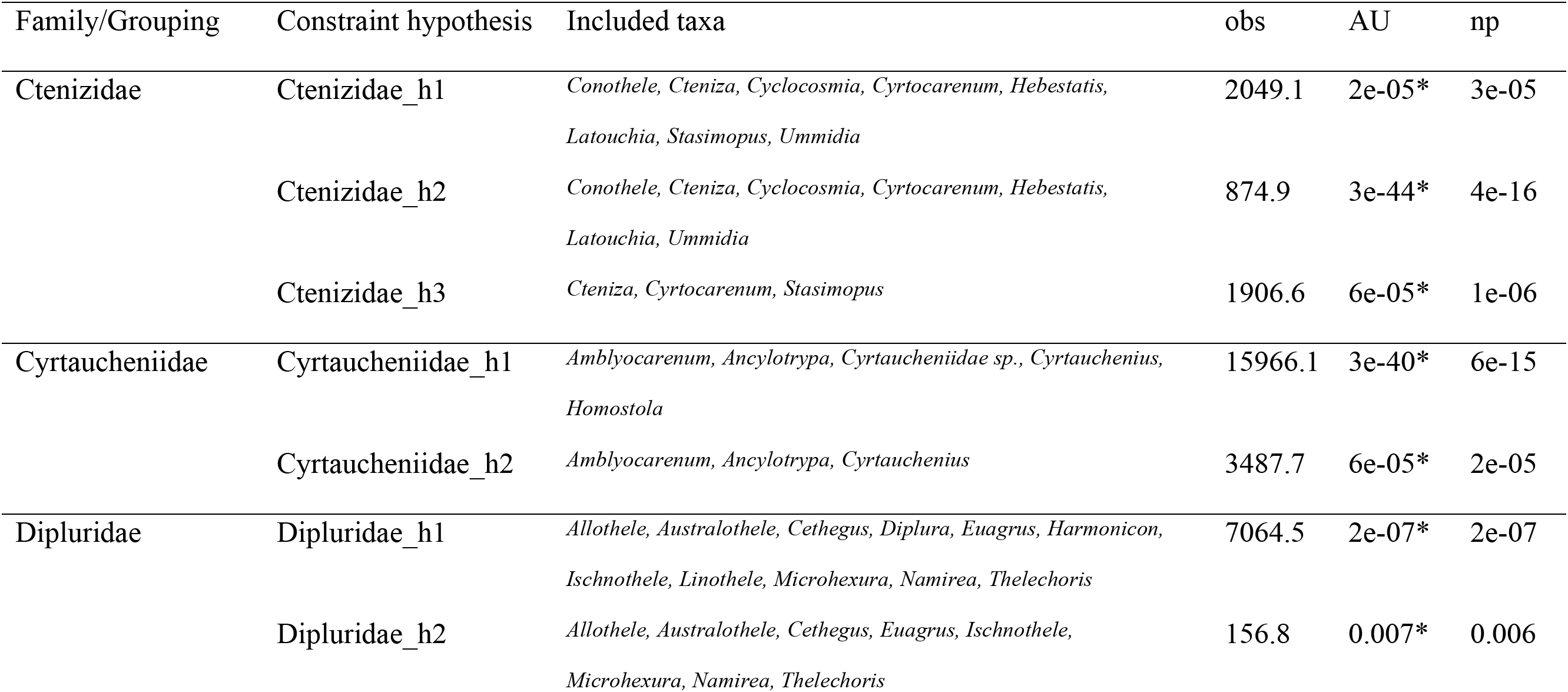
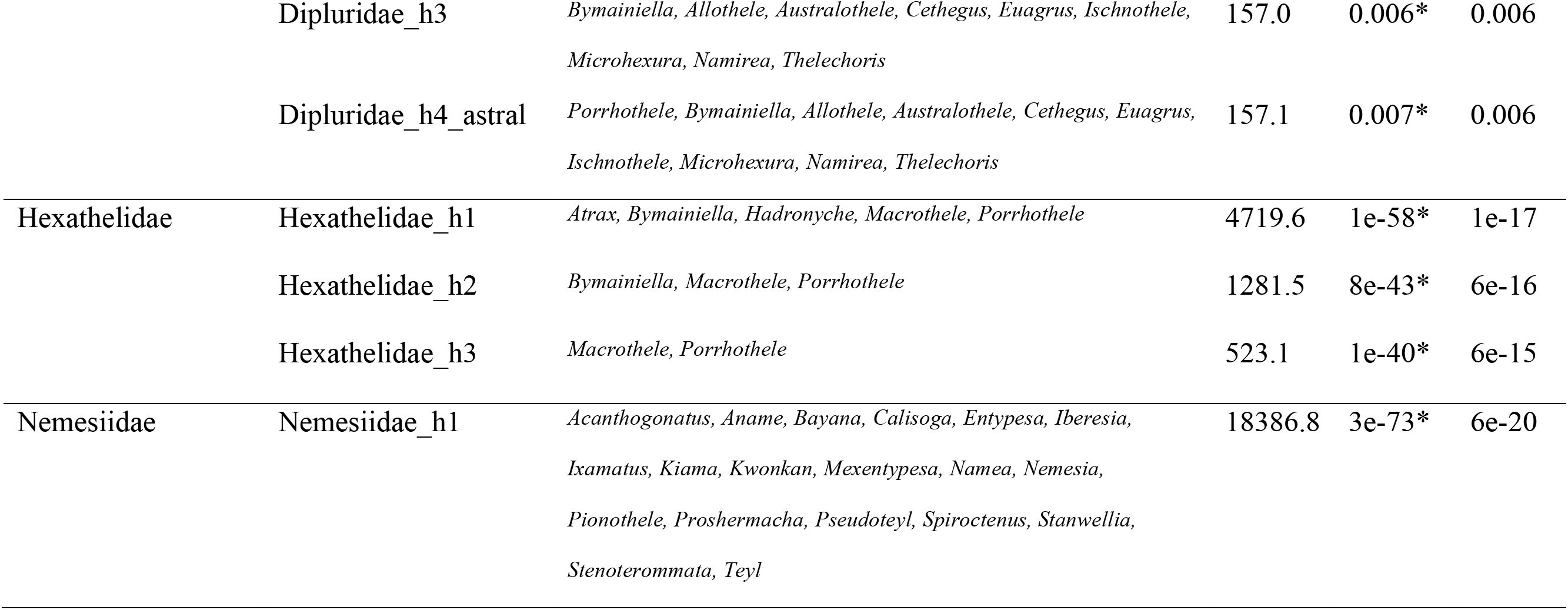
Maximum likelihood topology test results: obs, the observed log-likelihood difference to the best unconstraint topology (Fig. 3), AU, p-value of the approximately unbiased test conducted in CONSEL; np, bootstrap probability calculated from the multiscale bootstrap; significant results (P<0.05) *

Bayesian Inference (BI) analyses of the “All_taxa” dataset was conducted in ExaBayes v 1.4.1 (Aberer et al. 2014) implementing the same partition scheme and nucleotide substitution model as in the ML analyses. Two independent runs of 4×10^7^ generations with four coupled chains each, starting from a parsimony tree with resampling every 1000 generations, were run simultaneously. Standard deviation of split frequencies was monitored (<0.01), and the first 25% were discarded as a *burn-in* for the analyses. An extended majority rule consensus tree was obtained in the ExaBayes accompanying program consense (Aberer et al. 2014).

A species tree was inferred from 472 gene trees using ASTRAL v 4.11.2 (Mirarab and Warnow 2015). Node support was assessed using ASTRAL’s local posterior probabilities. Single gene trees were inferred from AHE nucleotide alignments implementing GTR+G model in RAxML; the best ML trees were selected from 1000 independent iterations for each locus individually.

Divergence times were estimated in the penalized likelihood environment (Sanderson 2002) using the treePL algorithm (Smith and O’Meara 2012) on the tree topology obtained in RAxML, and calibrated with six fossils. The optimal settings for the thorough analysis was determined with the prime option in treePL; smoothing value = 1, determined by random subsample and replicate cross-validation method (RSRCV), was used for both dating analysis and bootstrapping (see below). The calibrations (shown in, Supplemental Fig.7) are as follows: 1) Mygalomorphae: Due to the lack of relevant fossil calibration points for the Mygalomorphae – Mesothelae split, we used the information available from phylogenomic studies based on extensive outgroup fossil calibrations (Garrison et al. 2016; Fernández et al. 2018) in order to constrain the age of the root node. Namely, we used the 95% confidence intervals estimated for this split as the minimum (287 Ma) and maximum bounds (398 Ma); 2) Avicularioidea: The age of the oldest mygalomorph fossil *Rosamygale grauvogely* (Selden and Gall 1992) from Gres-a-Voltzia Formation, Vosges, France dated to Middle Triassic (Anisian) 242 Million years ago (Ma) was assigned as the minimum bound for the Avicularioidea – Atypoidea split. The fossil has been tentatively assigned to the family “Hexathelidae” (*sensu* Raven) but given the polyphyly of the family (Hedin et al. 2018), we rather interpret *Rosamygale* as an Avicularioidea crown-group. The maximum bound for the Avicularioidea – Atypoidea split was assigned the age of 323 Ma, corresponding to the oldest age of the Bashkirian stage (Pennsylvanian, Carboniferous) that yielded the oldest spider fossil *Arthrolycosa* sp. putatively assigned as Mesothelae stem-group (Selden et al. 2014; Garwood et al. 2016). This age also overlaps with the oldest boundary of the 95% confidence intervals for the Avicularioidea – Atypoidea split estimated by recent phylogenomic studies (Garrison et al. 2016; Fernández et al. 2018); 3) “Nemesioidina” clade: The age of *Cretamygale chasei* 125 Ma (Selden 2002) from Cretaceous (Barremian, Hughes and McDougall 1990) currently classified as Nemesiidae was conservatively assigned as the minimum bound for the split between Bemmeridae/Theraphosidae and “Nemesioidina” clades. The maximum bound was assigned 242 Ma, corresponding to the age of *Rosamygale* fossil. 4) Atypoidea: Atypidae – Antrodiaetidae split was assigned minimum bound of 100 Ma representing *Ambiortiphagus ponomarenkoi* (Eskov and Zonshtein 1990) from the Lower Cretaceous (Albian) proceeding from Dund-Ula Mountain Ridge in Bayan-Khongor Province, Central Mongolia. The age of *Rosamygale* (242 Ma) was assigned as a maximum bound for the split; and 5) *Ummidia* fossils (44 Ma) from Baltic amber (Wunderlich 2011) were used to calibrate the minimum bound for the split between the North American and European species of *Ummidia*. Maximum bound of 90 Ma corresponds to the oldest boundary of the 95% confidence interval estimated for this particular split (Opatova et al. 2013). The confidence intervals for node ages were assessed by dating 100 ML bootstrap phylograms inferred in PAUP* (Swofford 2003). The trees were dated in TreePL using the same settings as described above; TreeAnnotator v. 2.4.5 (Bouckaert et al. 2018) was used for calculating the age statistics for the nodes.

Biogeographic analyses were conducted in Reconstruct Ancestral State in Phylogenies (RASP v 3.1, Yu et al. 2015) with the dispersal-extinction-cladogenesis (DEC) model (Ree and Smith 2008) implemented in C++ version of Lagrange (Smith 2010). Analyses were run with no dispersal constraints, assigning equal probabilities to dispersal events among all defined areas, and allowing two unit areas in the ancestral distributions. The terminal taxa represented in the tree were assigned to six distribution ranges: (A) North America; (B) Europe; (C) Asia; (D) South America; (E) Africa; (F) Australia plus New Zealand (Fig. 5b). The consensus tree obtained in ExaBayes was used as an input for the analyses.

For ancestral state reconstruction analyses of foraging construct, each terminal taxon was scored using six character states: (0) burrow with trapdoor(s); (1) brush sheet/funnel web; (2) burrow with collar door; (3) purse web; (4) open burrow; (5) turret. Character codings were scored from personal observations and/or the primary literature as documented in Supplemental Table S3. Taxa that forage as cursorial hunters were scored as unknown or missing (Coddington et al. 2018); polymorphic character states were assigned to terminals for instances where species build multiple/combination of constructs (e.g., some *Cyrtauchenius* terminals build turrets as juveniles but cover those turrets with trapdoors as adults, Opatova *persn. obsv*.), or closely related species build more than one construct e.g., both open burrows and trapdoors (some bemmerids and pycnothelids) and both turrets and collars (e.g., some *Antrodiaetus*). Ancestral character state reconstructions were conducted using the R package *corHMM* (Beaulieu et al. 2013) on an ultrametric scaled tree with node.states=marginal; *corHMM* was preferred over other methods given its capacity to accommodate missing and polymorphic characters. The R package *ape* was used to scale the preferred tree topology (‘chronopl’) with an assigned lambda value of 0.1. Character optimizations using equal (ER), symmetric (SYR), and all rates different (ARD) models were explored for these categorical data using ‘rayDISC’ (*corHMM*). The preferred model was chosen by comparing AICc values calculated using *corHMM*.

All relevant data, DNA and amino acid sequence alignments, phylogenetic trees, and miscellaneous data matrices, were deposited in the Dryad Data Repository (doi:XXXXXXXXXXX) on XX XX 2019.

## Results

### Concatenated analyses, all taxa

The complete dataset “All_taxa” supermatrix comprises 472 loci (93,410 nucleotides) for 113 terminals (Supplemental Table S1), whereas the reduced “No_Paratropis” dataset comprises 111 terminals. The proportion of missing data in the datasets is 14.7% and 14.5%, respectively. The combined “DNAAA_matrix” includes 30 terminals, with 14 individuals represented by both genomic (472 AHE loci) and transcriptomic data (1519 loci proceeding from Garrison et al. (2016)). The total length of the “DNAAA_matrix” is 305,860 characters; gaps and missing data represent 54.5% of the dataset. For details see Supplemental Table S2.

Both ML (-ln 1728131.7475) and BI analyses performed on the “All_taxa” dataset yielded similar tree topologies (Fig. 3; Supplemental Fig. 1, 2, 3). Overall, the resulting trees are highly supported (bootstrap > 90, posterior probability (PP) > 0.95, SH-aLRT > 95), with the exception of a few deeper nodes. The ML and BI analyses both recover two main clades Atypoidea and Avicularioidea with strong support. The relationships within Atypoidea are fully supported; the family Atypidae is recovered as a sister to Antrodiaetidae inclusive of *Hexura* (Mecicobothriidae).

**Figure 3.**
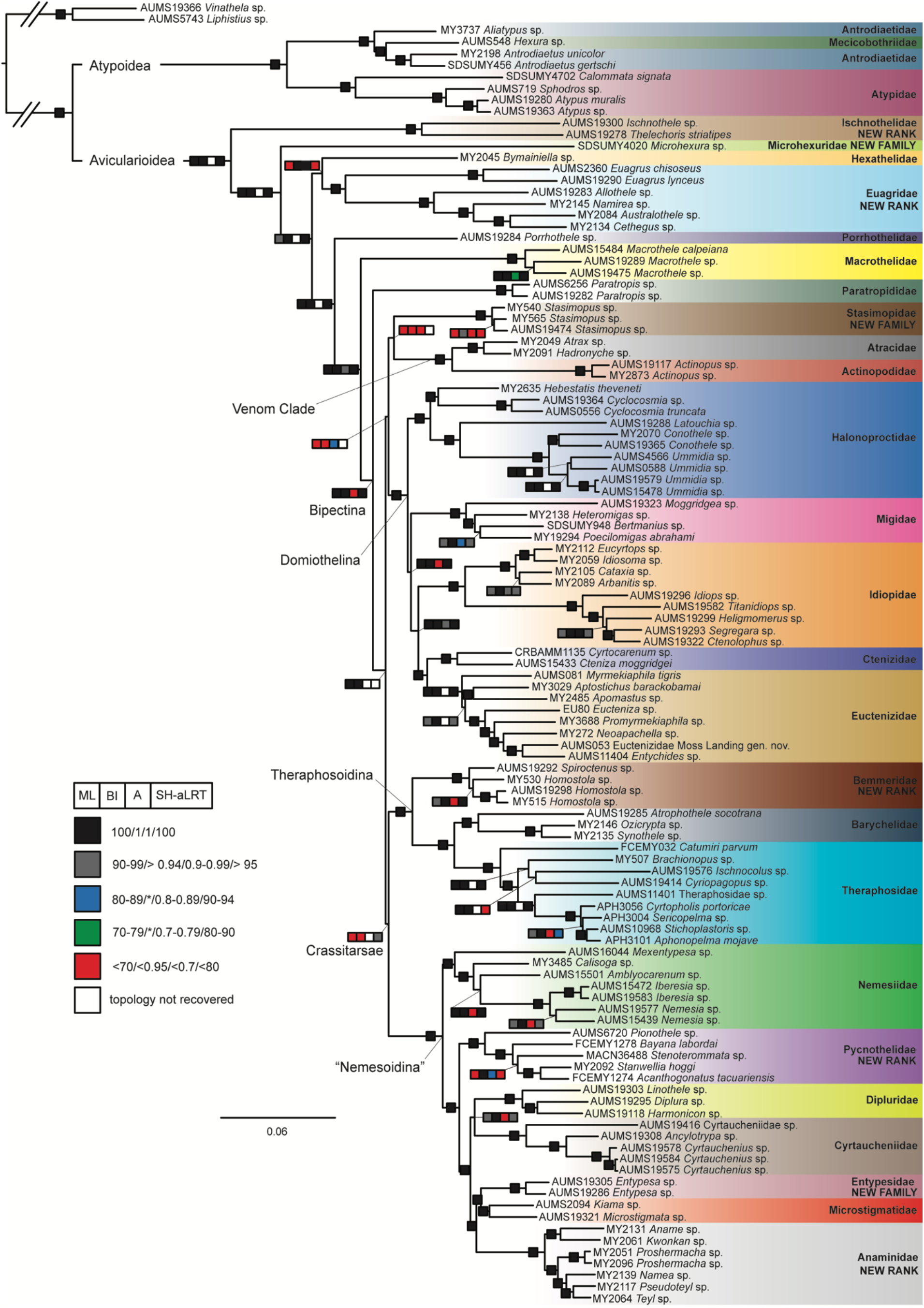
Phylogenetic tree summarizing results from concatenated and species tree approaches. Topology obtained in the Maximum Likelihood analyses. Boxes on nodes denote support values obtained in each approach (left to right): RAxML bootstrap support, ExaBayes Bayesian posterior probabilities (PP), ASTRAL support values, IQ-Tree SH-aLRT support values. Color coding of boxes corresponds to distinct support level categories depicted in bottom left corner. One filled box indicates full support in all analyses; white box=topology not recovered in species tree analysis.

Four mygalomorph families belonging to Avicularioidea are recovered as para- or polyphyletic (namely Dipluridae, Ctenizidae, Cyrtaucheniidae and Nemesiidae), prompting extensive higher-level changes to mygalomorph classification (see below). Several lineages formerly belonging to the families Dipluridae and Hexathelidae (see Hedin et al. (2018)) successively branch off near the root node of Avicularioidea as follows: Ischnothelidae (NEW RANK), Microhexuridae (NEW FAMILY), Hexathelidae, Euagridae (NEW RANK), Porrhothelidae and Macrothelidae. The Bipectina clade (*sensu* Goloboff (1993)) inclusive of the family Atracidae is recovered in our analysis. The family Paratropididae is recovered as sister to all the remaining Bipectina. The remaining taxa are placed into a clade comprising Stasimopidae and the “Venom Clade” (Atracidae + Actinopodidae) and two additional clades corresponding to Domiothelina and Crassitarsae (*sensu* Bond et al. (2012), but excluding *Stasimopus*). The monogeneric family Stasimopidae (NEW FAMILY, formerly Ctenizidae), is placed as sister to “Venom Clade”, albeit with low support. The ML analysis performed in IQ-TREE resulted in different placement of Stasimopidae and the “Venom Clade”. The “Venom Clade” is supported as sister to all Bipectina minus Paratropididae, whereas the family Stasimopidae is recovered with high SH-aLRT support as sister to Domiothelina. Domiothelina form a monophyletic group with well resolved relationships in both ML and BI analyses. The family Halonoproctidae is recovered as sister to all remaining lineages of Domiothelina. The family Migidae is recovered as sister to Idiopidae, Ctenizidae (now comprising *Cteniza* and *Cyrtocarenum*) and Euctenizidae. Crassitarsae (*sensu* Bond et al. (2012)), recovered with high SH-aLRT support in IQ-TREE ML inference, is subdivided into two clades supported in all analyses. The first one comprises the family Bemmeridae (NEW RANK, formerly Cyrtaucheniidae), recovered as sister to the Barychelidae and Theraphosidae. The second clade, “Nemesioidina”, comprises the Nemesiidae plus three lineages formerly belonging to the Nemesiidae (Pycnothelidae (NEW RANK), Entypesidae (NEW FAMILY) and Anamidae (NEW FAMILY)), as well as Dipluridae, Cyrtaucheniidae and Microstigmatidae. Nemesiidae is recovered as sister to all the remaining families; Pycnothelidae is recovered as sister to the Dipluridae/Cyrtaucheniidae clade and the Entypesidae /Microstigmatidae/Anamidae clade.

### No_Paratropis analysis

Maximum likelihood analyses of the “No_Paratropis” dataset recovered a moderately different Bipectina topology contra the analyses on the full dataset (Supplemental Fig. 4; ML best tree –ln = 1702659.2948). The discordance, however, mostly affects nodes unsupported in both analyses. In the “No_Paratropis” analyses the clade comprising Bemmeridae + Barychelidae + Theraphosidae is recovered with high support as sister to all the remaining Bipectina families. The Stasimopidae + “Venom Clade” receive moderate bootstrap support, but its sister relationship to the Domiothelina clade remains unsupported, albeit with increased support when compared to the analyses of the full dataset.

### DNAAA analysis

Similar to the “No_Paratropis” dataset, analyses of “DNAAA” (ML best tree –ln = 2543315.0197) resulted in different topological arrangements within the Bipectina clade (Fig. 4). The Domiothelina clade is recovered as sister to all the remaining diversity. Stasimopidae + “Venom Clade” (moderately supported in this analysis) is recovered with low support as sister to Crassitarsae. The Crassitarsae clade is highly supported and inclusive of *Paratropis*.

**Figure 4.**
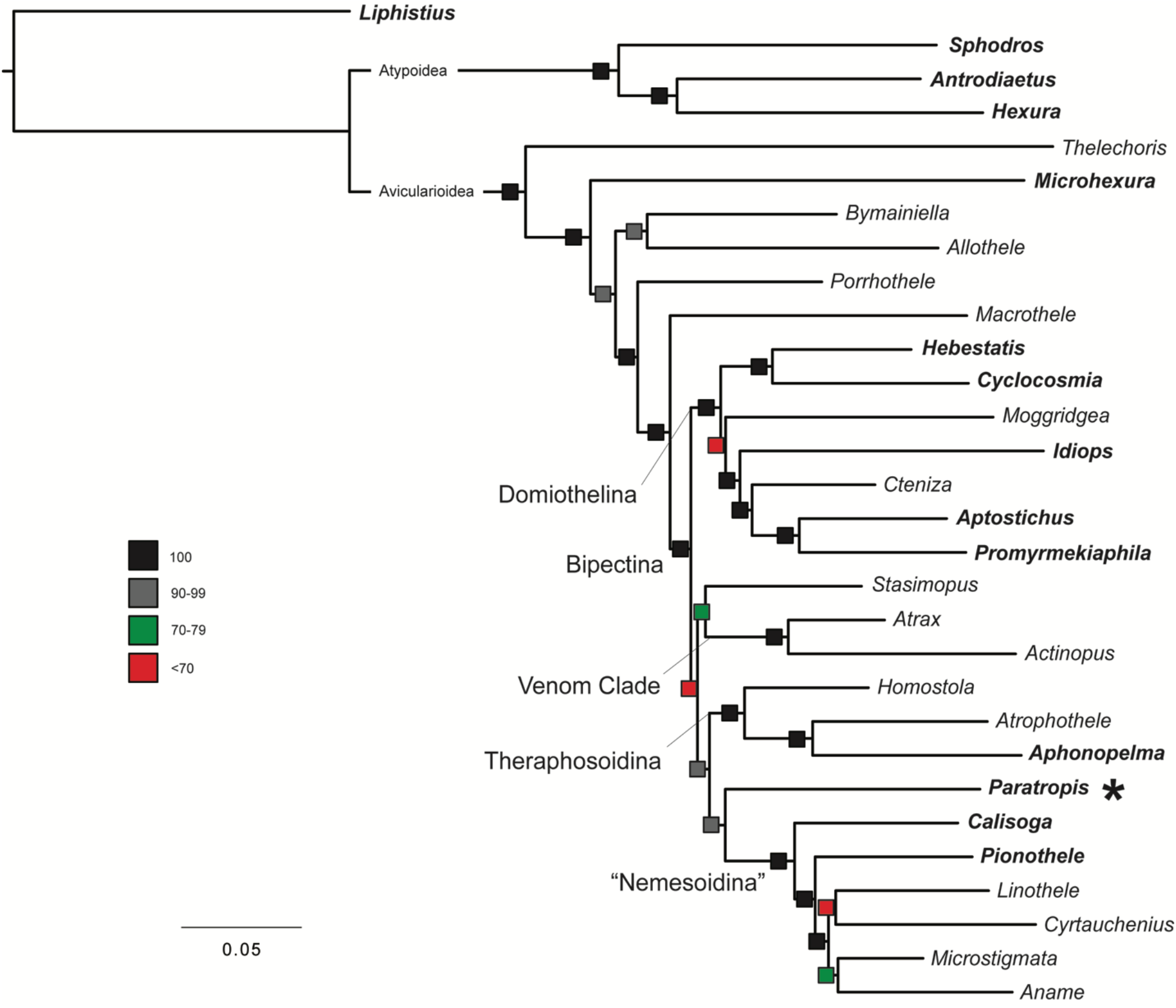
Phylogenetic tree inferred from combined AHE loci and transcriptomic data (“DNAAA” matrix). Tree topology obtained in maximum likelihood (ML) analyses conducted in RAxML. Asterisk marks placement of *Paratropis* within Crassitarsae. Boxes near the branches denote bootstrap support values; colors correspond to distinct support level categories depicted on left.

### Topology tests

The AU topology test (Table 2) rejected the monophyly of all alternative arrangements (i.e., alternatives using existing compositions of genera) of the families Ctenizidae (*sensu* Raven (1985)), Cyrtaucheniidae (*sensu* Bond et al. (2012)), Dipluridae, Hexathelidae (*sensu* Raven (1985)) and Nemesiidae in favor of the results of the unconstrained search.

### Species tree analysis

Coalescence-based analysis produced a species tree from the individual 472 input gene trees obtained in RAxML. The resulting quartet-based supertree estimated in ASTRAL (Supplemental Fig. 5) comprises 1,228,051,216 induced quartet trees from the input gene trees, representing 65.2% of all quartets present in the species tree. ASTRAL yielded a different topology than the concatenated approach (Supplemental Fig. 6). The analyses recover both Atypoidea and Avicularioidea clades. Ischnothelidae (NEW RANK), Microhexuridae (NEW FAMILY), Hexathelidae, Euagridae (NEW RANK) and Porrhothelidae, a grade of lineages branching at the base of the Avicularioidea clade in the concatenated analyses, form a highly supported clade in the species tree approach. The topology of the Bipectina clade (unsupported in ASTRAL) is also somewhat differed from the concatenated analyses. Bemmeridae/Barychelidae/Theraphosidae clade is recovered as sister to moderately supported Domiothelina + Stasimopidae/“Venom Clade”, rendering the Crassitarsae paraphyletic.

### Molecular dating

The resulting dated tree is depicted in Fig. 5; Supplemental Figures 7, 8. The age of the root (398 Ma) corresponds to the maximum age constraints assigned to this split. The split between Atypoidea and Avicularioidea is dated at ∼ 323 Ma. The early branching events within Avicularioidea dated between 254 Ma [95% confidence interval (CI): 259 – 247] and 174 Ma (179 – 162) (Ischnothelidae through Macrothelidae divergences). The first diversification of Bipectina is dated to 166 Ma (171 – 155) (divergence of Paratropididae) and continues at 158 Ma (163 – 147), with diversification into Domiothelina + Stasimopidae/“Venom Clade” and Crassitarsae. The divergence time estimates for individual families are reported in Table 3.

**Figure 5.**
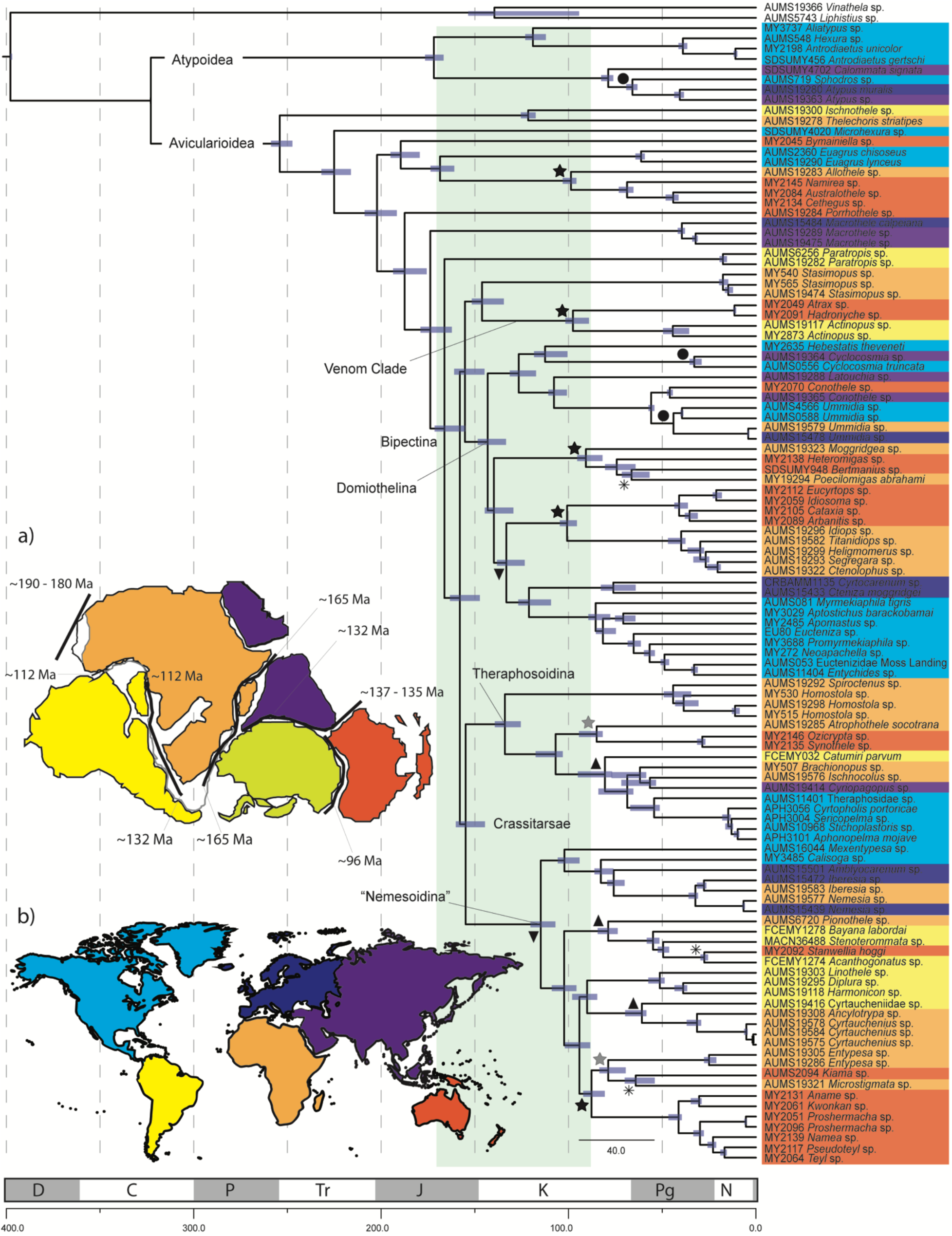
Divergence time estimates inferred by treePL on a topology obtained in RAxML. The x-axis is time in million years. Geologic time scale abbreviations: N: Neogene, Pg: Paleogene, K: Cretaceous, J: Jurassic, Tr: Triassic, P: Permian, C: Carboniferous, D: Devonian. Light green block marks time frame of Gondwana breakup. Symbols on the tree nodes denote vicariant or dispersal events hypothesized for the divergences (left to right): inverted triangle=Pangea breakup, star=East-West Gondwana breakup, triangle=West Gondwana breakup, dot=Laurasia breakup, asterisk=long distance dispersal. Left corner: Map showing the position of the continents (a) prior to Gondwana breakup (adapted from (Will and Frimmel 2018), thick black lines mark the zones of continental drifting, times indicate the initiation of drifting in each zone; (b) present day. Terminal tree taxa are color coded according to geographic areas of their sampling locations depicted in the map as follows: yellow=South America, orange=Africa, purple=Asia, red=Australia, light blue=North America, dark blue=Europe, lime green=Antarctica (no taxa assigned).

**Table 3.**
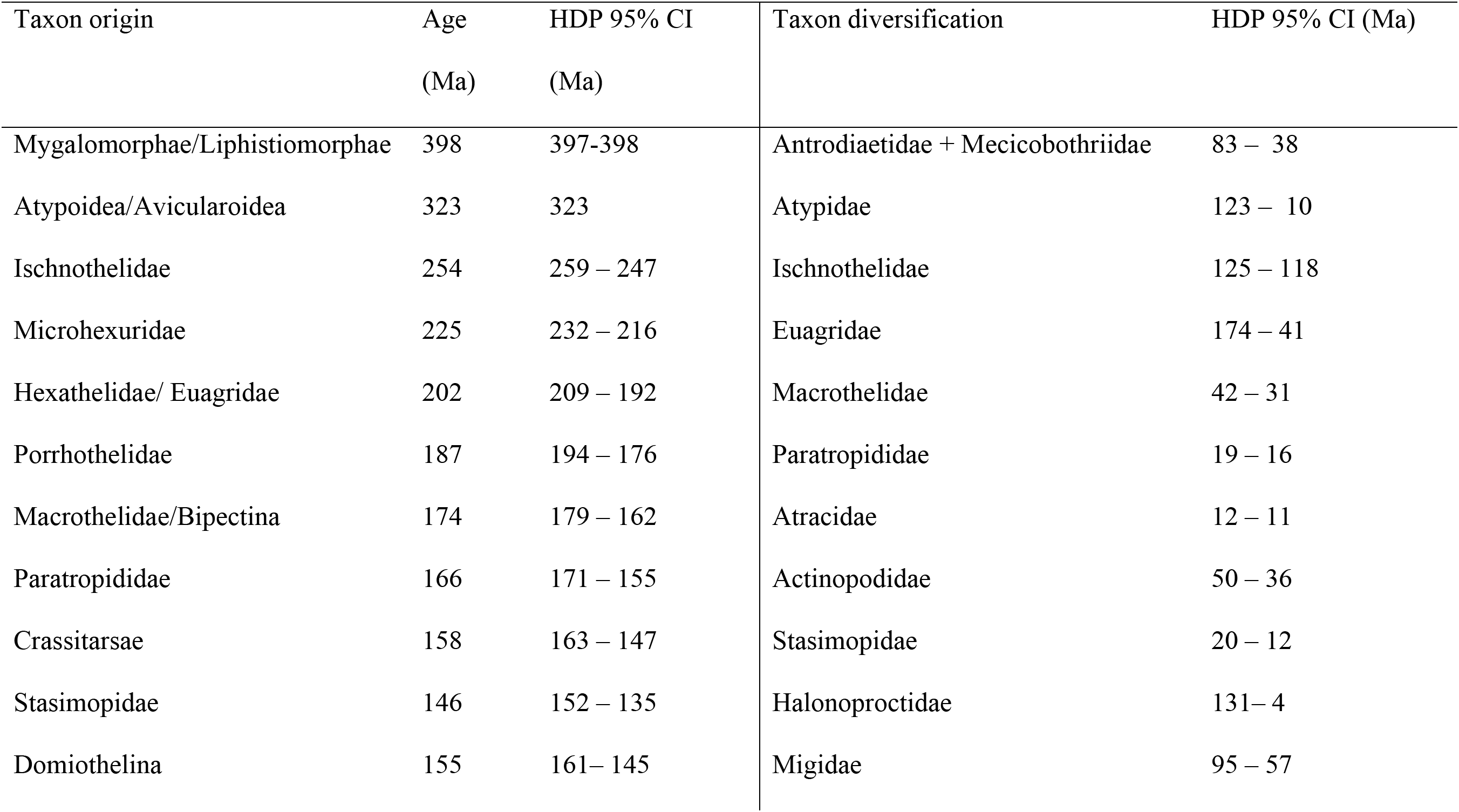
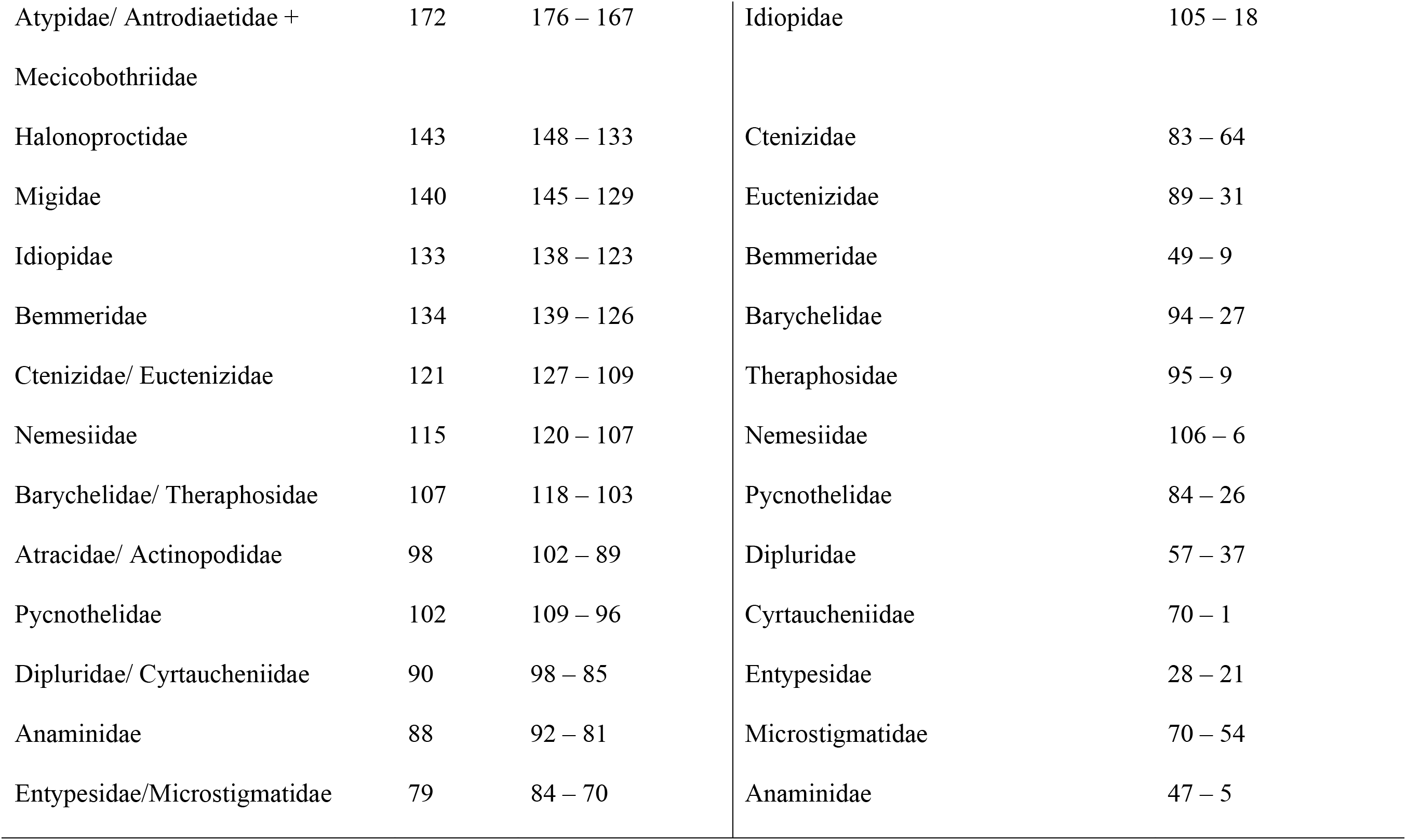
Divergence time estimates obtained by treePL algorithm (Smith and O’Meara 2012) representing taxa origins and the respective timeframe of their diversification inferred for tree topology depicted in Fig. 5. Confidence intervals (CI) for the highest posterior density (HDP) were inferred from 100 bootstrap replicates.

### Historical biogeography

The results of DEC analyses are summarized in Fig. 6. The ancestral distributional area of the mygalomorph ancestor and the early branching Avicularioidea lineages remain unresolved. The distribution of the Atypoidea ancestor is inferred to have originated in the Northern hemisphere continents (North America and Asia); Bipectina likely originated in Africa. The ancestral ranges of several Avicularioidea families are inferred to areas spanning more than one continent, for example Africa and Australia (Migidae, Idiopidae, Barychelidae, Microstigmatidae); Africa and South America (Cyrtaucheniidae, Pycnothelidae). On the other hand, some Avicularioidea families experienced *in situ* diversification within a single area. Stasimopidae, Bemmeridae, and Entypesidae likely evolved in Africa, Paratropididae and Dipluridae in South America, Anamidae in Australia, Euctenizidae in North America and Ctenizidae in Europe. DEC analyses also infers a large number of dispersal and vicariance events across the mygalomorph phylogeny, however, most of the events receive low probabilities (<0.7).

**Figure 6.**
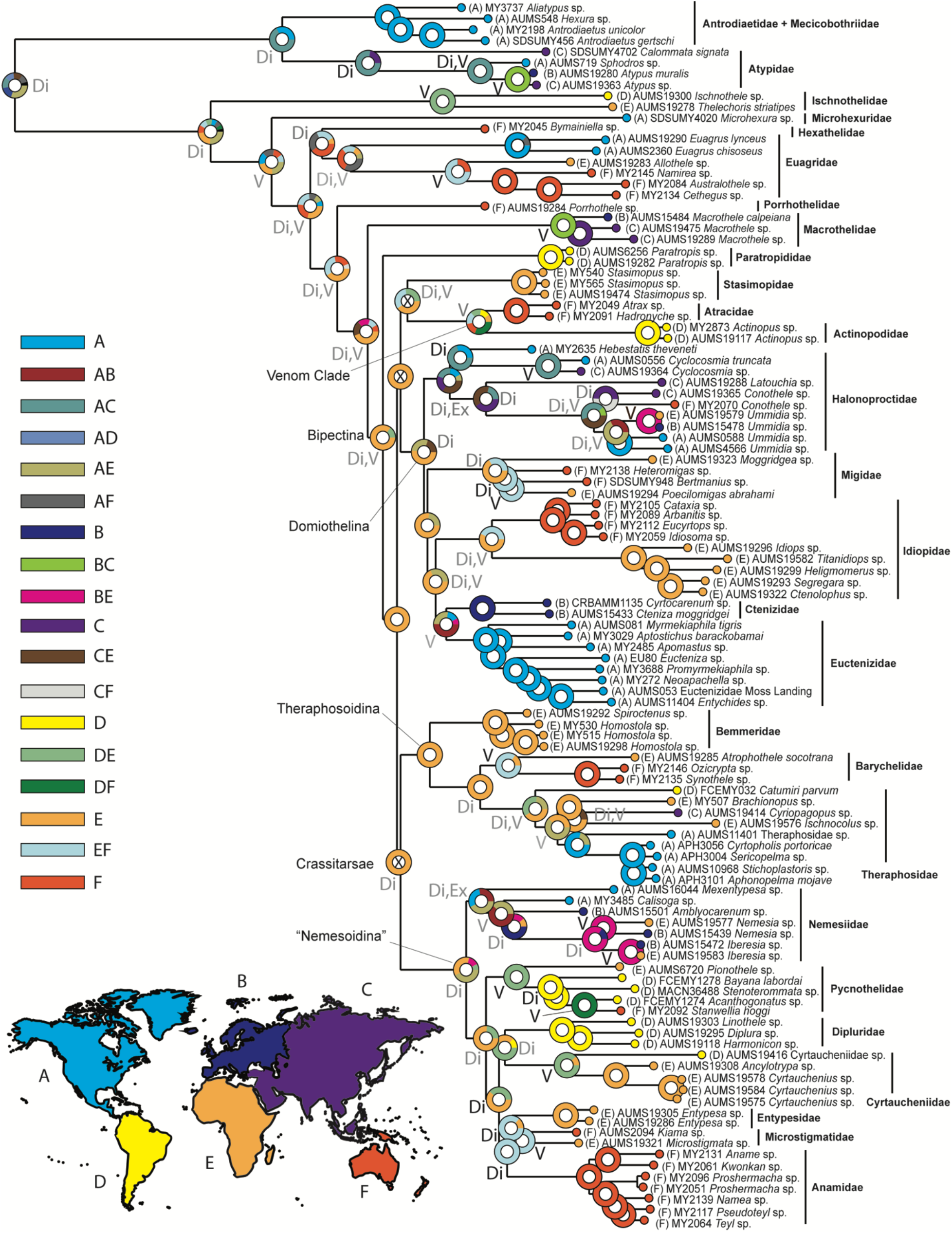
Ancestral areas of distribution inferred with DEC analyses implemented in RASP. Tree topology obtained in ExaBayes. Terminal tree taxa are color coded according to the biogeographic regions of their sampling locations depicted in the map (bottom left) as follows: A=North America, B=Europe, C=Asia, D=South America, E=Africa, F=Australia. Color coding of the inferred ancestral distributions corresponds to the assigned biogeographic regions, or combination of thereof depicted in the combined area legend on the left. Biogeographic events marked on the nodes: Di=dispersal, V=vicariance, Ex=extinction; black letters: events inferred with probability > 0.7.

### Evolution of foraging construct

We scored foraging construct for 109 taxa. Taxa that are cursorial hunters (three terminals), *Paratropis* and *Microstigmata*, were scored as missing/inapplicable. The foraging construct of nemesiid *Mexentypesa* is unknown to us. Because bemmerids and pycnothelids construct both trapdoors and open burrows they were scored as polymorphic. Based on AICc values the equal rates model (ER) produced the preferred reconstruction (AICc = 168.3399). Although the ER model was preferred, ancestral state reconstructions for the other models were generally very similar.

The ancestral state reconstruction of mygalomorph foraging construct, summarized in Fig. 7, shows a rather simple and concise pattern. Towards the root of the phylogeny there appears to be an equal probability of an ancestor having either a funnel-and-sheet web or a burrow with a trapdoor. Atypoids likely had a most recent common ancestor (MRCA) with a funnel-and-sheet web, modified as a purse web in atypids or as a collar, turret, or trapdoor in antrodiaetids. As discussed below, atypoids demonstrate the full scope of variability in burrow construct modifications that can evolve from a sheet web ancestral form. Avicularioidea have a MRCA that optimizes as having a sheet web. The MRCA for the Bipectina clade likely had either a trapdoor or funnel-and-sheet web (equivocal). A burrow with a trapdoor more definitively optimizes as the ancestral foraging construct for Crassitarsae with subsequent losses of the trapdoor covering (optimized as the open burrow state) four times in nemesiids, pycnothelids, theraphosids, and the euctenizid genus *Apomastus*; some pycnothelid and anamid taxa independently lose the trapdoor and instead forage from burrows with distinct turret or collar door constructs. Funnel-and-sheet webs are regained twice, once in atracids and again in diplurids.

**Figure 7.**
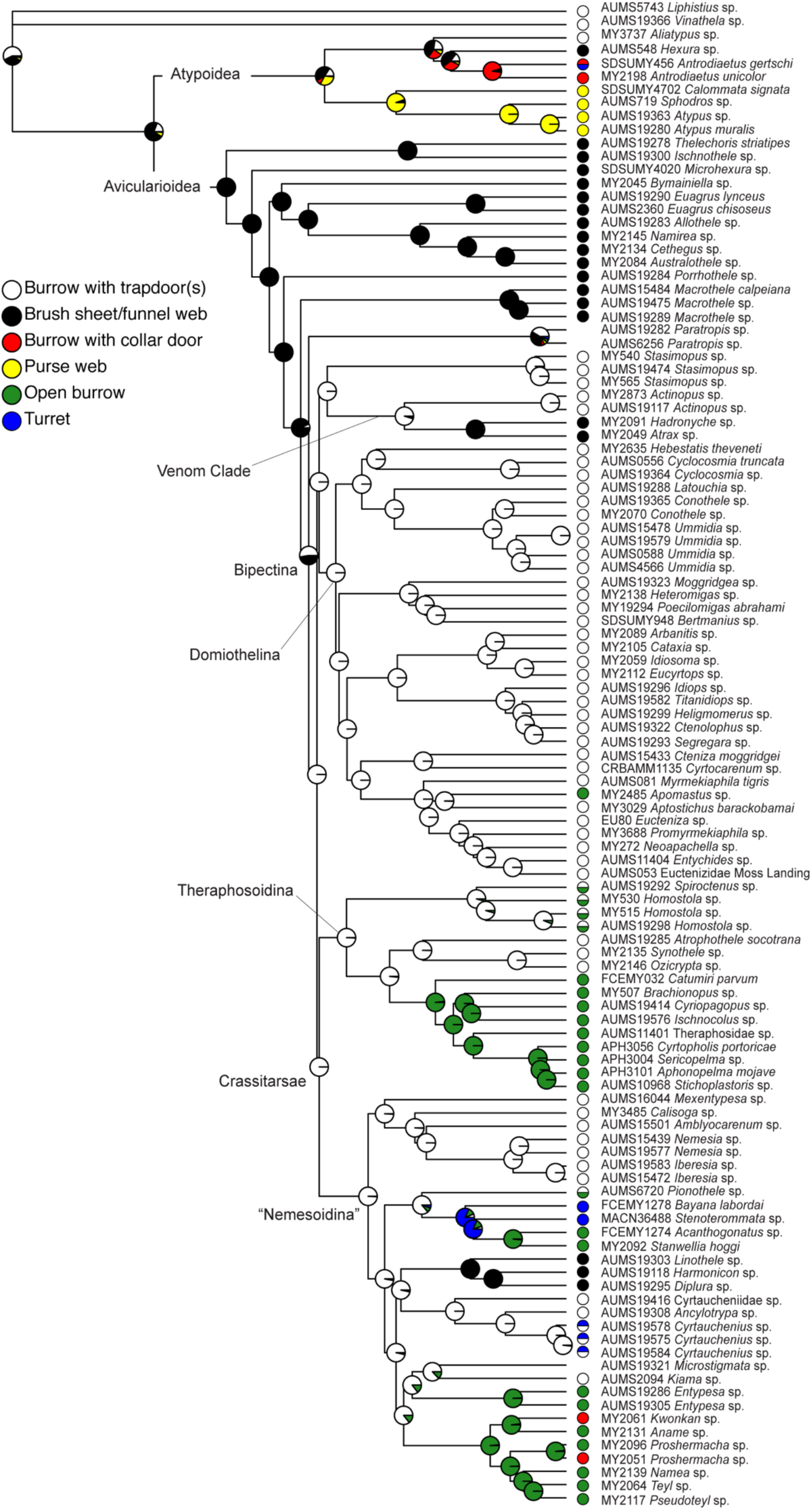
Ancestral state reconstruction of mygalomorph foraging constructs. Preferred ancestral state reconstruction of foraging type using an equal rates model in the R-package *corHMM* with cursorial hunting taxa treated as inapplicable (?); tree was modified as ultrametric; AICc = 173.9083. Terminal tree taxa are color coded according the type of spinning structure depicted in the legend panel of the left.

## Discussion

In light of the results reported herein and other past molecular based studies of mygalomorph phylogeny, two points seem clear. First, morphological data *alone* are unsuitable for resolving the evolutionary history of mygalomorphs. Second, if our aim is to have a classification scheme that reflects evolutionary history, other characters are needed. The results we discuss below allow us to assess the status of nearly all mygalomorph families and resolve several long-standing questions in the group’s phylogeny and higher-level classification. As a consequence, we propose extensive taxonomic changes that better reflect the origins and evolutionary relationships of the taxa. We discuss in detail 1) the temporal and biogeographical context of mygalomorph family-level diversification; 2) revisit longstanding questions regarding silk use in non-araneomorph spiders; and 3) provide a detailed account regarding the taxonomic implications of our newly formed phylogenetic hypothesis (Fig. 8). The Appendix to this paper (Bond et al. 2019) outlines the formal taxonomic changes proposed herein and discusses in more detail the systematics of each mygalomorph family.

**Figure 8.**
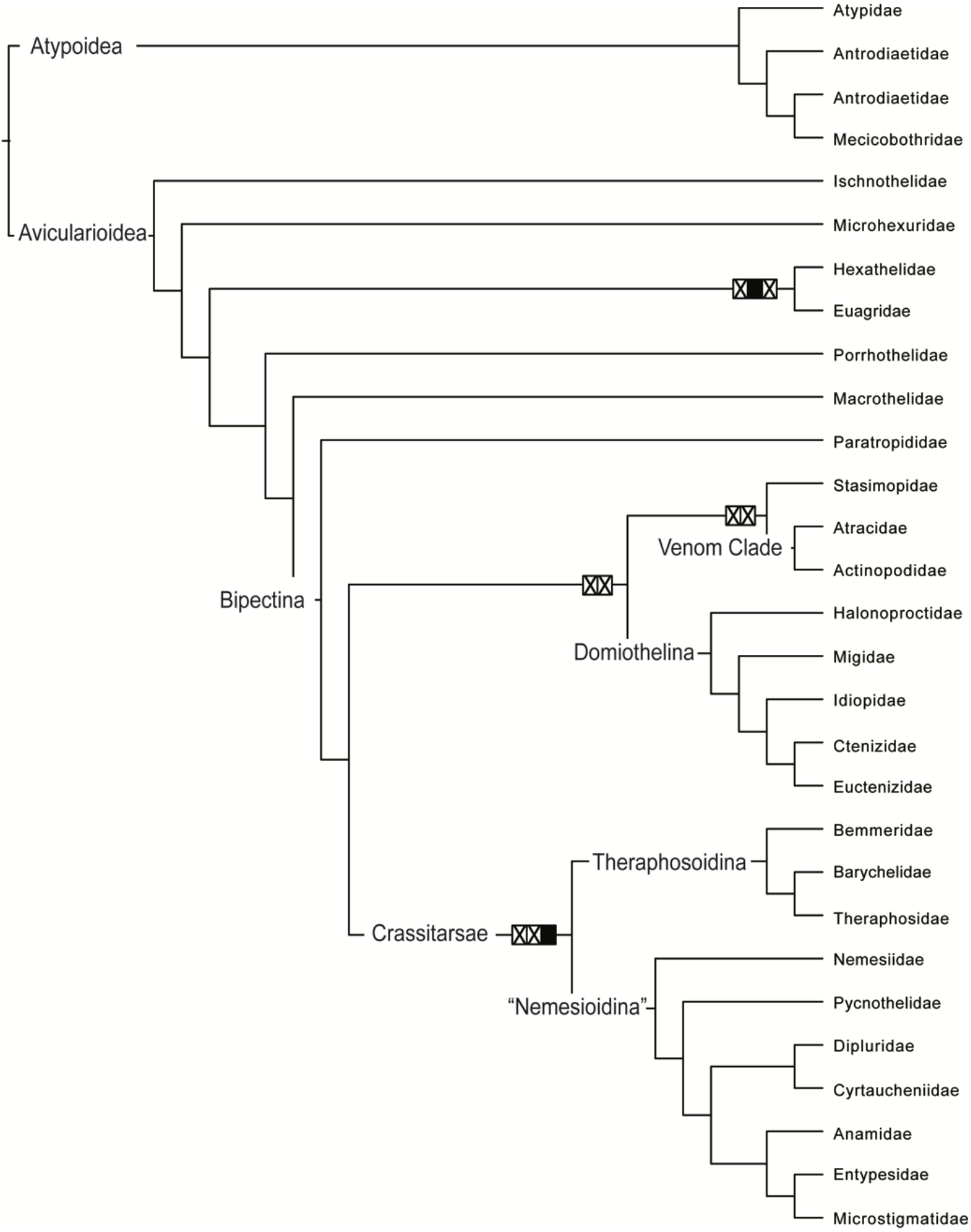
Cladogram summarizing the phylogenetic relationships of mygalomorph families recovered in this paper. The topology corresponds to the arrangement obtained in the concatenated analyses (RAxML, ExaBayes, IQ-TREE). The crossed boxes on the branches represent unsupported relationships: left ML bootstrap support < 70%, right Bayesian inference Posterior Probability< 0.95, SH-aLRT < 95). Filled box=node supported only in Bayesian inference, otherwise relationships supported in both analyses.

### Diversification and Biogeography of Mygalomorph Spiders

Mygalomorph spiders with nearly a cosmopolitan distribution, highly sedentary nature (Raven 1985; Bond et al. 2001) and evolutionary history dating back to Carboniferous (Ayoub et al. 2007; Starrett et al. 2013; Garrison et al. 2016), represent the type of group that has captivated the attention of biogeographers for decades (Cracraft 1988; Wiley 1988; Morrone and Crisci 1995). By implementing divergence time estimation methods, many distribution patterns observed in mygalomorphs could be linked to specific geologic events such as continental breakups (Opatova et al. 2013; Opatova and Arnedo 2014a), Mediterranean tectonics (Opatova et al. 2016; Mora et al. 2017) and opening of the Okinawa Trough (Su et al. 2016). However, recent findings suggest that dispersal via rafting or could be a plausible mechanism for overcoming both short (Hedin et al. 2013; Opatova and Arnedo 2014b) and long distances (Harrison et al. 2017).

The results of our biogeographic analyses (Fig. 6) place the ancestral area of distribution of Atypoidea within the Northern hemisphere (across North America and Asia), whereas the ancestor of Porrhothelidae, Macrothelidae and Bipectina most likely evolved in the Southern hemisphere (across Africa and Australia). In our dating analyses, we recovered two repeated patterns of divergences occurring among taxa with present day distributions confined mostly to the Southern hemisphere that in dispersal-limited invertebrate groups have been repeatedly linked to vicariant events related to the different phases of Gondwana breakup (Wood et al. 2012; Frazão et al. 2015; Kim and Farrell 2015; Xu et al. 2015; Andújar et al. 2016; Toussaint et al. 2017).

The initial phase of Gondwana breakup (Fig 5), i.e., the formation of East and West Gondwana, began approximately 165 Ma with South America and Africa drifting away from India, Madagascar, Antarctica and Australia. The pattern concordant with the East – West Gondwana breakup observed in our results involved splits within the families Euagridae, Migidae, Idiopidae (putatively also Barychelidae), and divergence between Atracidae – Actinopodidae and Anamidae – Entypesidae /Microstigmatidae clade (marked with stars on the chronogram in Fig. 5) was dated between 101 – 88 Ma. The second pattern, concordant with West Gondwana fragmentation into South America and Africa occurring between 132 – 112 Ma (Will and Frimmel 2018) involved divergences within Theraphosidae, Pycnothelidae and Cyrtaucheniidae (marked as triangles) was dated to 81 – 61 Ma. The divergence times estimated in our analyses were considerably younger than the ages estimated for other sedentary taxa with former Gondwanan distribution, such as velvet worms or bothriurid scorpions (Murienne et al. 2014; Sharma et al. 2018). Our inferred divergences also postdate the onset of continental drifting in both instances by approximately 50 My, however, the initial phases of different Gondwana breakup stages were often characterized by slow tectonic movement, or even quiescence (Boger 2011; Will and Frimmel 2018; Young et al. 2018). In case of East and West Gondwana, the drifting was occurring at an extremely slow rate for at least the first ten million years (Young et al. 2018). A connection between South America and Australia also likely persisted via Antarctica until late Eocene, approximately 35 Ma (Sanmartín and Ronquist 2004), which links the divergence time estimates between Neotropical and Australasian groups of Darwin’s stag beetles dated to 47 and 58 Ma, respectively, to Gondwana breakup (Kim and Farrell 2015).

Conversely, the divergence time estimates inferred in our analyses could also be affected by the limited fossil record for mygalomorphs, and our conservative approach towards node calibration. For example, the *Cretamygale chasei* fossil, formerly interpreted as Nemesiidae, was used to set a minimum bound for the whole “Nemesioidina” clade (see methods), rather than arbitrarily assigning the fossil to the family Nemesiidae (*sensu* this study). Despite the apparent advantages in the ability of handling genomic data, the dating analyses used in this study (Smith and O’Meara 2012), does not allow to assign probability distribution priors to the calibration points or infer confidence intervals for the divergences. Confidence intervals are often essential for connecting the age estimates and specific geological events and while assessing them via bootstrapping potentially informs our analyses, this approach likely does not capture the full span of variation as accurately as when node calibration priors are taken into account.

Alternatively, the observed patterns could be explained by long-distance dispersal events as in case of other poorly dispersing groups such as euedaphic beetles (Andújar et al. 2016) or short-tailed whip-scorpion order Schizomida (Clouse et al. 2017). Although evidence suggests that mygalomorph spiders are capable of trans-oceanic dispersal (Harrison et al. 2017), repeated divergence patterns imply multiple long-distance dispersal events occurring independently among unrelated taxa during the same timeframe. Continental-level vicariance thus provides a more plausible explanation for the observed pattern.

The East – West Gondwana vicariant origin of the taxa is also supported by the results of the ancestral range reconstruction analyses. The inferred ranges for all the taxon groups displaying a distribution pattern concordant with the vicariance (Euagridae, Atracidae and Actinopodidae, Migidae, Idiopidae, Barychelidae, Entypesidae and Anamidae) comprised former Gondwana continents, although the ancestral range inferred for Idiopidae, Barychelidae and Migidae (Africa and Australia) could potentially represent an artifact stemming from a geographical bias in our sampling (i.e. lacking their South American representatives in our analyses). Vicariance was also inferred with high probability in the divergences between *Allothele* and Australian euagrids, *Atrophothele* and remaining barychelids. The West Gondwana vicariant split between Africa and South America can likely be attributed to the structure within Pycnothelidae (*Pionothele* – remaining pycnothelids) and South American and African Cyrtaucheniids which also optimizes as an ancestral distribution spanning across both continents in both groups.

Another repeated pattern observed across our tree includes taxa with currently disjunct amphi-Atlantic distributions (marked as dots on the chronogram). Such patterns of divergence were observed within *Ummidia* and *Cyclocosmia* (Halonoproctidae) and between *Sphodros* and *Atypus* (Atypidae), and were dated to 44 Ma, 33 Ma (35 – 29) and 39 Ma (41 – 37), respectively. This pattern has previously been connected with the breakup of Laurasia (Opatova et al. 2013; Godwin et al. 2018) dating to the end of Paleocene, approximately 60 – 55 Ma (Sanmartín et al. 2001) and it is supported by the ancestral distributions of the taxa projected to the Northern hemisphere continents in our analyses. Our divergences postdate both the geological event and previous dating results (Opatova et al. 2013). The discrepancy between these time estimates and the Laurasian breakup could be explained by either an artefact of the analyses (discussed above), or alternatively by a persistent connection of North America and Europe via a North American land bridge. This connection existed to some extent until 25 Ma and is hypothesized to have played an important role in plant dispersal (Tiffney and Manchester 2001; Milne 2006), suggesting also a suitable dispersal route for ballooning spiders like *Atypus*, *Sphodros*, *Ummidia* (Opatova et al. 2013). The current distribution of *Cyclocosmia* (East Asia and South East of North America), represents a well-known biogeographic pattern linked to continental vicariance (Boufford and Spongberg 1983; Bolotov et al. 2016), which is further supported by the inferred vicariant origin of the genus. Alternatively, similar distribution patterns could have resulted from dispersal events between Palearctic and Nearctic regions via Beringia (Li et al. 2015; Bolotov et al. 2016). However, the timing of the dispersal in other organisms postdates the results of our analyses (Li et al. 2015; Maguilla et al. 2018).

Older divergences recovered in our analyses are more challenging to explain within a biogeographic context. If we follow the same line of reasoning as explained above, and assume that the actual ages of the splits are older than indicated by the results, the splits between the Idiopidae and Ctenizidae/Euctenizidae clade, and Nemesiidae and the remaining “Nemesioidina” diversity (marked as inverted triangles), could correspond to the disintegration of the supercontinent Pangea occurring in Lower Jurassic (190 – 180 Ma) (Beutel et al. 2005; Will and Frimmel 2018). Divergences proceeding the breakup of Pangea could be potentially linked to paleoclimatic events. Since its formation around 520 – 510 Ma (Veevers 2004; Domeier and Torsvik 2014; Will and Frimmel 2018), large parts of Gondwana experienced repeated glaciations in the Silurian, Devonian, late Carboniferous and early Triassic. Additional glaciations were caused by the orogenic processes prompted by the collision of Gondwana with Laurussia and subsequent formation of Pangea ∼320 Ma (Golonka 2007; Nance et al. 2014; Matthews et al. 2016) in the Permian. Suitable habitats were thus repeatedly fragmented, likely causing taxon distributional shifts and large-scale extinction events (Coccioni and Galeotti 1994; Zachos et al. 2001; Chen and Benton 2012). The consequences of these events could be potentially reflected in the mygalomorph phylogeny by the existence of many early-branching monogeneric lineages such as Microhexuridae, Porrhothelidae, Macrothelidae, Paratropididae and Stasimopidae, that do not seem to have any close relatives among extant taxa. Deep divergences far preceding the breakup of Pangea have also been observed for a similarly ancient and sedentary group – the velvet worms (Boyer et al. 2007; Murienne et al. 2014), indicating that paleoclimatic events and habitat heterogeneity played an important role in the evolution of these ancient groups.

We also detected potential long-distance dispersal events across the tree (denoted with “*”). The Australasian genus *Conothele* (Halonoproctidae), shows divergence between Australian and Vietnamese species dating back to approximately 46 Ma (47 – 44), which postdates their vicariance during the northward drifting of India ∼ 135 Ma (Will and Frimmel 2018). The genus exhibits some behavioral traits associated with ballooning (Main 1957), and it is possible that aerial dispersal played a role in *Conothele* distribution. Other cases involve Australian and African taxa. *Poecilomigas* (family Migidae) diverged from its Australia inhabiting sister taxon *Bertmainius* approximately 66 Ma (72 – 57), and *Kiama* and *Microstigmata* (both Microstigmatidae) split around 64 Ma (70 – 54). Both divergences are likely too young to reflect the East – West Gondwana vicariance (inferred by the biogeographic analyses) that started approximately 165 Ma, even when a possible bias in our dating analyses is considered. Trans-oceanic dispersal has previously been reported from the family Migidae (Harrison et al. 2017), which suggests that dispersal could plausibly explain the disjunct distribution between *Bertmainius* and *Poecilomigas*.

Another unusual pattern occurs within the family Pycnothelidae, where the closest relatives of the Australian genus *Stanwellia* have South American distributions. The split dates back to approximately 28 Ma (30 - 26) which by far postdates the East – West Gondwana vicariance but might actually reflect a connection between South America and Australia via Antarctica that presumably persisted until 35 Ma (Sanmartín and Ronquist 2004). The same vicariant scenario has been postulated to explain a similar distribution pattern recovered in Stag beetles (Kim and Farrell 2015). The biogeographic analyses also inferred vicariance for the *Stanwellia* – *Acanthogonatus* favoring South America – Australia connection over dispersal event.

### The Role of Silk in Prey Capture by Non-Araneomorph Spiders Revisited

As noted by Coyle (1986), progress in our understanding of the role silk plays in non-araneomorph spiders lags far behind what is known about araneomorphs; this was an accurate statement in 1986, and largely remains true today. As we discuss below, a prevailing perception considered by us misleading, if not erroneous, is that mygalomorph spiders do not use silk directly in foraging for prey; that is, it has been perceived that they do not spin webs defined in either a restricted or broad sense. For the purposes of our discussion we consider two characteristics as fundamental to ‘qualifying’ silk as playing a role in prey capture: 1) silk is employed by the spider in a sensory capacity to transmit information and extend prey sensing space outward to localize prey; and/or 2) silks can also be used by the spider to slow, impede, or restrain prey items. These two criteria are not mutually exclusive, particularly in mygalomorphs where visual clues are not known to play a significant role in prey capture; simply put silk is employed to increase predator speed (fast predator and/or slower prey (Coddington et al. 2019)). Note that Vollrath and Selden (2007) had a much broader definition of a web that included any spider structure made of silk. Because the question we address here is specifically related to the use of silk in prey capture, our working definition is narrower than theirs. It is our hypothesis, forwarded herein, that mygalomorphs, and consequently all spiders, plesiomorphically used silk to both localize and impede prey for capture. Whereas not all mygalomorphs use silken constructs to impede prey (criterion 2), nearly all employ it in some fashion to transmit vibrations to facilitate prey localization (criterion 1).

As documented in Figure 7, the MRCA for all mygalomorphs shows an unambiguous optimization for a funnel-and-sheet web versus trapdoor. Although the optimization of this character is equivocal at the root node (the MRCA of liphistiids and mygalomorphs), we would contend that the likely character state at this node, if araneomorphs and uraraneids (e.g., *Chimerarachne yingi*, (Wang et al. 2018)) were to be included in the analysis, would likely resolve as a funnel-and-sheet web. As noted by Vollrath and Selden (2007), there is a clear link between body structure and behavior in extant spiders. As such, the proportionally long spinnerets of spider sister taxon *Chimerarachne* suggest that they were funnel-and-sheet web builders (mygalomorph sheet web builders have proportionally longer spinnerets than their counterparts – see Ischnothelidae). Eskov and Selden (2005) hypothesized that *Permarachne* (at the time considered an extinct mesothele – now placed in the Uraraneida) constructed a funnel- and-sheet web but likely evolved *from* a protypical mesothele ancestor that constructed a trapdoor burrow with radiating lines. Alternatively, the uraraneid placement as the sister group to all spiders (Wang et al. 2018) likely requires the optimization that funnel-and-sheet webs are the plesiomorphic condition with trapdoor burrows independently derived in Liphistiidae – a lineage that is morphologically unique among spiders (see below). Funnel-and-sheet webs meet both criteria outlined above (Coyle 1986); that is, they transmit and extend prey localization from a tubular retreat (typically) and serve to entangle prey. Although silk may not have initially evolved for this function in the group’s common ancestor (that is, the common ancestor of Uraraneida + Araneae), these results strongly suggest that early silk use likely played a role in prey capture. Some authors have hypothesized that silk was initially used in reproduction (e.g., protecting eggsacs, sperm webs, etc. (Shultz 1987) or played a sensory role (Bond 1994)). However, these early functions may not be mutually exclusive; silken threads extending from an eggsac could have allowed a spider to care for her young while still able to sense prey (or predators) from a distance.

Irrespective of the uraraneid sheet web hypothesis originally proposed by Eskov and Selden (2005), the funnel-and-sheet web optimization at the mygalomorph node inferred here may be considered by some to be surprising given that the sister group to all other spiders (Opisthothelae), Mesothelae, build burrows covered by trapdoors. As we discuss in more detail below, a trapdoor covered burrow may have evolved from a sheet web (certainly the case among more derived mygalomorph taxa). The Mygalomorphae MRCA with high probability spun a sheet web from which trapdoor burrows were derived independently twice across the tree (once in the Atypoidea and a second time in the Bipectina). Nevertheless, we consider trapdoor burrow entrances to play a role in prey localization (discussed in more detail below), further enhanced in some liphistiids and other trapdoor spiders by extending the prey sensing surface of their burrow opening via silken trip lines or leaves, and other organic debris.

The evolutionary plasticity of the ancestral funnel-and-sheet web is best illustrated in the Atypoidea. The antrodiaetid clade includes *Antrodiaetus* (plus *Atypoides*), *Aliatypus*, and *Hexura* – all of which construct collared burrows, turreted burrows, trapdoors, and sheet webs, respectively. As demonstrated in a number of experiments conducted by Coyle (1986), the trapdoor spider *Aliatypus* likely suffers from reduced foraging efficiency relative to its more open burrow (collared door) and sheet web related genera. The sister group, Atypidae, builds purse webs, which are silken tubes that typically extend vertically up trees or horizontally along a substrate. As discussed by Coyle (1986), these spiders respond rapidly to vibrations along the tube surface; atypid purse webs definitively extend the spider’s prey sensory environment. An optimization of web constructs across the Atypoidea conducted by Hedin et al. (2019) recovered a similar pattern.

The Avicularioidea clade unequivocally has an MRCA that forages from a funnel-and-sheet web. As discussed in the classification section below, diplurids are polyphyletic, with most genera forming a grade of lineages that are sister to the remaining mygalomorphs. The characteristics of the ischnothelid, euagrid, etc. and hexathelid funnel-and-sheet webs are well documented for their abilities to aid in sensing/localizing prey, and in many taxa also impeding and/or entangling prey items. These silken constructs play a significant role in prey capture and are by any definition a “foraging web.” Funnel-and-sheet webs are independently derived twice in atracids (formerly hexathelids) and much further up the tree in the clade that includes “true” diplurids. The atracid funnel-and-sheet webs are highly modified from silken tubes (Gray 2010) and likely not homologous to the ancestral funnel-and-sheet web.

The Bipectina clade resolves as having an MRCA with a trapdoor. This clade includes typical trapdoor spiders like idiopids, migids, euctenizids, and nemesiids. As already discussed above, and in a number of papers (Coyle 1986; Coyle et al. 1992; Bond and Coyle 1995), trapdoor spiders employ silk in prey detection at the burrow entrance and in the door directly. Moreover, many trapdoor spider species included in this clade add additional silk lines, plant material, and tabs to the burrow entrance further emphasizing the role that the silken lined burrow entrance plays in detecting and localizing prey. Although trapdoor covered burrows do not seemingly serve to entangle or impede prey (criterion 2, above), aspects of the burrow do serve to enhance the sensory capacity of the spider. Trapdoors are independently lost in a number of Bipectina families in favor of other burrow constructs and foraging methods – Atracidae: funnel-and-sheet web; Theraphosidae: open burrows (though trapdoors have reappeared ≥3 times; Hamilton *persn. obsv.*); Pycnothelidae: open burrows, turrets; Dipluridae: funnel-and-sheet webs; Anamidae: open burrows and collar doors; Microstigmatidae: cursorial hunters, scored as inapplicable (*sensu* paratropidids). Additionally, species of one euctenizid genus, *Apomastus*, forages from an open burrow. In a number of these taxa, extensive field observations demonstrate that vibrations at the silken burrow entrance can be used to coax the spider to the surface, illustrating the functional role that silk plays in prey detection for these taxa.

As mentioned above, Coyle (1986) demonstrated that trapdoors appear to reduce foraging efficiency in some taxa. As such it begs the question as to why some mygalomorph spiders have abandoned funnel-and-sheet webs in favor of a foraging construct that appears to be less effective at detecting prey and clearly lacks the ability to impede or entangle prey items. Although this remains an open question, trapdoor spiders likely required additional protection from predators (e.g., parasitoid insects like pompilid wasps), desiccation, or even flooding. The reversal to an open burrow, often with an expanded collar for prey sensing (e.g., *Apomastus* burrows, (Bond 2004)), suggests that once environmental conditions or predator selective pressures change, the trapdoor can be easily lost.

It is our supposition that most mygalomorphs build foraging webs; that is, the majority of species employ silk either in a funnel-and-sheet web or at a burrow entrance to detect and localize prey and that a large number of taxa use silk to impede or entangle prey (the latter considered to be the plesiomorphic condition). These points are important to make particularly in light of a recent study by Fernández et al. (2018) that scored most mygalomorph taxa as having a non-foraging burrow. Based on the results and data discussed here, not surprisingly, we would question any analysis that hypothesizes mygalomorphs (and spiders as whole) to have a non-web foraging MRCA. Such an interpretation likely stems from limited taxon sampling and major differences in character state coding (Coddington et al. 2019). The latter in particular includes an interpretation that a spider foraging from a burrow is not employing silk in the process of prey capture, which, in our opinion contradicts the information available in the literature and direct observations in the field.

### Mygalomorphae Systematics and Higher-level Classification

The criterion of diagnosability, dictated by the ICZN, that all nominal taxa be diagnosable (morphologically or otherwise) presents some problems for higher level classification schemes based on these genomic data. Although one may recognize a molecularly defined lineage as a family rank taxon (with respect to other previously defined lineages at that equivalent rank), it is unreasonable to necessarily expect that some set of morphological features will reveal themselves as diagnostic for this newly identified lineage/taxon. A process of reciprocal illumination may lead to the discovery of such diagnostic characteristics, but that there are no guarantees. One might infer (based on character homoplasy) that the overriding notion of taxic homology, that diagnostic characters are used to unite groups (Panchen 1992), was an implicit driving force in previous attempts delineate mygalomorph families and other higher-level taxa (*sensu* Raven (1985)); an approach that at times seemingly interpreted diagnosability as synapomorphy. A diagnostic character(s) for a family does not necessarily infer synapomorphy any more than characters in a taxonomic key. A lineage that is hierarchically located at the same level among other family rank taxa should potentially be viewed as a family rank taxon regardless of whether or not it is morphologically diagnosable.

Taxonomic changes referenced herein are formally documented and new family-level taxa diagnosed in Appendix 1 (authorship to be formally attributed as Bond, Opatova, and Hedin 2019). We acknowledge that the taxonomic changes we propose are unlikely to go without criticism and will most likely require future revision and refinement. The four guiding principles we used attempted to realign family-level limits as clearly as possible but remain conservative in order to minimize errors: 1) revise higher-level mygalomorph classification and familial-level taxonomy such that it better reflects the evolutionary relationships indicated by data available (i.e., the sum of collected to data, not just this analysis); 2) further refine the classification, where possible, based on previous morphological/molecular hypotheses in which subfamilial taxa are clearly defined and supported by unambiguous synapomorphies; in some instances we transferred unsampled genera to newly defined families in cases where their placement in said group appears to be well supported in other studies; 3) minimize changes in family-level classification to the extent possible that maintains a consistent phylogenetic level across the tree for the family-level rank; and 4) instances where generic-level taxon sampling and criterion 2 above preclude an unambiguous family-level assignment of a genus (for relimited families), we retain the original family assignment but denote its placement as *incertae sedis* (operationally for the purposes of this study). We discuss below the restructuring of the major mygalomorph groups and their constituent families with a particular emphasis on longstanding problems in the families Dipluridae, Cyrtaucheniidae, and Nemesiidae.

#### Higher Level Groupings Atypoidea – Avicularioidea – Bipectina – Domiothelina Crassitarsae

The interpretation of the significance of similarities in spinneret morphology, constituted one of the center pieces in the debate revolving around the higher-level classification of the Mygalomorphae (Platnick and Gertsch 1976; Raven 1985; Eskov and Zonshtein 1990; Goloboff 1993) and taxon composition of some families (Raven 1985). Traditionally, mygalomorph spiders were divided into two families: Atypidae and Aviculariidae (=Theraphosidae; Simon 1864), later established as superfamilies Atypoidea and Ctenizoidea (Chamberlin and Ivie 1946). Atypoidea, a sister group to all the remaining mygalomorph families, consisted of Atypidae, Antrodiaetidae, and Mecicobothriidae. However, the external similarities of mecicobothriids and diplurids motivated some authors to reject the placement of Mecicobothriidae within Atypoidea (Platnick and Gertsch 1976), and subsequently even the concept of Atypoidina (Atypidae + Antrodiaetidae; Raven 1985) being a sister group to all remaining mygalomorphs was abandoned in favor of an alternative arrangement (Fig. 1a; Raven 1985). Eskov and Zonshtein (1990) disagreed with some of Raven’s character assessments and alternatively supported Atypoidea (*sensu* Chamberlin and Ivie (1946)), arguing that the shared traits in mecicobothriids and the diplurids, for example the elongation of spinnerets, constitute an ecological convergence related to a construction of a funnel-and-sheet web (Coyle 1986). However, their alternative hypothesis was rejected in what was the first computer-based cladistic treatment of mygalomorph relationships (Goloboff 1993), on the basis of different interpretation of Atypoidea unifying characters presented by Eskov and Zonshtein (1990).

Contrary to the prevailing morphology-based hypothesis (Raven 1985; Goloboff 1993), the first studies using molecular data strongly supported the classic Atypoidea hypothesis (Hedin and Bond 2006). The root node bifurcation of Mygalomorphae into Atypoidea and Avicularioidea was further established after adding additional molecular data (Ayoub et al. 2007; Bond et al. 2012). Although with limited taxon sampling, the same arrangement has been consistently recovered in subsequent studies employing transcriptome data (Bond et al. 2014; Garrison et al. 2016) and is also supported in our analyses. A recent study by Hedin et al. (2019); published while this study was in review, provided a much needed overhaul of the Atypoidea, recognizing two additional families (Hexurellidae and Megahexuridae).

The Avicularioidea clade was recovered in all the analyses with high support and a degree of internal resolution never achieved in past comprehensive mygalomorph phylogenetic studies. Similarly, to the results obtained in the past (Goloboff 1993; Hedin and Bond 2006; Ayoub et al. 2007; Bond et al. 2012), the concatenated analyses recovered a grade of early branching Avicularioidea lineages representing the so-called “non-diplurine diplurids” (Goloboff 1993), the Hexathelidae, Porrhothelidae and Macrothelidae. Alternatively, the species tree analysis performed in ASTRAL recovered these lineages, excluding Macrothelidae, in a single highly supported clade. We statistically explored alternative tree topologies (Table 2) to: 1) assess the monophyly of the family Dipluridae (*sensu* Raven (1985)), 2) test the species tree arrangement and 3) further test the monophyly of the family Hexathelidae (*sensu* Raven (1985)) within a wider outgroup sampling framework than in Hedin et al. (2018). The rejection of the alternative topologies by the AU topology tests, the reciprocal monophyly of the “non-diplurine diplurid” lineages correlated with a unique combination of morphological characters present in each taxon, provides strong support for taxonomic and nomenclatural changes proposed here and by other authors (Hedin et al. 2018).

As in the previous analyses (Hedin and Bond 2006; Ayoub et al. 2007; Bond et al. 2012), all Avicularioidea families, with exception of the grade of “diplurid” and former “hexathelid” lineages, were recovered in a single clade, close in taxon composition to the original delineation of Bipectina (*sensu* Goloboff (1993)). We recovered Paratropididae and Actinopodidae (removed from Bipectina by Bond et al. (2012)) alongside the Atracidae as part of that clade. The Bipectina including these three families represent the most inclusive monophyletic arrangement that is consistently recovered in all the analyses. For this reason, we choose to relimit the Bipectina clade (*sensu* Bond et al. (2012)) to include Actinopodidae, Atracidae, Stasimopidae, and Paratropididae alongside the Crassitarsae and Domiothelina.

A monophyletic group roughly consistent with Domiothelina (*sensu* Bond et al. (2012)) was recovered in all of our analyses. However, given the recent taxonomic relimitation of the family Ctenizidae (Godwin et al. 2018) and further taxonomic changes made in this paper, we formally recircumscribe the Domiothelina to include the following families: Halonoproctidae, Migidae, Idiopidae, Ctenizidae and Euctenizidae. We also relimit the family Ctenizidae to include only the genera *Cteniza* and *Cyrtocarenum* and establish a new family Stasimopidae (NEW FAMILY) to accommodate *Stasimopus* herein removed from Ctenizidae.

Albeit generally accepted as a monophyletic group (Hedin and Bond 2006; Bond et al. 2012), Crassitarsae (Raven 1985; Goloboff 1993) is an unstable clade in our analyses. The group was recovered in the concatenated analyses (supported in IQ-TREE ML inference and genomic/transcriptomic dataset), whereas other analyses supported an alternative placement for the Theraphosoidina clade (*sensu* Bond et al. (2012)), namely as sister to Domiothelina and the Stasimopidae/Atracinae/Actinopodidae clade (species tree analysis) or as the sister group to all Bipectina lineages (No_Paratropis dataset). Despite the uncertain status of Crassitarsae, both Theraphosoidina and the clade comprising Cyrtaucheniidae, Dipluridae, Microstigmatidae and former “Nemesiidae” lineages (the “Nemesioidina”, see below), were highly supported and mostly resolved at internal levels in all analyses. With the results at hand, we can now address the long-standing taxonomic issues involving the non-monophyly of the families Dipluridae, Cyrtaucheniidae and Nemesiidae. As a result, we recircumscribe the families Cyrtaucheniidae, Dipluridae and Nemesiidae and establish the new families Anamidae (NEW RANK), Entypesidae (NEW FAMILY), Bemmeridae (NEW RANK), and Pycnothelidae (NEW RANK).

#### “Dipluridae”

The monophyly of the diplurids was first questioned by Goloboff (1993) relatively soon after the last relimitation of the family (Raven 1985), which without a formal change being made remains still reflected in the current classification scheme. Diplurids in their original composition comprised a variety of, now in retrospect, very distantly related lineages possessing to some extent widely spaced “lower” spinnerets and elongated “upper” spinnerets (Simon 1889); which typifies the pitfalls of establishing groups of the basis of homoplastic, convergently evolved characteristics. Mecicobothriidae (Gertsch 1979) and Hexathelidae (Raven 1980) were subsequently removed from Dipluridae and established as independent families and a large number of taxa was then transferred to the family Nemesiidae (Raven 1985). Despite that, the elongation of posterior lateral spinnerets and the wide separation of posterior median spinnerets remained among characters uniting the family Dipluridae (Raven 1985). Goloboff (1993) debated the character state scoring of the spinneret elongation in diplurids, hexathelids, and mecicobothriids, alluding that the character definition was ambiguous, even within Dipluridae. Goloboff (1993) recovered the Dipluridae (*sensu* Raven (1985)) as four independent lineages which portended the future relimitation of diplurids to the subfamily Diplurinae (sensu Raven (1985)). The non-monophyly of the diplurids was also cautiously echoed by Coyle (1995) who, while conducting a taxonomic revision of the subfamily Ischnothelinae, did not find any strong synapomorphies establishing a sister relationship between ischnothelines and any other diplurid taxa (Coyle 1995). Diplurids have been recovered both as monophyletic (Ayoub et al. 2007; Bond et al. 2012) and paraphyletic (Hedin and Bond 2006; Wheeler et al. 2017; Hedin et al. 2018). However, with exception of Hedin et al. (2018), the taxon sampling was restricted to members of the subfamily Euagrinae and the seemingly contradictory outcome was likely a result of a lack of strong phylogenetic signal in the sequenced markers. In our analyses, we sampled representatives from three diplurid subfamilies (Diplurinae, Ischnothelinae, Euagrinae) plus *Microhexura*, a genus with uncertain placement (Raven 1985); Masteriinae was not included in this study. Our results are congruent with Goloboff’s (1993) hypothesis that Dipluridae constitutes an assemblage of distantly related lineages that do not share a common ancestor. The subfamilies Ischnothelinae, Euagrinae + *Microhexura* were recovered as three independent lineages at the base of the Avicularioidea clade (“non-diplurine diplurids” (Goloboff 1993; Hedin et al. 2018)), whereas Diplurinae taxa were placed as sister to cyrtaucheniids within proximity to nemesiids (Supplemental figure Fig. 9). The presumed synapomorphy of diplurids (sensu Raven (1985)), elongated and widely spaced spinnerets, could thus be an example of a retention of plesiomorphic condition (see discussion on silk use) of the “non-diplurine diplurids” that evolved convergently in Diplurinae – the genomic architecture likely already in place (see Uraraneida: *Chimerarachne yingi* (Wang et al. 2018)). As already pointed out by Coyle (1971) and Eskov and Zonshtein (1990), the traits related to the morphology of the spinnerets are likely correlated to the type of foraging structure produced by the spiders. “Diplurids” generally build conspicuous three-dimensional sheet-like webs with a funnel-like retreat (Coyle 1988; Coyle and Ketner 1990; Coyle 1995; Eberhard and Hazzi 2013; Passanha and Brescovit 2018), a trait present in many other mygalomorph taxa that have little to no evolutionary proximity to diplurids (e.g., Mecicobothriidae (Gertsch 1979), Macrothelidae (Snazell and Allison 1989; Shimojana and Haupt 1998), Atracidae (Gray 2010), and is also paralleled in araneomorph funnel-web spider family Agelenidae (Nentwig 1983). With the results of our phylogenetic analyses, it seems safe to assume that spinneret morphology is a plastic trait that has little to no informative value for *higher-level* classification in mygalomorph spiders. This observation is further supported by the results of the topology test rejecting the monophyly of Dipluridae (sensu Raven (1985)) as well as the monophyly of a clade consisting of the “non-diplurine diplurids” (Goloboff 1993), Hexathelidae and Porrhothelidae. We therefore substantially relimit the boundaries of the family Dipluridae (new circumscription). We elevate the subfamily Ischnothelidae (NEW RANK) and Euagridae (NEW RANK) to family-level and establish the new family Microhexuridae (NEW FAMILY).

#### “Nemesiidae”

The cosmopolitan family Nemesiidae belongs among the most speciose mygalomorph groups. As currently defined, it comprises over 400 nominal species that are placed in 45 genera. The family was diagnosed by Raven (1985) as those taxa possessing two rows of teeth on the paired tarsal claws and having scopulate tarsi (Raven 1985); a set of characters found in other groups and thus likely homoplasious, and later reinterpreted as plesiomorphic (Goloboff 1993). Not surprisingly, the monophyly of the family Nemesiidae has never been recovered in any broad scale molecular phylogenetic analyses of mygalomorphs (Hedin and Bond 2006; Ayoub et al. 2007; Bond et al. 2012). Moreover, even for analyses based solely on morphological characters, the family was seemingly split into five different lineages (Goloboff 1993). Paralleling the non-monophyletic nature of the family, the subfamilies established by Raven (1985) were also typically recovered as para- or polyphyletic (Goloboff 1993; Goloboff 1995; Hedin and Bond 2006; Bond et al. 2012). For this current study, we sampled five out of six nemesiid subfamilies (Raven 1985) and about 40% of the generic diversity to more thoroughly assess the composition of the family, particularly with an emphasis on resolving the position of the family Microstigmatidae and several difficult to place “cyrtaucheniid” genera (Goloboff 1993; Hedin and Bond 2006; Ayoub et al. 2007; Bond et al. 2012; Wheeler et al. 2017). Like all other studies conducted to date (Goloboff 1993; Hedin and Bond 2006; Ayoub et al. 2007; Bond et al. 2012; Garrison et al. 2016; Wheeler et al. 2017), we did not recover the Nemesiidae as monophyletic (Supplemental Fig. 9). The South African genus Spiroctenus was placed within Bemmeridae, whereas the remaining nemesiids were parceled among five reciprocally monophyletic lineages forming a clade with the cyrtaucheniids, diplurids, and microstigmatids (the “Nemesioidina” clade). Both microstigmatids and some “cyrtaucheniids” have been recovered as lineages embedded within Nemesiidae in the past (Goloboff 1993; Hedin and Bond 2006; Ayoub et al. 2007; Bond et al. 2012; Wheeler et al. 2017). Goloboff (1993), suggested that the some of the microstigmatid synapomorphies may be of neotenic nature (as proposed previously by Raven (1985) and Griswold (1985)), foreshadowing the eventual synonymy of microstigmatids with the nemesiids, or alternatively, splintering Nemesiidae into multiple families. Similar alternatives were also proposed for Cyrtaucheniidae (*sensu* Raven (1985)) based on the position of *Acontius, “Cyrtauchenius”*, and *Fufius* (Hedin and Bond 2006; Bond et al. 2012). Unfortunately, we cannot assess the position of *Acontius* and *Fufius* in this study. However, the sample representing “*Cyrtauchenius*” in Bond et al. (2012), in fact, corresponds to *Amblyocarenum*, later removed from its synonymy (Decae and Bosmans 2014). Although the phylogenetic placement of both genera could be conservatively interpreted as “embedded within Nemesiidae *sensu lato*, the placement of *Amblyocarenum along* with other genera likely has more severe taxonomic implications. *Amblyocarenum* is placed as a sole “cyrtaucheniid” within a monophyletic clade comprising the genus *Nemesia*, the type genus of the family. The affinities between *Amblyocarenum* and other nemesiids were also noted when the genus was removed from synonymy with *Cyrtauchenius* (Decae and Bosmans 2014). On the other hand, *Cyrtauchenius* forms a well-supported clade with two other cyrtaucheniid genera and is placed as sister to another monophyletic and morphologically distinguishable clade, the Dipluridae (*sensu* this study). The Dipluridae – Cyrtaucheniidae relationship and their placement within Nemesiidae was previously recovered in the pilot assessment of mygalomorph phylogeny with AHE data (Hamilton et al. 2016b), however, the sampling limitations of the study and the interpretation of *Fufius* and “*Linothele*” (later identified as *Diplura*) as taxonomically misplaced nemesiids alluded to a different conclusion than presented in this paper. The situation is similar for Microstigmatidae, presumably another “candidate” nemesiid lineage (Hedin and Bond 2006; Ayoub et al. 2007; Bond et al. 2012). The genus *Microstigmata* has been recovered in proximity to *Ixamatus* and another Australian nemesiid genus *Kiama* (Bond et al. 2012), which is congruent with our results. *Kiama* presents a unique combination of morphological features that, to some extent, overlaps with diagnostic characters for a handful of families, and as a result, the placement of the genus vacillated among Dipluridae (Main and Mascord 1969), Cyrtaucheniidae (Raven 1985) and Nemesiidae (Bond et al. 2012). Both *Kiama* and *Microstigmata* possess a uniquely modified cuticle and share a similar tarsal organ morphology (Raven 1985; Bond and Opell 2002), however these similarities were dismissed as convergent or plesiomorphic by others (Raven 1985; Goloboff 1993). The homology of these characters consequently needs to be reassessed in a framework using other lines of evidence other than just morphology (Bond et al. 2012; Hedin et al. 2018).

The dilemma of whether to dissolve the Nemesiidae, or redefine the family boundaries to include microstigmatids and cyrtaucheniids has been repeatedly discussed in the literature (Goloboff 1993; Hedin and Bond 2006; Bond et al. 2012), with tendencies gravitating towards the later solution (e.g. Bond et al. 2012; Fig. 1C in Hedin et al. (2018)). However, there were several factors at play reinforcing the argument for the “inclusive” Nemesiidae. First and foremost, the limited taxon sampling in the previous analyses (Hedin and Bond 2006; Ayoub et al. 2007; Bond et al. 2012; Hamilton et al. 2016b) partially skewed the interpretation of the results. The polyphyly of the family Cyrtaucheniidae (*sensu* Raven (1985)), misplacement of *Cyrtauchenius* (see above) and lack of adequate representation of the diplurines (*sensu* Raven (1985)) presented major hurdles for accurately interpreting nemesiid relationships. Recovering *Microstigmata* or a few genera of an apparently polyphyletic family embedded within an enlarged “nemesiid” construct understandably supported the notion that a broad synonymization (lumping rather than splitting) might be the more stable and thus preferable taxonomic solution. Adding to the overall ambiguity of the results, large sections of the recovered trees often lacked support (Hedin and Bond 2006; Ayoub et al. 2007; Bond et al. 2012; Wheeler et al. 2017), adding uncertainty to the monophyly of the groups in question. We would suggest that our improved sampling and high support values brings long-needed insight and resolution to the “nemesiid” problem, which is essential for ultimately answering the question of whether or not the Nemesiidae should be fragmented into independent families or not. The family Nemesiidae (*sensu* Raven (1985)) has lacked a well-supported synapomorphy, a widely recognized fact (Goloboff 1993; Goloboff 1995; Bond et al. 2012; Harvey et al. 2018), since the family was first delimited (Raven 1985). On the other hand, some groups (e.g., subfamilies) currently recognized within the family, do possess diagnostic synapomorphic characters (Harvey et al. 2018), which is also the case for the diplurids and cyrtaucheniids (*sensu* this study) that were recovered as mutually exclusive lineages within the larger more inclusive “Nemesioidina” clade. We thus argue that the original concept of the family (*sensu* Raven (1985)) must be abandoned in favor of establishing multiple monophyletic families that can also be ultimately clearly defined morphologically. Another fact in favor of such an arrangement is the relatively deep divergences separating the lineages (Kuntner et al. 2018). Major lineages across the entire clade diversified into their respective individual groups between 115 – 79 Ma, roughly congruent with dated divergences among other families (e.g., Theraphosidae and Barychelidae split around 107 Ma and Actinopodidae diverged from Atracidae approximately 98 Ma). In agreement with the results of our analyses, we therefore relimit the family Nemesiidae (new circumscription) as outlined in the Appendix 1.

## Conclusions And Future Directions

It is our hope that this study represents a significant leap forward in stabilizing mygalomorph spider classification via a robust, well-supported phylogenetic framework. Extensive taxon sampling combined with enhanced phylogenetic resolution using genomic-scale data facilitates the proposal of an alternative classification scheme that more accurately reflects the evolutionary relationships of the infraorder. Consequently, these results brought to their logical fruition require extensive changes to existing familial composition as well as the establishment of a number of new family-rank taxa. Understandably, such changes require considerable downstream scrutiny and criticism. This newly derived framework is a hypothesis that begs further testing through additional data and increased taxon sampling.

One observation in particular from this study that seems clear is that morphological data are not uniformly suitable for reconstructing mygalomorph higher-level relationships, and to a slightly lesser degree for delimiting family rank taxa (e.g., subfamily relationships seem to be relatively stable as evidenced by the number of subfamilies elevated to family rank). Raven’s (1985) results were in no small part afflicted by the rampant homoplasy that seemingly plagues this group of spiders. Given the data and lack of computational resources available when the analyses were conducted, the results Raven achieved represented the strongest prevailing hypothesis at that time and provided a framework upon which to test new hypotheses. In 1985 and shortly thereafter (Eskov and Zonshtein 1990) disagreement regarding morphological evidence for various clades and familial-level relationships and identity were in retrospect matters of subjective opinion and consequently difficult to test (likely explaining why some seemingly correct hypotheses, e.g., Atypoidea, were ignored). Alternatively though, it is important to recognize that a relatively large percentage of the new family rank taxa were once morphologically defined subfamilies – a number of which predate Raven (1985) – and thus relatively correct.

Issues regarding spider classification notwithstanding, phylogenomics provides an exciting new framework for considering broad scale questions related to spider evolution. Our biogeographic and dating analyses suggest that ancient vicariance and dispersal shaped the currently observed biogeographic patterns. Much of the deep phylogenetic structure across the infraorder may be attributable to climatic heterogeneity and oscillation prior to the breakup of Gondwana/Pangea. Such deep phylogenetic and ancient divergence, coupled with low vagility, makes these spiders ideal candidates to test biogeographic hypotheses.

Furthermore, the recently held notion that most mygalomorph spiders capture prey from non-foraging burrows (Fernández et al. 2018) has critical implications for how we might understand and interpret silk across spiders as a whole. The reformulated hypothesis posits that spiders likely foraged primitively from a silken funnel-and-sheet web that served classically-held web functions like increasing predator speed and enhancing prey capture success. How mygalomorph spiders build their webs and retreats, employ silk in prey capture, and the nature of their spinning apparatus remains a rich area of study that has received little attention since Coyle’s (1986) treatment.

Considerable work remains in achieving final resolution of mygalomorph relationships and stable fully-resolved familial composition. Despite our best efforts, which included a combination of transcriptomic and AHE data, we are still unable to resolve several of the deepest nodes; Paratropididae remains a difficult taxon to place as do the newly formed Stasimopidae. Despite our best efforts and major relimitation, many nemesiid, diplurid, and cyrtaucheniid genera remain in an uncertain taxonomic state. Although we could speculate about placement of many taxa (e.g., Aporoptychinae cyrtaucheniids, Masteriinae diplurids, as well as numerous South American nemesiids), such questions will have to be left to future studies. At the very least, it is our hope that this study facilitates further interest and provides a solid foundation for future work in this intriguing, and ancient group of spiders.

## Acknowledgements

We thank M. Arnedo, R. Booysen, A. Decae, M. Forman, N. Garrison, R. Godwin, J. Grismer, C. Haddad, S. Huber, V. Hula, P. Just, J. Malumbres-Olarte, T. Mazuch, JA. Neethling, J. Rodriguez, F. Šťáhlavský, J. Starrett, C. Stephen, and D. Weinmann for help with field collecting and providing additional samples. We are grateful to M. Harvey, M. Kuntner, C. Griswold, S. Zonshtein, B. Wiegmann, J. Ledford, and an anonymous reviewer for comments on earlier versions of the manuscript. The computational analyses were completed in part with resources provided by the Auburn University Hopper supercomputing cluster. This project was funded in part by National Science Foundation grants DEB-0841610 to JEB, Doctoral Dissertation Improvement Grant DEB-1311494 to CAH, Czech Science Foundation grant 16-10298S to JK, Auburn University Department of Biological Sciences and College of Sciences and Mathematics (JEB), an Auburn University Cellular and Molecular Biosciences Peaks of Excellence Research Fellowship (CAH), and Evert and Marion Schlinger Foundation, University of California Davis (JEB).

## Appendix 1: Revised Taxonomy for some of the families of the Spider Infraorder Mygalomorphae (Araneae)

### Supplemental Files

Data available from the Dryad Digital Repository: http://dx.doi.org/10.5061/dryad.[NNNN]

#### Tables and Figures

Table S1: Specimen locality data

Table S2: Specimens used in the combined analyses of genomic and transcriptomic data (DNAAA_matrix). AHE dataset: specimens sequenced in this study, AA dataset: data proceeding from Garrison et al. (2016) “BCC-75” matrix.

Table S3: Character coding for web type ancestral state reconstruction

Supplemental Figure 1. Tree topology obtained in maximum likelihood (ML) analyses conducted in RAxML. Values on the nodes correspond to bootstrap support.

Supplemental Figure 2. Tree topology obtained in Bayesian inference (BI) analyses conducted in ExaBayes. Values on the nodes correspond to the Bayesian posterior probability (PP).

Supplemental Figure 3. Tree topology obtained in maximum likelihood (ML) analyses conducted in IQ-TREE. Values on the nodes correspond to SH-aLRT support.

Supplemental Figure 4. Phylogenetic tree inferred after removal of Paratropididae terminals (“No_Paratropis” dataset). Tree topology obtained in maximum likelihood (ML) analyses conducted in RAxML. Boxes near the branches denote full bootstrap support values; numbers report bootstrap support values obtained in this analysis (black) and the analysis of the full “All_taxa” dataset (red). Asterisks mark differences in topology obtained from the “All_taxa” dataset.

Supplemental Figure 5. Tree topology obtained in species tree analysis conducted in ASTRAL. Values on the nodes correspond to the ASTRAL support values.

Supplemental Figure 6. Comparison of resulting tree topologies obtained by concatenated analyses conducted in RAxML and ExaBayes (left) and species tree approach conducted in ASTRAL (right). Asterisks mark differences in topologies. Doted grey line denote array of lineages forming a highly supported clade in species tree analysis. Boxes on nodes denote support values obtained in each approach, color coded according to distinct support level categories depicted in bottom left corner in following order (left to right): RAxML bootstrap support, ExaBayes Bayesian posterior probabilities, ASTRAL support values.

Supplemental Figure 7. Divergence time estimates of mygalomorph spiders inferred by treePL on a topology obtained in RAxML, values near the nodes correspond to the inferred timing of the divergences. Numbered nodes represent placement of the calibrations and their minimum and maximum ages (in brackets), 1) Mygalomorphae (Garrison et al. 2016; Fernández et al. 2018), 2) Avicularoidea: *Rosamygale grauvogely* (Selden and Gall 1992), 3) “Nemesioidea”: *Cretamygale chasei* (Selden 2002), 4) Atypoidea: *Ambiortiphagus ponomarenkoi* (Eskov and Zonshtein 1990), 5) *Ummidia* (Wunderlich 2011). The x-axis is time in million years.

Supplemental Figure 8. Divergence time estimates of mygalomorph spiders inferred by treePL on a topology obtained in RAxML, values near the nodes correspond to the confidence intervals (CI) for the highest posterior density (HDP) inferred from 100 bootstrap replicates. The x-axis is time in million years.

Supplemental Figure 9. Phylogenetic tree of Mygalomorphae summarizing results from concatenated and species tree approaches within a framework of a current classification scheme (World Spider Catalog 2018). Topology obtained in the Maximum Likelihood analyses. Boxes on nodes denote support values obtained in each approach (left to right): RAxML bootstrap support, ExaBayes Bayesian posterior probabilities (PP), ASTRAL support values, IQ-Tree SH-aLRT support values. Colour of the boxes corresponds to distinct support level categories depicted in bottom left corner. One filled box indicates full support in all analyses; white box=topology not recovered in species tree analysis.

#### Data Files

ASTRAL_all_taxa.tre

ASTRAL_all_taxa.trees

ExaBayes_all_taxa.tre

RASP_ingroup.tre_dis.csv

RAxML_DNAAA.nex

RAxML_DNAAA.tre

RAxML_DNAAA_partitions.txt

RAxML_ExaBayes_all_taxa.nex

RAxML_ExaBayes_all_taxa_partitions.txt

RAxML_No_Paratropis.nex

RAxML_No_Paratropis.tre

RAxML_No_Paratropis_partitions.txt

RAxML_all_taxa.tre

silkUseAncStateRecon.txt

treePL_dated.tre

#### Families remaining unchanged in their composition

##### Actinopodidae – Atracidae (The Venom Clade)

###### Remarks

The family Atracidae comprises three genera endemic to eastern Australia. Formerly designated as the “hexathelid” subfamily Atracinae (Gray 2010), the group was recently elevated to family level by Hedin et al. (2018) and supported by the topology test performed on our dataset. Contrary to the hypothesis based on morphological characters (Raven 1985; Goloboff 1993), Atracidae has consistently been recovered as sister to Actinopodidae (Hedin and Bond 2006; Ayoub et al. 2007; Bond et al. 2012; Hamilton et al. 2016b; Hedin et al. 2018), further corroborated by our results. Actinopodidae comprises three genera with a disjunct South American – Australian distribution and on the basis of morphological characters (eyes widely spread across the carapace), was placed as sister to Migidae (Raven 1985; Goloboff 1993). However, this relationship has never been supported in any molecular phylogenetic study. Despite the differences in the morphology of the families, the phylogenetic proximity of Atracidae/Actinopodidae is supported not only by the overwhelming amount of evidence proceeding from molecular data, but also by similarities in venom composition (for details see Hedin et al. (2018). Certain members of both families are medically important; the bites of the Sydney funnel-web spider, *Atrax robustus*, cause lethal or life-threatening conditions, and other taxa can also be dangerous to humans (Isbister et al. 2005, 2015). The Atracidae/Actinopodidae clade was recovered within the Bipectina, which contradicts the results of a number of previous analyses (Goloboff 1993; Hedin and Bond 2006; Bond et al. 2012). Unfortunately, the exact relationships of the Atracidae/Actinopodidae with other taxa remain uncertain. The group was consistently placed in all analyses as sister to the Stasimopidae, although with varying degrees of support (see below). In our analyses, both families were represented by two out of three genera; *Atrax* and *Hadronyche* for Atracidae, and *Actinopus* and *Missulena* for Actinopodidae, however, the monophyly of both families, based on complete generic sampling, appears to be strong (Hedin et al. 2018).

##### Antrodiaetidae – Megahexuridae – Atypidae – Mecicobothriidae – Hexurellidae

###### Remarks

Atypoid spider classification has recently undergone major revision with the designation of new families and phylogenetic structure; see Hedin et al. (2019).

##### Barychelidae

###### Remarks

Barychelidae is a highly diverse family (42 genera, 495 species) with a Gondwanan distribution spanning continental islands and also remote islands of volcanic origin (Raven 1994) indicating the ability to disperse. Like Theraphosidae, Barychelidae possess well developed scopulae and dense claw tufts, but differs in the number of cuspules, clavate trichobothria, and the shape of the maxillary anterior lobe (Raven 1985; Dippenaar-Schoeman 2002). Due to the potential ambiguity of these characters (Raven 1985; Goloboff 1993; Bond et al. 2012) some authors have considered Barychelidae, at least in part, a paraphyletic lineage of Theraphosidae (Goloboff 1993). Stemming from differences in character state interpretations, taxa tend to vacillate between the two families, with proposed membership by some authors in one family (Raven 1985; Guadanucci 2014) to be later rejected by others (Gallon 2002; Schmidt 2002; Ríos Tamayo 2017). One example of this problem, the family placement of *Brachionopus* (Raven 1985; Gallon 2002; Schmidt 2002) was not successfully resolved until the implementation of a molecular approach (Lüddecke et al. 2018). With our limited sampling, we recovered the reciprocal monophyly of both Barychelidae and Theraphosidae. However, a thorough sampling of both families will be needed to stabilize the family boundaries. Barychelidae (as currently defined) is composed of two common and divergent “phenotypes” (a “naked” group that is much less setose and resemble most mygalomorph groups, and a “hairy” group that resemble small theraphosids. These two “phenotypes” have been the major drivers of taxa historically bouncing back and forth between the two families.

##### Euctenizidae

###### Remarks

The group was originally designated as a subfamily of Cyrtaucheniidae (Raven 1985), but with increasing amount of evidence for a non-monophyletic Cyrtaucheniidae (Bond and Opell 2002; Bond and Hedin 2006), Euctenizidae were elevated to family level (Bond et al. 2012). The family comprises 76 species within seven genera across a North American distribution (Bond 2017). We sampled all described genera and recovered the family as monophyletic, as well as the sister group to Ctenizidae (*sensu* this study) and Idiopidae.

##### Halonoproctidae

###### Remarks

The family currently comprises six genera and has a nearly cosmopolitan distribution. It was recently removed from Ctenizidae (Godwin et al. 2018), whose monophyly was doubted both on morphological (Raven 1985) and molecular grounds (Hedin and Bond 2006; Bond et al. 2012; Opatova et al. 2013). Similar to Godwin et al. (2018), we recovered Halonoproctidae as monophyletic in all analyses. Furthermore, the topology test rejected the monophyly of the family Ctenizidae (*sensu* Raven (1985)) as well as the monophyly of Halonoproctidae + Ctenizidae (*sensu* this study), verifying the status of the family within the context of a wider taxonomic framework. It should be noted that we did not include the genus *Bothriocyrtum* in our analyses, but given its stable position as sister to *Hebestatis* (Hedin and Bond 2006; Bond et al. 2012; Opatova et al. 2013; Godwin et al. 2018), we believe that it had little to no effect on the resulting topology and support values.

##### Hexathelidae – Porrhothelidae – Macrothelidae

###### Remarks

The monophyly of the family Hexathelidae in its original composition (*sensu* Raven (1985)) has been challenged on molecular grounds on numerous occasions (Hedin and Bond 2006; Ayoub et al. 2007; Bond et al. 2012; Opatova and Arnedo 2014a; Hamilton et al. 2016b). Formal taxonomic changes were put on hold until the implementation of UCEs alongside complete generic sampling which resulted in parceling of the family Hexathelidae (*sensu* Raven (1985)) into four distinct families (Hedin et al. 2018). We sampled all the families in our analyses and similar to the results of Hedin et al. (2018), we recovered the families Hexathelidae, Porrhothelidae and Macrothelidae among the early branching lineages near the Avicularioidea root; though Atracidae, was recovered within the Bipectina clade sister to Actinopodidae. The family Hexathelidae now comprises nine genera with a disjunct distribution (Australia, New Zealand, South America), of which we only sampled the Australian *Bymainiella*. It was recovered as sister to the family Euagridae with high support, which was previously indicated by EF-1g data (Ayoub et al. 2007), albeit the support for this relationship was low. Both Porrhothelidae and Macrothelidae were recovered as stand-alone lineages placed near the Avicularioidea root node. Both lineages are monogeneric and were previously considered sister taxa (Raven 1985), but this relationship was statistically rejected in the past (Opatova and Arnedo 2014a) and their unique position appears stable even in light of considerable genomic data ((Hedin et al. 2018); this study). Given the wide taxonomic framework of our analyses, we tested the monophyly of the family Hexathelidae, in its original composition (*sensu* Raven (1985)), as well the alternative arrangement recovered in the species tree analysis (Table 2); both were statistically rejected, confirming Hedin et al. (2018).

Note. Because the new family Porrhothelidae was electronically published but not registered in ZooBank, the online registration system for the ICZN it is redscribed here.

Family Porrhothelidae (NEW FAMILY) Hedin and Bond

**Type genus:** *Porrhothele* Simon, 1892 [urn:lsid:nmbe.ch:spidergen:00021] (type species *Porrhothele antipodiana* (Walkenaer, 1837).

##### Diagnosis

As a consequence of its monotypy, characters used to diagnose the family Porrhothelidae are those characters attributed to the type genus. *Porrhothele* was thoroughly diagnosed and described by Forster (1968) with additions by Raven (1980). Members of this family can be diagnosed on the basis of the following unique combination of characters: 1) small posterior sternal sigilla; 2) single row of promarginal cheliceral teeth; 3) male tibia swollen with dense pattern of strong promarginal spines (illustrated in Forster (1968) e.g., p. 170, figs. 543– 548). This monotypic family is found in New Zealand.

##### Idiopidae

###### Remarks

Idiopids are another family with predominantly Gondwanan distribution (South America, Asia, Africa, Australia and New Zealand), though some species are also known from the Middle East and Central America. The family is well characterized by male palpal synapomorphies (Raven 1985) and its monophyly has never been called into question by previous studies (Goloboff 1993; Hedin and Bond 2006; Ayoub et al. 2007; Bond et al. 2012). Therefore, it comes as no surprise that the family was recovered as monophyletic in our analyses. Idiopids are among the most diverse of the mygalomorphs; the family comprises 22 genera and three subfamilies: Arbanitinae, Genysinae and Idiopinae (Raven 1985; Rix et al. 2017c), of which we did not sample Genysinae. The position of Idiopidae within the Domiothelina clade was consistently recovered by both morphological (Goloboff 1993) and molecular approaches (Bond et al. 2012), but given the incomplete sampling of the family Ctenizidae (*sensu* Raven (1985)), the position within the clade varied. In our analyses, the family was placed as sister to Ctenizidae and Euctenizidae with high support.

##### Migidae

###### Remarks

The family Migidae currently comprises 11 genera with a classical Gondwanan distribution (Australia, New Zealand, Africa, Madagascar and South America). The family is well defined morphologically as taxa having longitudinal keels on cheliceral fangs and lack a rastellum (Raven 1985; Griswold and Ledford 2001). Similar to previous studies (Hedin and Bond 2006; Bond et al. 2012), our taxon sampling is rather limited. We only sampled four genera, lacking a neotropical representative; therefore, we cannot assess the monophyly of the group. The family has been recovered as sister to either *Stasimopus* (Hedin and Bond 2006) or Halonoproctidae (Bond et al. 2012), as well as in close proximity to Idiopidae and Euctenizidae (Ayoub et al. 2007; Bond et al. 2012), as we report here. In the concatenated analyses, the family was recovered with high support as sister to all the remaining Domiothelina taxa, except for Halonoproctidae. Despite the congruent topology, this relationship was poorly supported in the species tree analysis.

##### Paratropididae

###### Remarks

Paratropididae is a poorly known, and highly distinct family comprising four genera known from Central and South America. Its phylogenetic placement has been regarded as “difficult” (Bond et al. 2012), given the conflict between the morphology-based hypotheses and the results of molecular analyses. The family was originally placed within a clade “Theraphosoidina”, alongside the Theraphosidae and Barychelidae (Raven 1985; Goloboff 1993), however this relationship has never been recovered in any molecular analyses (Hedin and Bond 2006; Bond et al. 2012; Hamilton et al. 2016b) and, in fact, a close relationship between Paratropididae and any other mygalomorph taxon has never been supported. Instead, the paratropidids were placed among the grade of early branching Avicularioidea (Hedin and Bond 2006; Bond et al. 2012). In our analyses, the paratropidids represent a stand-alone lineage diverging at the root of Bipectina, however, similar to previous studies (Hedin and Bond 2006; Bond et al. 2012), we only sampled representatives of the genus *Paratropis*. This not only precludes the evaluation of the family’s monophyly, but may also be a mitigating factor in placing it on the tree. Moreover, the stand-alone status of the lineage may also have an impact on the inferred tree topology as a whole, a fact demonstrated by the improved support values at particular nodes and the topology changes in the unsupported nodes after paratropidids were removed from the dataset (Supplemental Fig. 4). Our exploratory analysis combining AHE and transcriptomic data recovered paratropidids with high support as sister to the “Nemesioidina” clade (Fig. 5). However, the sampling in this analysis was limited both in terms of taxa and data occupancy. Despite the equivocal nature of the placement of *Paratropis* within the Bipectina, it seems clear that there is no support for its placement within Theraphosoidina (*sensu* Raven (1985)).

##### Theraphosidae

###### Remarks

The family Theraphosidae is the most diverse family of mygalomorphs with a nearly cosmopolitan distribution. Currently comprising 144 genera and over 970 species, the family is well-defined (aside from Barychelidae) by dense claw tufts and leg scopulae (Raven 1985). The monophyly is generally undoubted (Hedin and Bond 2006; Bond et al. 2012; Lüddecke et al. 2018; Turner et al. 2018), although paraphyly with respect to Barychelidae has occurred due to taxon misplacement (Goloboff 1993; Bond et al. 2012). Until recent molecular phylogenetics work (Lüddecke et al. 2018; Turner et al. 2018), attempts to reconstruct the internal relationships of Theraphosidae mostly relied on the analysis of morphological characters (Pérez-Miles 1994; Pérez-Miles et al. 1996; Pérez-Miles 2002; Guadanucci 2005, 2011; Guadanucci 2014). Molecular approaches have become more common to resolve taxonomic issues at shallow levels, such as species delimitation or uncovering species-specific phylogeographic patterns (Hamilton et al. 2011; Hendrixson et al. 2013; Hamilton et al. 2014; Graham et al. 2015; Montes de Oca et al. 2016; Ortiz et al. 2018). In our analyses, we sampled eight genera from North and South America, Africa, and Asia, recovering the family as monophyletic in all the analyses. The subfamily Theraphosinae, here represented by the tribe Theraphosini (*Aphonopelma*, *Cyrthopholis*, *Sericopelma* and *Stichoplastoris*) (Turner et al. 2018), was also recovered with high support. On the other hand, our results substantiate the para- or polyphyletic status of the subfamily Ischnocolinae, already debated on the basis of morphological characters (Raven 1985; Guadanucci 2005; Guadanucci 2014; Guadanucci and Wendt 2014) and karyotype structure (Král et al. 2013). The genus *Catumiri* (Ischnocolinae) was recovered as sister to the rest of the theraphosid lineages, whereas *Ischnocolus*, regarded as the most “basal group” of the family (Raven 1985), appears to be related to Ornitoctoninae and Harpactirinae (here represented by *Brachionopus*). The non-monophyletic status of higher-level taxonomic units, and apparent polyphyly of some wide spread genera arises from high levels of homoplasy in the morphological characters traditionally used in Theraphosidae taxonomy (Guadanucci and Wendt 2014; Gabriel and Longhorn 2015; Hamilton et al. 2016a; Turner et al. 2018). Molecular approaches and more thorough taxonomic sampling will be essential to establishing an accurate classification scheme reflecting the evolutionary relationships within the family.

*\*denotes genera included in analyses in the accompanying paper (Opatova et al. 2019); @ denotes that the taxon was definitively placed by another study supporting the designated placement here*

#### Relimitation of the family Dipluridae Simon, 1889

##### Family Dipluridae Simon, 1889 (new circumscription)

In our analyses, the family Dipluridae is represented by three South American genera *Diplura*, *Linothele* and *Harmonicon* belonging to the diplurid subfamily Diplurinae (Raven 1985). We recovered the Dipluridae as sister to Cyrtaucheniidae placed within the same clade alongside Nemesiidae. These results are roughly congruent with Goloboff (Goloboff 1993), who analyzed four genera of diplurids (*sensu* Raven (1985)) and recovered *Diplura* as the sister to the rest of the Bipectina clade. As in Raven (1985), Goloboff (1993) also noted shared characters in *Diplura* and the Crassitarsae, and hypothesized that the diplurids would be restricted to the subfamily Diplurinae in the future. Unfortunately, we did not conduct an exhaustive sampling of the “diplurid” genera and acknowledge that even after removing the Euagridae, Ischnothelidae and Microhexuridae the family likely remains non-monophyletic. Given the rough congruence between Raven’s diplurid subfamilies (Raven 1985) and the lineages removed from the Dipluridae in this study, it is possible that the distinct subfamily Masteriinae (Raven 1985; Passanha and Brescovit 2018) will be elevated to family status as well.

**Type genus: *Diplura*** C. L. Koch, 1850 (type species *Mygale macrura* Koch, 1841)

###### Remarks

The family Dipluridae is relimited here to include members of the subfamilies Diplurinae Simon, 1889 and Masteriinae Simon, 1889. Of the Diplurinae genera, *Trechona* C. L. Koch, 1850 and *Troglodiplura* Main, 1969 were not included in our analysis but are nevertheless retained, at least until such time that they are included in a molecular study; *Trechona’s* inclusion appears justified based on stridulatory and palpal endite characters (Raven 1985), whereas Main (1993) justifies the placement of *Troglodiplura* in Diplurinae contra Raven (1985). The relimited family herein conservatively also includes Masteriinae which we predict is likely to be recognized as a standalone family, closely related to Microhexuridae at some point in the future; the subfamily was recently reviewed by Passanha and Brescovit (2018), however, the authors provide no insight regarding affinities to other mygalomorph taxa. The relimited Dipluridae will require a significantly revised diagnosis at some point in the future.

List of included subfamilies and genera

**Diplurinae Simon, 1889**

*\*Diplura* C. L. Koch, 1850

*\*Harmonicon* F. O. Pickard-Cambridge, 1896

*\*Linothele* Karsch, 1879

*Trechona* C. L. Koch, 1850

*Troglodiplura* Main, 1969

**Masteriinae Simon, 1889**

*Masteria* C. L. Koch, 1873

*Siremata* Passanha and Brescovit, 2018

*Striamea* Raven, 1981

##### Family Ischnothelidae F.O.P Cambridge, 1897 NEW RANK

The family corresponds to the former diplurid subfamily Ischnothelinae, defined by six morphological synapomorphies (Raven 1985; Coyle 1995). In our analyses, we sampled the genera *Ischnothele* (South America) and *Thelechoris* (Africa) and recovered these together in all analyses. In previous analyses based on morphological characters, *Ischnothele* was placed as a sister group to Bipectina (Goloboff 1993), but our results suggest that ischnothelids belong among the first groups that diverged within the Avicularioidea clade. Given the morphological cohesiveness of the group, and the fact that it was thoroughly revised in the past (Coyle 1995), we retain all the genera of the subfamily Ischnothelinae (Raven 1985; Coyle 1995) despite our incomplete sampling. The family Ischnothelidae comprises five genera (*Andethele*, *Indothele*, *Ischnothele*, *Latrothele*, *Thelechoris*) with a disjunct distribution spanning across Central and South America, the Antilles, Africa, Madagascar, India and Sri Lanka. All genera build conspicuous three-dimensional webs and often exhibit subsocial behavioral (Coyle 1995).

**Type genus: *Ischnothele*** Ausserer, 1875 (type species *Ischnothele caudata* Ausserer, 1875)

###### Remarks

Our sampling of ischonothelines comprised two genera that span the taxonomic breadth of the subfamily (Coyle 1995, Figures 21, 22) but omit three others. As discussed by Coyle (1995) in his revision of the subfamily, the group is supported by a number of unique morphological characters that taken in combination can be used for diagnosis. Distinguishing characteristics for this newly elevated family are: 1) an elongate cymbial apophysis; 2) collariform trichobothrial bases; and 3) fused silk spigots.

List of included genera

*\*Ischnothele* Ausserer, 1875

*Andethele* Coyle, 1995

*Indothele* Coyle, 1995

*Lathrothele* Benoit, 1965

*\*Thelechoris* Karsch, 1881

##### Family Euagridae Raven, 1979 NEW RANK

The family roughly corresponds to the former subfamily Euagrinae (Raven 1985) that currently comprises 11 genera with distribution spanning across Australia, Asia, Africa, and South and North America. Euagrinae differ from other “diplurids” by a combined lack of cuspules and broad and short serrula. In our analyses, we only sampled five Euagrinae genera: *Allothele*, *Australothele*, *Cethegus*, *Euagrus* and *Namirea*, which were all recovered forming a monophyletic group towards the root node of Avicularioidea. Similar results were obtained in previous analyses (Ayoub et al. 2007; Bond et al. 2012), though with limited taxon sampling. The internal relationships of these taxa resemble their morphological affinities (Raven 1985). The “austral genera” possess a crescent of hirsute cuticle at the base of the posterior median spinnerets (“australothelinae crescent”) and were recovered as a monophyletic group sister to *Euagrus* that lacks this character (Raven 1985; Coyle 1988). The lack of a strong synapomorphy uniting the “austral” genera and *Euagrus* + *Phyxioschema* (Raven 1985; Coyle 1988) alongside deep divergence dating back to nearly 169 Ma (174 – 161), would provide the support for the division of Euagrinae in two different families. However, the decision would be premature given our taxon sampling, which represents less than half of the described generic diversity of Diplurinae, and in particular, lacks representatives of the subfamily Masteriinae (Raven 1985) – a group that overlaps in some characters with Euagrinae. We therefore prefer to elevate the subfamily Euagrinae to family rank, retaining its current generic composition – i.e., the taxa sampled in this paper along with the genera *Caledothele*, *Carrai*, *Chilehexops*, *Leptothele*, *Phyxioschema*, *Stenygrocercus* and *Vilchura*.

**Type genus: *Euagrus*** Ausserer, 1875 (type species *Euagrus mexicanus* Ausserer, 1875)

###### Remarks and revised diagnosis

Unfortunately, our sampling of Euagrinae diplurids is depauperate (and should include other *Euagrus* species, see below) and thus our placement of the remaining diplurids in the subfamily Euagrinae follows authors subsequent to Raven’s (1985) establishment of the family (i.e., inclusion in the subfamily is maintained per Raven (1991), Coyle (1986), Raven and Schwendinger (1995), and Ríos-Tamayo and Goloboff (2017)). Note that *Euagrus*, was considered by Coyle (1988) to be largely central/south American with the two species, *Euagrus atropurpureus* (S. Africa) and *Euagrus formosanus* (Taiwan) as misplaced. It is important to recognize that some of these taxa may be allocated to other formerly “diplurid” families at some point in the future (e.g., *Chilehexops, Caledothele, Vilchura*). Although morphological evidence (*sensu* Coyle (1988)) seems strong that *Phyxioschema* is the sister group to *Euagrus*, the morphological cladistic analysis conducted by Ríos-Tamayo and Goloboff (2017) provides little encouragement that the subfamily is monophyletic and should indicate that any morphological diagnosis of this remaining assemblage of taxa is likely without substance. Following Raven (1985), the family can be diagnosed as those taxa having a distinct spur on the second male tibia and a centrally raised tarsal organ (also following from Ríos-Tamayo and Goloboff (2017)); Euagrinae taxa differ from australothelines by lacking an australotheline crescent (see details in the diagnosis below).

List of included Euagrinae genera

*\*Euagrus* Ausserer, 1875

*Caledothele* Raven, 1991

*Chilehexops* Coyle, 1986

*Leptothele* Raven and Schwendinger, 1995

*Phyxioschema* Simon, 1889

*Vilchura* Ríos-Tamayo and Goloboff, 2017

#### Australothelinae NEW SUBFAMILY Bond, Opatova, and Hedin

**Type genus: *Australothele*** Raven, 1984 (type species *Australothele maculata* Raven, 1984)

##### Diagnosis

Per Raven (1985), australothelines can be diagnosed from other euagrids as those taxa having an “australotheline crescent” on the posterior median spinnerets – which comprises a “crescent of hirsute cuticle isolated by pallid glabrous anterior to bases” and lacking cymbial spines (the latter character is shared by numerous other mygalomorph taxa (mentioned by Ríos-Tamayo and Goloboff (2017)). As noted by Raven (1985), a similar such crescent is also found in the putatively true diplurid genus *Masteria*. The subfamily is also supported as monophyletic in the analysis conducted by Ríos-Tamayo and Goloboff (2017) and Hedin et al. (2018).

List of genera included

*\*Australothele* Raven, 1984

*\*Allothele* Tucker, 1920

*Carrai* Raven, 1984

*\*Cethegus* Thorell, 1881

*\*Namirea* Raven, 1984

*Stenygrocercus* Simon, 1892

#### Microhexuridae NEW FAMILY Bond, Opatova, and Hedin

*Family Microhexuridae (NEW FAMILY)*.—Microhexuridae is a monogeneric family comprising two nominal species endemic to North America. Both species are very small (> 6 mm) and have allopatric distributions restricted to the high-elevation peaks in the Appalachian Mountains and Pacific Northwest Cascade Range (Coyle 1981; Hedin et al. 2015). Despite being recognized as a diplurid (*sensu* Raven (1985)), *Microhexura* exhibits a combination of characters not present in other diplurid genera. Most notably *Microhexura* possess a longitudinal fovea and spinnerets somewhat shorter than in other diplurids, which made the genus difficult to place (Coyle 1981; Raven 1985). The unique position of the genus noted in the previous studies (Coyle 1981; Raven 1985) is further supported by our results that recovered the genus *Microhexura* as a stand-alone lineage among other “non-diplurine diplurids” at the base of the Avicularioidea clade. The topology test results reject both its placement within the Dipluridae (*sensu* Raven (1985)), but also within the “non-diplurine diplurids”/Hexathelidae/Porrhothelidae clade recovered in the species tree analyses.

**Type genus. *Microhexura*** Crosby and Bishop, 1925 (type species *Microhexura montivaga* Crosby and Bishop, 1925)

##### Diagnosis

Because the family is monogeneric, characters used to diagnose Microhexuridae are those that can be attributed to the genus by Coyle (1981). Among the smallest mygalomorph spiders known, the family is diagnosed on the basis of the following unique combination of characters: 1) longitudinal fovea; 2) four spinnerets – posterior lateral spinnerets long (nearly the length of the abdomen); 3) lacking abdominal tergites; and 4) male pedipalp lacking conductor.

List of included genera

*\*Microhexura* Crosby and Bishop, 1925

### Relimitation of the family Ctenizidae Thorell, 1887

#### Ctenizidae Thorell, 1887 (new circumscription)

The generic composition of the family Ctenizidae was dramatically altered by Raven’s (1985) morphology-based cladistic analysis, but it became apparent that the family most likely comprised an assemblage of diverse lineages that do not share a common ancestor (Hedin and Bond 2006). This observation was repeatedly confirmed by subsequent molecular-based studies (Bond et al. 2012; Opatova et al. 2013; Wheeler et al. 2017). However, a lack of perceived phylogenetic signal, incomplete sampling, or a combination thereof precluded previous authors from making taxonomic changes that would rectify the problem. The polyphyletic nature of the family was partially resolved by Godwin et al. (2018) when six genera were transferred to the family Halonoproctidae, circumscribing Ctenizidae to *Cteniza*, *Cyrtocarenum* and *Stasimopus*. Godwin et al. (2018) recognized that *Stasimopus* did not share a common ancestor with the remaining genera, but incomplete outgroup sampling limited their ability to definitively address the phylogenetic placement of the taxon. In our analyses, we recovered *Cteniza* and *Cyrtocarenum* as sister taxa, which agrees with previous findings (Opatova et al. 2013; Godwin et al. 2018). *Cteniza* and *Cyrtocarenum* were placed as sister to the family Euctenizidae, whereas *Stasimopus* was recovered outside of Domiothelina in a weakly supported clade as the sister lineage to Atracidae and Actinopodidae. Given the distant phylogenetic position of *Stasimopus*, with regards to the remaining ctenizids, and considering that the topology test rejected the monophyly of the three genera, we remove *Stasimopus* from the family Ctenizidae and designate it to the family Stasimopidae (NEW FAMILY). Ctenizidae is circumscribed to contain only the genera *Cteniza* and *Cyrtocarenum*. The family Ctenizidae thus now constitutes a much smaller family comprising six species with a predominantly Mediterranean distribution.

**Type genus**: ***Cteniza*** Latreille, 1829 (type species *Cteniza sauvagesi* (Rossi, 1788))

##### Remarks and revised diagnosis

As discussed by others (e.g., Decae 1996; Godwin et al. 2018), *Cteniza* and *Cyrtocarenum* are virtually indistinguishable from each other and are very similar to other taxa like halonoproctids and stasimopids, save some questionable eye group characteristics. As outlined by Godwin et al. (2018), Ctenizidae, as delimited herein, can be distinguished from other similar taxa by lacking curved spines on the anterior legs, having a more arched and rounded caput and flatter ventral posterior carapace, and having a more oblong abdomen (rather than ovoid); legs tend to be less stocky. Ctenizids differ from their sister group, Euctenizidae, by lacking scopulae.

List of included genera

*\*Cteniza* Latreille, 1829

*\*Cyrtocarenum* Ausserer, 1871

### Stasimopidae NEW FAMILY Bond, Opatova, and Hedin

Although the placement of *Stasimopus* within the family Ctenizidae (Raven 1985) was repeatedly called into question based on earlier molecular studies (Hedin and Bond 2006; Bond et al. 2012; Opatova et al. 2013; Godwin et al. 2018), establishing its exact phylogenetic position has been difficult. In previous phylogenetic studies of the entire infraorder, *Stasimopus* was recovered as sister to Migidae (Hedin and Bond 2006), or as the sister group to the entire Domiothelina, albeit with low support (Bond et al. 2012). Importantly, when morphological characters were added to the analyses, *Stasimopus* was recovered within the Ctenizidae (Bond et al. 2012), confirming the homoplasious nature of some morphological characters. Our results place *Stasimopus* as a stand-alone lineage either forming a clade with Atracidae and Actinopodidae or as sister to Domiothelina. The sister relationship with the “Venom Clade” was also recovered by Hamilton et al. (2016b), though in a much reduced AHE dataset, and similarly to our results, the clade was poorly supported. The support values for this clade increased with the removal of the paratropidid terminals (Supplemental Fig. 4) and with addition of transcriptomic data (Fig. 4), suggesting that adding more data to the matrix could be instrumental for stabilizing Stasimopidae placement. Despite the partial positional uncertainty of *Stasimopus*, the topology test rejects its inclusion within Ctenizidae. Considering the results of our analyses, alongside the genus’ restricted distribution to southern Africa – a biodiversity hotspot with high levels of endemicity (Myers et al. 2000; Mittermeier et al. 2011), it seems that there is sufficient evidence to support the hypothesis that *Stasimopus* represents a unique lineage within the Mygalomorphae and deserves family-level status.

**Type genus: *Stasimopus*** Simon, 1892 (type species *Stasimopus caffrus* (C. L. Koch, 1842)

#### Diagnosis

At this time, the family Stasimopidae is monogeneric, thus characteristics used to diagnose the genus are the distinguishing characters for the family. Per Raven (1985) and summarized by Engelbrecht and Prendini (2012), stasimopids can be distinguished from other ctenizid-like taxa on the basis of the following combination of unique characters: 1) lacking a saddle-like depression on tibia III; 2) having a very wide eye group (twice as wide as it is long); and 3) palpal endites with a modified anterior lobe (described as “produced”). Based on the breadth of morphological diversity contained within the genus, which includes somewhat bizarre carapace modifications (Engelbrecht and Prendini 2012), it is possible that *Stasimopus* could be split in to multiple genera at some point in the future.

List of included genera

*\*Stasimopus* Simon, 1892

### Relimitation of the family Cyrtaucheniidae Simon, 1889

#### Cyrtaucheniidae Simon, 1889 (new circumscription)

Following the results of our phylogenetic analyses, we transfer the genus *Homostola* to the newly established family Bemmeridae (NEW RANK, see below). We also formally transfer the Mediterranean genus *Amblyocarenum* to the family Nemesiidae. The family Cyrtaucheniidae is thus relimited to nine genera (three of them monotypic) distributed across Africa, Asia and South America. In this study, we analyzed two African genera: *Ancylotrypa*, *Cyrtauchenius* and one South American lineage that is morphologically proximate to *Acontius*. As a result, Cyrtaucheniidae were recovered as monophyletic in all analyses and placed as sister to Dipluridae. At present, we retain the genera *Acontius*, *Anemesia*, *Angka*, *Bolostromoides, Bolostromus*, *Fufius*, and *Rhytidicolus* (not included in this study), within Cyrtaucheniidae. We acknowledge that the family still likely remains poly- or paraphyletic, because *Fufius* is known to have affinities with the Nemesiidae (Bond et al. 2012), however, we argue that even the partial Cyrtaucheniidae taxon pruning performed in this study is preferable to retaining the high level of polyphyly. This is particularly true when obtaining fresh samples may be exceptionally difficult, especially for the African and South American representatives of the genera *Acontius* and *Bolostromus*.

**Type genus: *Cyrtauchenius*** Thorell, 1869 (type species *Cyrtauchenius terricola* (Lucas, 1846))

##### Remarks

Long recognized as a polyphyletic assemblage of genera, we herein relimit Cyrtaucheniidae to the Aporoptychinae minus *Kiama* and *Angka* based on Raven’s (1985) original composition of the subfamily. Given the diversity of the taxa left remaining in the family, it is unlikely that a revised diagnosis will have any lasting value (if even possible to formulate, given the diverse array of taxa remaining) and as such we decline to provide one. We anticipate significant changes to the composition of the family as additional aporoptychines are added to the tree. We justify our transfer of *Angka* to Microstigmatidae based on the original description of the genus by Raven and Schwendinger (1995), where they considered it as a clear sister taxon to *Kiama* (also transferred, by us below, to the family Microstigmatidae); the transfer of *Amblyocarenum* to Nemesiidae is justified on the basis of its phylogenetic position.

List of included genera

*\*Cyrtauchenius* Thorell, 1869

*\*Ancylotrypa* Simon, 1889

*Incertae sedis*

*Acontius* Karsch, 1879

*Anemesia* Pocock, 1895

*Bolostromoides* Schiapelli and Gerschman, 1945

*Bolostromus* Ausserer, 1875

*Fufius* Simon, 1888

*Rhytidicolus* Simon, 1889

Transferred to other families (not included in the relimitation of Cyrtaucheniidae)

*\*Amblyocarenum* Simon, 1892 here transferred to Nemesiidae justified on the basis of phylogenetic placement.

*^@^Angka* Raven and Schwendinger, 1995 here transferred to Microstigmatidae on the basis of its putative sister relationship to *Kiama*.

*\*Homostola* Simon, 1892 here transferred to Bemmeridae justified on the basis of phylogenetic placement.

### Relimitation of the family Nemesiidae Simon, 1889

#### Nemesiidae Simon, 1889 (new circumscription)

The family Nemesiidae is a cosmopolitan group. Our sampling comprised five genera, including the type genus *Nemesia*, spanning species sampled from three continents (Africa, Europe and North America). These genera form a single lineage and are placed in a clade that is sister to all the former “nemesiid” lineages plus Cyrtaucheniidae, Dipluridae and Microstigmatidae. In agreement with our results, we transfer the genus *Spiroctenus* to the newly established family Bemmeridae and transfer *Amblyocarenum* here from Cyrtaucheniidae (see discussion on Cyrtaucheniidae for justification). We remove the genera *Pionothele*, *Bayana*, *Stenoterommata*, *Stanwellia*, *Acanthogonatus, Pycnothele* and *Entypesa* and transfer them into newly erected families Pycnothelidae (NEW RANK) and Entypesidae (NEW FAMILY). We further remove the taxa included in the Anamini tribe (Harvey et al. 2018) and elevate it to the rank of family: Anamidae (NEW RANK). We also transfer the genera *Kiama, Angka*, and *Xamiatus* to the family Microstigmatidae (see details below). We acknowledge that many taxa, particularly those inhabiting South America and Asia, remain severely under-sampled in our study and therefore we cannot speak to their definitive placement at this time; as such they remain for time being assigned to Nemesiidae *incertae sedis*.

**Type genus: *Nemesia*** Audouin, 1826 (type species *Nemesia cellicola* Audouin, 1826)

##### Remarks

To a greater extent than Cyrtaucheniidae (above), what remains of the family Nemesiidae is a likely para-polyphyletic assemblage of genera. As such we consider the placement of a large number of genera in the family to be uncertain at best; many of which may prove to be pycnothelids. Consequently, much work remains, particularly if we are able to place many of the South American genera.

List of included genera

*\*Nemesia* Audouin, 1826

*\*Amblyocarenum* Simon, 1892

*\*Calisoga* Chamberlin, 1937

*\*Iberesia* Decae and Cardoso, 2006

*\*Mexentypesa* Raven, 1987

Incertae sedis

*Atmetochilus* Simon, 1887

*Brachythele* Ausserer, 1871

*Chaco* Tullgren, 1905

*Chilelopsis* Goloboff, 1995 – most likely Pycnothelidae

*Damarchilus* Siliwal, Molur, and Raven, 2015

*Damarchus* Thorell, 1891

*Diplothelopsis* Tullgren, 1905

*Flamencopsis* Goloboff, 1995

*Gravelyia* Mirza and Mondal, 2018

*Hermacha* Simon, 1902 – likely to be transferred to Entypesidae

*Hermachura* Mello-Leitāo, 1923

*Lepthercus* Purcell, 1902 – likely Entypesidae

*Longistylus* Indicatti and Lucas, 2005; removed from Anamidae *contra* Indicatti and Lucas (2005)

*Lycinus* Thorell, 1894

*Neostothis* Vellard, 1925

*Prorachias* Mello-Leitão, 1924

*Psalistopoides* Mello-Leitão, 1934

*Pselligmus* Simon, 1892

*Rachias* Simon, 1892; likely to be transferred to Pycnothelidae per Goloboff (1995)

*Raveniola* Zonshtein, 1987

*Sinopesa* Raven and Schwendinger, 1995

Transferred to other families (not included in the relimitation of Nemesiidae)

*Xamiatus* Raven, 1981; here transferred to Microstigmatidae on the basis of its putative sister/close relationship with *Ixamatus* and *Kiama* (Raven 1981,1985)

*Ixamatus* Simon, 1887 here transferred to Microstigmatidae on the basis of its putative phylogenetic position per Harvey et al. (2018).

#### Anamidae Simon, 1889 NEW RANK

The family Anamidae corresponds to the Australian endemic tribe Anamini (Raven 1985; Harvey et al. 2018) formerly placed within the “nemesiid” subfamily Anaminae (Raven 1985).

The family comprises genera *Aname*, *Chenistonia*, *Hesperonatalius*, *Kwonkan*, *Namea*, *Proshermacha*, *Teyl*, *Teyloides* and *Swolnpes*, of which we analyzed six (including *Pseudoteyl* currently synonymized with *Teyl*). The family formed a monophyletic lineage in all analyses and was recovered as sister to Entypesidae/Microstigmatidae clade. The phylogenetic relationships of the group were recently evaluated, employing both extensive taxonomic sampling and multilocus data (Harvey et al. 2018). Harvey et al. (2018) performed a complete generic sampling of the Australian Anamini and recovered the group as a well-supported clade, also including genera not sampled in our analyses. Following the results of our study and Harvey et al. (2018), we therefore transfer all the genera listed above into the family Anamidae. Morphology and biogeography do not support the inclusion of *Longistylus* Indicatti and Lucas, 2005 in the Anamidae contra the tentative hypothesis proposed by Indicatti and Lucas (2005).

**Type genus: *Aname*** L. Koch, 1873 (type species *Aname pallida* L. Koch, 1873)

##### Remarks

A detailed diagnosis for this new family rank taxon is provided by Harvey et al. (2018).

List of included genera

*\*Aname* L. Koch, 1873

*\*Proshermacha* Simon, 1908

*^@^Hesperonatalius* Castalanelli et al. 2017 per Harvey et al. (2018)

*\*Kwonkan* Main, 1983 (=*\*Yilgarnia* Main, 1986)

*\*Namea* Raven, 1984

*^@^Chenistonia* Hogg 1901 per Harvey et al. (2018)

*^@^Swolnpes* Main and Framenau, 2009 per Harvey et al. (2018)

*\*Teyl* Main, 1975

*^@^Teyloides* Main, 1985 but check Bond et al. (2012) and Hedin and Bond (2006) – may be able to place elsewhere) Harvey et al. (2018)

#### Bemmeridae Simon, 1903 NEW RANK

The family Bemmeridae comprises 35 species classified in the genera *Homostola* and *Spiroctenus*, both endemic to South Africa. The group was repeatedly recovered as sister to Theraphosoidina (Hedin and Bond 2006; Bond et al. 2012), which is further substantiated by our analyses. Previous analyses also suggested the inclusion of the genus *Ancylotrypa* (Hedin and Bond 2006), an assumption that was unfortunately based on sample misidentification. The genus *Homostola* is being transferred to this family from Cyrtaucheniidae, the genus *Spiroctenus* from Nemesiidae. Both genera comprise medium-size spiders, constructing underground burrows with or without a trapdoor.

**Type genus**: ***Spiroctenus*** Simon, 1892 (type species *Spiroctenus personatus*, Simon, 1888)

##### Remarks and Diagnosis

The family name is derived from the available family-level taxon Bemmereae Simon, 1903; *Bemmeris* Simon, 1903 is a junior synonym of *Spiroctenus*. The identity and consequently family-level placement of *Spiroctenus* and *Homostola* is contentious and likely to remain so. Raven (1985) considered the two genera unrelated and placed in separate families (Nemesiidae and Cyrtaucheniidae respectively), attributing Hewitt’s (1915) hypothesized synonym of the two to a misidentification. To date, it would seem that a male *Homostola* specimen has never been described in detail, further complicating the situation (but see Hewitt (1916)). Based on careful specimen identification and locality data, we are confident that our sample comprises both genera. The *Spiroctenus* exemplar represents an identical male and female sequence of *Spiroctenus flavopunctatus* (Purcell, 1903); *Homostola* terminals are likely *H. pardalina affin*. (Hewitt, 1913) and *H. vulpecula* Simon, 1892. Based on our examination of specimens, Bemmeridae females can be diagnosed on the basis of the following unique combination of characters: 1) eyes on a low tubercle; 2) metatarsus and tarsus with moderate to thick scopula; 3) preening combs on leg III; 4) labium with relatively dense patch of cuspules; and 5) a rastellum comprising a row of thickened, blunt spines not borne on an apophysis. Male *Spiroctenus* species have a very distinct mating clasper on leg I comprising a number of large spines and an apophysis at the tibia/metatarsus junction; we would predict a similar morphology for *Homostola* males.

List of included genera

*\*Homostola* Simon, 1892

*\*Spiroctenus* Simon, 1889

#### Pycnothelidae Chamberlin, 1917 NEW RANK

*Family Pycnothelidae (NEW RANK)*.—The family Pycnothelidae is newly established here in order to accommodate taxa formerly belonging to a diverse mix of “nemesiid” subfamilies (Raven 1985). Some of the close relationships are not entirely novel. For example the sister relationship between *Stanwellia* and *Acanthogonatus* was previously recovered by Hedin and Bond (2006). We recovered the African genus *Pionothele*, formerly placed within the subfamily Bemerinnae, as sister to *Spiroctenus*, and sister to all pycnothelid diversity. The family distribution spans across Africa, Australia, South America and New Zealand, therefore it could be expected that more genera will be eventually transferred here following a thorough assessment, particularly of African and South American lineages.

**Type genus: *Pycnothele*** Chamberlin, 1917 (type species *Pycnothele perdita* Chamberlin, 1917)

##### Remarks and Diagnosis

The genus *Pycnothele* was not included in our analysis but hypothesized morphological affinities (Goloboff 1995), have attributed it to be closely aligned with *Stanwellia*, *Acanthogonatus*, and *Stenoterommata*. Harvey et al. (2018) stated: “If a close relationship between *Stanwellia*, *Acanthogonatus*, *Stenoterommata* and *Pycnothele* is maintained, as proposed by Goloboff (1995), the subfamily-group name Pycnothelinae Chamberlin, 1917 is available for this clade (Chamberlin 1917).” In the interests of taxonomic stability, we believe the conservative decision is to transfer *Pycnothele* from Nemesiidae and include it in our newly defined clade (justified on the basis of morphological similarity). The alternative would be to propose a new family rank name (e.g., “Stanwellidae”) which will be inevitably deprecated at a later time. At this time, the family Pycnothelidae can be diagnosed on the basis of the following unique combination of characters: 1) rastellum absent; 2) aspinose tarsi; 3) preening combs absent, although reduced in some *Stanwellia* and *Acanthogonatus*; 4) labium lacks cuspules; 5) tibial spur on male leg I absent; and 6) most taxa have a reduced or absent inferior tarsal claw. We anticipate that this diagnosis might change as other nemesiids (noted as *incertae sedis* above) are added to the tree.

List of included genera

*^@^Pycnothele* Chamberlin, 1917

*\*Stanwellia* Rainbow and Pulleine, 1918

*\*Acanthogonatus* Karsch, 1880 (note: most likely polyphyletic)

*\*Bayana* Pérez-Miles, Costa and Montes de Oca, 2014

*\*Pionothele* Purcell, 1902

*\*Stenoterommata* Holmberg, 1881

#### Entypesidae NEW FAMILY Bond, Opatova, and Hedin

We establish the family Entypesidae to accommodate the genus *Entypesa*, which currently comprises 7 species known from Madagascar and South Africa. The genus was originally placed within the nemesiid subfamily Anaminae (Raven 1985), however this taxon has been shown to be polyphyletic (Bond et al. 2012). Despite the rather undesirable monogeneric state of the family, we believe that establishing Entypesidae as a family is justified by both its reciprocal monophyly in relation to its sister taxon Microstigmatidae and the deep divergence between them, dating back to 79 Ma. We believe the monogeneric nature of the family is only temporary and other genera (particularly fellow South African genera *Lepthercus* and *Hermacha*) will eventually be transferred here following a thorough assessment of Nemesiidae phylogeny.

**Type genus: *Entypesa*** Simon, 1902 (type species *Entypesa nebulosa* Simon, 1902)

#### Diagnosis

Because the family is monogeneric, characters used to distinguish it are those that can be attributed to diagnosing the genus. Based on examination of specimens, Raven’s (1985) description and Dippenaar-Shoeman (2002), Entypesidae may be diagnosed on the basis of the following unique combination of characters: 1) narrow straight thoracic groove; 2) distinctly raised eye tubercle; 3) labium lacks cuspules; 4) weak rastellum comprising numerous short, stout bristles; 5) dense scopula on tarsi I-III; 6) leg III preening comb, present; and 7) long, digitiform spinnerets. We anticipate the inclusion of *Hermacha* and *Lepthercus* in this family at some point in the future.

Included genus

*\* **Entypesa*** Simon, 1902

### Relimitation of the family Microstigmatidae Roewer, 1942

#### Family Microstigmatidae Roewer, 1942 (new circumscription)

The family Microstigmatidae is well defined by morphological characters, such as small, round booklung openings and pustulose cuticle (Raven 1985; Hedin and Bond 2006). The monophyly of the family has never been assessed within a robust molecular framework because all previous studies exclusively included the South African genus *Microstigmata* (Hedin and Bond 2006; Ayoub et al. 2007; Bond et al. 2012; Wheeler et al. 2017); but see (Goloboff 1993). Following the results of our analyses we relimit the family Microstigmatidae to include the genus *Kiama*. The morphological similarities in tarsal organ and cuticle structure linking *Kiama* to microstigmatids are well documented (Raven 1985; Bond and Opell 2002). Similar characteristics are also found in the genera *Ixamatus* and *Angka* (Bond et al. 2012) suggesting their close relationships to the microstigmatids.

**Type genus: *Microstigmata*** Strand, 1932 (type species *Microstigmata geophila* (Hewitt, 1916))

##### Remarks

The family Microstigmatidae was traditionally diagnosed as taxa have possessing, circular booklung openings. With Goloboff’s (1995) inclusion of *Pseudonemesia* and *Spelocteniza* in the family, as well as *Envia* (*sensu* Ott and Höfer (2003)), that feature now only serves to diagnose a subset of these relatively unique genera. Although a scaly or pustulose cuticle may be a diagnostic character uniting all microstigmatids (Bond and Opell 2002), future SEM study of all genera will be necessary to confirm. As noted above, the transfer of *Angka* to the Microstigmatidae is based on its affinities with *Kiama* whose phylogenetic position clearly aligns here.

List of included genera

*\*Microstigmata* Strand, 1932

*^@^Angka* Raven and Schwendinger, 1995

*Envia* Ott and Höfer, 2003

*^@^Ixamatus* Simon, 1887 transferred here per Harvey et al. 2018; as noted by Raven (1985), the genus has a pustulose cuticle, a potentially diagnostic character of the family.

*\*Kiama* Main and Mascord, 1969

*Micromygale* Platnick and Forster, 1982

*Ministigmata* Raven and Platnick, 1981

*Pseudonemesia* Caporiacco, 1955

*Spelocteniza* Gertsch, 1982

*^@^Xamiatus* Raven, 1981 transferred here on the basis of morphological similarity with *Ixamatus* and its putative sister group relationship with *Kiama*; as noted by Raven (1985), the genus has a pustulose cuticle, a potentially diagnostic character of the family.

*Xenonemesia* Goloboff, 1989

## Literature cited

Aberer AJ, Kobert K, Stamatakis A. 2014. ExaBayes: massively parallel Bayesian tree inference for the whole-genome era. Mol Biol Evol, 31:2553–2556.

Andújar C, Faille A, Pérez-González S, Zaballos JP, Vogler AP, Ribera I. 2016. Gondwanian relicts and oceanic dispersal in a cosmopolitan radiation of euedaphic ground beetles. Mol.Phylogenet.Evol., 99:235–246.

Anisimova M, Gil M, Dufayard J-F, Dessimoz C, Gascuel O. 2011. Survey of branch support methods demonstrates accuracy, power, and robustness of fast likelihood-based approximation schemes. Syst Biol, 60:685–699.

Ayoub NA, Garb JE, Hedin M, Hayashi CY. 2007. Utility of the nuclear protein-coding gene, elongation factor-1 gamma (EF-1g), for spider systematics, emphasizing family level relationships of tarantulas and their kin (Araneae: Mygalomorphae). Mol.Phylogenet.Evol., 42:394–409.

Beaulieu JM, O’Meara BC, Donoghue MJ. 2013. Identifying hidden rate changes in the evolution of a binary morphological character: the evolution of plant habit in campanulid angiosperms. Syst Biol, 62:725–737.

Beutel E, Nomade S, Fronabarger A, Renne P. 2005. Pangea’s complex breakup: A new rapidly changing stress field model. Earth and Planetary Science Letters, 236:471–485.

Boger SD. 2011. Antarctica—before and after Gondwana. Gondwana Research, 19:335–371.

Bolotov IN, Vikhrev IV, Bespalaya YV, Gofarov MY, Kondakov AV, Konopleva ES, Bolotov NN, Lyubas AA. 2016. Multi-locus fossil-calibrated phylogeny, biogeography and a subgeneric revision of the Margaritiferidae (Mollusca: Bivalvia: Unionoida). Mol.Phylogenet.Evol., 103:104–121.

Bond J, Opell BD. 2002. Phylogeny and taxonomy of thegenera of south-western North American Euctenizinae trapdoor spidersand their relatives (Araneae: Mygalomorphae, Cyrtaucheniidae). Zool. J. Linn. Soc., 136:487–534.

Bond JE. 1994. Seta-spigot homology and silk production in first instar Antrodiaetus unicolor spiderlings (Araneae: Antrodiaetidae). J. Arachnol.:19–22.

Bond JE. 2004. Systematics of the Californian euctenizine spider genus *Apomastus* (Araneae: Mygalomorphae: Cyrtaucheniidae): the relationship between molecular and morphological taxonomy. Invertebr Systemat, 18:361–376.

Bond JE. 2017. Euctenizidae. In Ubick, D., Paquin, P. Cushing, P., Roth, V. (eds.) Spiders of North American identification manual, 2nd Edition, 55–57. New Hampshire, American Arachnological Society.

Bond JE, Coyle FA. 1995. Observations on the natural history of an *Ummidia* trapdoor spider from Costa Rica (Araneae, Ctenizidae). J. Arachnol., 23:157–164.

Bond JE, Garrison NL, Hamilton CA, Godwin RL, Hedin M, Agnarsson I. 2014. Phylogenomics resolves a spider backbone phylogeny and rejects a prevailing paradigm for orb web evolution. Current Biology, 24:1765–1771.

Bond JE, Hedin M. 2006. A total evidence assessment of the phylogeny of North American euctenizine trapdoor spiders (Araneae, Mygalomorphae, Cyrtaucheniidae) using Bayesian inference. Mol.Phylogenet.Evol., 41:70–85.

Bond JE, Hedin MC, Ramirez MG, Opell BD. 2001. Deep molecular divergence in the absence of morphological and ecological change in the Californian coastal dune endemic trapdoor spider *Aptostichus simus*. Mol Ecol, 10:899–910.

Bond JE, Hendrixson BE, Hamilton CA, Hedin M. 2012. A Reconsideration of the Classification of the Spider Infraorder Mygalomorphae (Arachnida: Araneae) Based on Three Nuclear Genes and Morphology. PLoS ONE, 7:e38753.

Bond JE, Stockman AK. 2008. An integrative method for delimiting cohesion species: Finding the population-species interface in a group of Californian trapdoor spiders with extreme genetic divergence and geographic structuring. Syst Biol, 57:628–646.

Bouckaert R, Vaughan TG, Barido-Sottani J, Duchene S, Fourment M, Gavryushkina A, Heled J, Jones G, Kuhnert D, De Maio N. 2018. BEAST 2.5: An Advanced Software Platform for Bayesian Evolutionary Analysis. BioRxiv:474296.

Boufford DE, Spongberg SA. 1983. Eastern Asian-eastern North American phytogeographical relationships-a history from the time of Linnaeus to the twentieth century. Ann Mo Bot Gard, 70:423–439.

Boyer SL, Clouse RM, Benavides LR, Sharma P, Schwendinger PJ, Karunarathna I, Giribet G. 2007. Biogeography of the world: a case study from cyphophthalmid Opiliones, a globally distributed group of arachnids. J Biogeography, 34:2070–2085.

Castalanelli MA, Huey JA, Hillyer MJ, Harvey MS. 2017. Molecular and morphological evidence for a new genus of small trapdoor spiders from arid Western Australia (Araneae: Mygalomorphae: Nemesiidae: Anaminae). Invertebr Systemat, 31:492–505.

Chamberland L, McHugh A, Kechejian S, Binford GJ, Bond JE, Coddington J, Dolman G, Hamilton CA, Harvey MS, Kuntner M. 2018. From Gondwana to GAAR landia: Evolutionary history and biogeography of ogre-faced spiders (*Deinopis*). J Biogeography, 45:2442–2457.

Chamberlin RV. 1917. New spiders of the family Aviculariidae. Bull. Mus. Comp. Zool., 61:25–75

Chamberlin RV, Ivie W. 1946. On several new American spiders. University of Utah.

Chen Z-Q, Benton MJ. 2012. The timing and pattern of biotic recovery following the end-Permian mass extinction. Nature Geoscience, 5:375.

Clouse RM, Branstetter MG, Buenavente P, Crowley LM, Czekanski-Moir J, General DEM, Giribet G, Harvey MS, Janies DA, Mohagan AB. 2017. First global molecular phylogeny and biogeographical analysis of two arachnid orders (Schizomida and Uropygi) supports a tropical Pangean origin and mid-Cretaceous diversification. J Biogeography, 44:2660–2672.

Coccioni R, Galeotti S. 1994. KT boundary extinction: Geologically instantaneous or gradual event? Evidence from deep-sea benthic foraminifera. Geology, 22:779–782.

Cock PJ, Antao T, Chang JT, Chapman BA, Cox CJ, Dalke A, Friedberg I, Hamelryck T, Kauff F, Wilczynski B. 2009. Biopython: freely available Python tools for computational molecular biology and bioinformatics. Bioinformatics, 25:1422–1423.

Coddington JA, Agnarsson I, Hamilton C, Bond JE. 2018. Spiders did not repeatedly gain, but repeatedly lost, foraging webs. PeerJ Preprints, 6:e27341v27341.

Coyle F. 1986. The role of silk in prey capture by nonaraneomorph spiders.

Coyle FA. 1971. Systematics and natural history of the mygalomorph spider genus Antrodiaetus and related genera (Araneae: Antrodiaetidae). Harvard Univ Mus Compar Zool Bull.

Coyle FA. 1981. The mygalomorph spider genus *Microhexura* (Araneae, Dipluridae). Bull. Am. Mus. Nat. Hist., 170:64–75.

Coyle FA. 1986. Chilehexops, a new funnelweb mygalomorph spider genus from Chile (Araneae, Dipluridae). American Museum Novitates; no. 2860.

Coyle FA. 1988. A revision of the American funnel web mygalomorph spider genus Euagrus (Araneae, Dipluridae). Bulletin of the American Museum of Natural History., 187:203–292.

Coyle FA. 1995. A revision of the funnelweb mygalomorph spider subfamily Ischnothelinae (Araneae, Dipluridae). Bulletin of the AMNH; no. 226.

Coyle FA, Dellinger RE, Bennet R. 1992. Retreat architecture and construction behavior of an East African idiopine trapdoor spider (Araneae, Idiopidae). Bulletin of the of the British Arachnological Society, 9:99–104.

Coyle FA, Ketner ND. 1990. Observations on the prey and prey capture behaviour of the funnelweb mygalomorph spider genus *Ischnothele* (Araneae, Dipluridae). Bull Br arachnol Soc, 8:97–104.

Cracraft J. 1988. Deep-history biogeography: retrieving the historical pattern of evolving continental biotas. Syst Biol, 37:221–236.

Decae AE. 1996. Systematics of the trapdoor spider genus *Cyrtocarenum* Ausserer, 1871 (Araneae, Ctenizidae). Bulletin of British arachnological Society, 10:161–170.

Decae AE, Bosmans R. 2014. Synonymy of the Trapdoor Spider Genera *Cyrtauchenius* Thorell, 1869 and Amblyocarenum Simon, 1892 Reconsidered (Araneae, Mygalomorphae, Cyrtaucheniidae). Arachnology, 16:182–192.

Dippenaar-Schoeman AS. 2002. Baboon and trapdoor spiders of Southern Africa: an identification manual. Agricultural Research Council Pretoria.

Domeier M, Torsvik TH. 2014. Plate tectonics in the late Paleozoic. Geoscience Frontiers, 5:303–350.

Eberhard WG, Hazzi NA. 2013. Web construction of *Linothele macrothelifera* (Araneae: Dipluridae). The Journal of Arachnology, 41:70–75.

Engelbrecht I, Prendini L. 2012. Cryptic diversity of South African trapdoor spiders: Three new species of *Stasimopus* Simon, 1892 (Mygalomorphae, Ctenizidae), and redescription of *Stasimopus robertsi* Hewitt, 1910. American Museum Novitates:1–42.

Eskov K, Zonshtein S. 1990. First Mesozoic mygalomorph spiders for the Lower Cretaceous of Siberia and Mongolia, with notes on the infraorder Mygalomorphae (Chelicerata: Araneae). Neue Jahrbuch für Geologie und Palaeontologie Abhandlungen, 178:325–368.

Eskov KY, Selden PA. 2005. First record of spiders from the Permian period (Araneae: Mesothelae). Bull Br arachnol Soc, 13:111–116.

Faircloth BC, McCormack JE, Crawford NG, Harvey MG, Brumfield RT, Glenn TC. 2012. Ultraconserved elements anchor thousands of genetic markers spanning multiple evolutionary timescales. Syst Biol, 61:717–726.

Fernández R, Hormiga G, Giribet G. 2014. Phylogenomic analysis of spiders reveals nonmonophyly of orb weavers. Current Biology, 24:1772–1777.

Fernández R, Kallal RJ, Dimitrov D, Ballesteros JA, Arnedo MA, Giribet G, Hormiga G. 2018. Phylogenomics, Diversification Dynamics, and Comparative Transcriptomics across the Spider Tree of Life. Current Biology, 28:1489–1497. e1485.

Forster R. 1968. The spiders of New Zealand. Part II. Ctenizidae, Dipluridae and Migidae. Otago Mus. Bull.

Frazão A, da Silva HR, de Moraes Russo CA. 2015. The Gondwana breakup and the history of the Atlantic and Indian oceans unveils two new clades for early neobatrachian diversification. PloS one, 10:e0143926.

Gabriel R, Longhorn SJ. 2015. Revised generic placement of Brachypelma embrithes (Chamberlin & Ivie, 1936) and Brachypelma angustum Valerio, 1980, with definition of the taxonomic features for identification of female Sericopelma Ausserer, 1875 (Araneae, Theraphosidae). ZooKeys:75.

Gallon RC. 2002. Revision of the African genera *Pterinochilus* and *Eucratoscelus* (Araneae, Theraphosidae, Harpactirinae) with description of two new genera. Bull Br arachnol Soc, 12:201–232.

Garrison NL, Rodriguez J, Agnarsson I, Coddington JA, Griswold CE, Hamilton CA, Hedin M, Kocot KM, Ledford JM, Bond JE. 2016. Spider phylogenomics: untangling the Spider Tree of Life. PeerJ, 4:e1719.

Garwood RJ, Dunlop JA, Selden PA, Spencer AR, Atwood RC, Vo NT, Drakopoulos M. 2016. Almost a spider: a 305-million-year-old fossil arachnid and spider origins. Proc. R. Soc. B, 283:20160125.

Gertsch WJ. 1979. A revision of the spider family Mecicobothriidae (Araneae, Mygalomorphae). American Museum novitates; no. 2687.

Gertsch WJ, Platnick NI. 1980. A revision of the American spiders of the family Atypidae (Araneae, Mygalomorphae). American Museum novitates; no. 2704.

Godwin RL, Opatova V, Garrison NL, Hamilton CA, Bond JE. 2018. Phylogeny of a cosmopolitan family of morphologically conserved trapdoor spiders (Mygalomorphae, Ctenizidae) using Anchored Hybrid Enrichment, with a description of the family, Halonoproctidae Pocock 1901. Mol.Phylogenet.Evol., 126:303–313.

Goloboff PA. 1993. A reanalysis of mygalomorph spider families (Araneae). American Museum Novitates, 3056:1–32.

Goloboff PA. 1995. A revision of the South American spiders of the family Nemesiidae (Araneae, Mygalomorphae): Part I: Species from Peru, Chile, Argentina, and Uruguay. Bull. Am. Mus. Nat. Hist.:1–189.

Golonka J. 2007. Late Triassic and Early Jurassic palaeogeography of the world. Palaeogeography, Palaeoclimatology, Palaeoecology, 244:297–307.

Graham MR, Hendrixson BE, Hamilton CA, Bond JE. 2015. Miocene extensional tectonics explain ancient patterns of diversification among turret-building tarantulas (*Aphonopelma mojave* group) in the Mojave and Sonoran deserts. J Biogeography, 42:1052–1065.

Gray M. 2010. A Revision of the Australian Funnel-web Spiders (Hexathelidae: Atracinae). Records of the Australian Museum, 62:285–392.

Griswold CE. 1985. A revision of the African spiders of the family Microstigmatidae (Araneae: Mygalomorphae). Annals of the Natal Museum, 27:1–37.

Griswold CE, Ledford J. 2001. A Monograph of the Migid Trap Door Spiders of Madagascar: And a Review of the World Genera (Araneae, Mygalomorphae, Migidae). California Academy of Sciences.

Guadanucci JPL. 2005. Tarsal scopula significance in ischnocolinae phylogenetics (Araneae, Mygalomorphae, Theraphosidae). J. Arachnol.:456–467.

Guadanucci JPL. 2011. Cladistic analysis and biogeography of the genus *Oligoxystre* Vellard 1924 (Araneae: Mygalomorphae: Theraphosidae). J. Arachnol., 39:320–326.

Guadanucci JPL. 2014. Theraphosidae phylogeny: relationships of the ‘Ischnocolinae’genera (Araneae, Mygalomorphae). Zool.Scr., 43:508–518.

Guadanucci JPL, Wendt I. 2014. Revision of the spider genus *Ischnocolus* Ausserer, 1871 (Mygalomorphae: Theraphosidae: Ischnocolinae). J. Nat. Hist., 48:387–402.

Hamelryck T, Manderick B. 2003. PDB file parser and structure class implemented in Python. Bioinformatics, 19:2308–2310.

Hamilton CA, Formanowicz DR, Bond JE. 2011. Species Delimitation and Phylogeography of *Aphonopelma hentzi* (Araneae, Mygalomorphae, Theraphosidae): Cryptic Diversity in North American Tarantulas. PLoS ONE, 6:e26207.

Hamilton CA, Hendrixson BE, Bond JE. 2016a. Taxonomic revision of the tarantula genus Aphonopelma Pocock, 1901 (Araneae, Mygalomorphae, Theraphosidae) within the United States. ZooKeys:1.

Hamilton CA, Hendrixson BE, Brewer MS, Bond JE. 2014. An evaluation of sampling effects on multiple DNA barcoding methods leads to an integrative approach for delimiting species: a case study of the North American tarantula genus *Aphonopelma* (Araneae, Mygalomorphae, Theraphosidae). Mol Phylogenet Evol, 71:79–93.

Hamilton CA, Lemmon AR, Lemmon EM, Bond JE. 2016b. Expanding anchored hybrid enrichment to resolve both deep and shallow relationships within the spider tree of life. BMC Evolutionary Biology, 16:212.

Harrison SE, Harvey MS, Cooper SJ, Austin AD, Rix MG. 2017. Across the Indian Ocean: A remarkable example of trans-oceanic dispersal in an austral mygalomorph spider. PloS one, 12:e0180139.

Harvey MS, Hillyer MJ, Main BY, Moulds TA, Raven RJ, Rix MG, Vink CJ, Huey JA. 2018. Phylogenetic relationships of the Australasian open-holed trapdoor spiders (Araneae: Mygalomorphae: Nemesiidae: Anaminae): multi-locus molecular analyses resolve the generic classification of a highly diverse fauna. Zool. J. Linn. Soc., 184:407–452

Hedin M, Bond JE. 2006. Molecular phylogenetics of the spider infraorder Mygalomorphae using nuclear rRNA genes (18S and 28S): Conflict and agreement with the current system of classification. Mol.Phylogenet.Evol., 41:454–471.

Hedin M, Carlson D, Coyle F. 2015. Sky island diversification meets the multispecies coalescent–divergence in the spruce-fir moss spider (*Microhexura montivaga*, Araneae, Mygalomorphae) on the highest peaks of southern Appalachia. Mol Ecol, 24:3467–3484.

Hedin M, Derkarabetian S, Alfaro A, Ramírez MJ, Bond JE. 2019. Phylogenomic analysis and revised classification of atypoid mygalomorph spiders (Araneae, Mygalomorphae), with notes on arachnid ultraconserved element loci. PeerJ 7:e6864 http://doi.org/10.7717/peerj.6864.

Hedin M, Derkarabetian S, Ramírez MJ, Vink C, Bond JE. 2018. Phylogenomic reclassification of the world’s most venomous spiders (Mygalomorphae, Atracinae), with implications for venom evolution. Scientific reports, 8:1636.

Hedin M, Starrett J, Hayashi C. 2013. Crossing the uncrossable: novel trans-valley biogeographic patterns revealed in the genetic history of low-dispersal mygalomorph spiders (Antrodiaetidae, *Antrodiaetus*) from California. Mol Ecol, 22:508–526.

Hendrixson BE, Bond JE. 2009. Evaluating the efficacy of continuous quantitative characters for reconstructing the phylogeny of a morphologically homogeneous spider taxon (Araneae, Mygalomorphae, Antrodiaetidae, *Antrodiaetus*). Mol.Phylogenet.Evol., 53:300–313.

Hendrixson BE, DeRussy BM, Hamilton CA, Bond JE. 2013. An exploration of species boundaries in turret-building tarantulas of the Mojave Desert (Araneae, Mygalomorphae, Theraphosidae, *Aphonopelma*). Mol Phylogenet Evol, 66:327–340.

Hewitt J. 1915. VI. Notes on several four-lunged spiders in the coliection of the Durban Museum with descriptions of two new forms. Durban Museum Novitates, 1:125–133.

Hewitt J. 1916. Descriptions of new South African spiders. Annals of the Transvaal Museum, 5:180–213.

Hughes N, McDougall A. 1990. New Wealden correlation for the Wessex basin. Proceedings of the Geologists’ Association, 101:85–90.

Indicatti RP, Lucas SM. 2005. Description of a new genus of Nemesiidae (Araneae, Mygalomorphae) from the Brazilian Cerrado. Zootaxa, 1088:11–16.

Isbister GK, Gray MR, Balit CR, Raven RJ, Stokes BJ, Porges K, Tankel AS, Turner E, White J, Fisher MM. 2005. Funnel-web spider bite: a systematic review of recorded clinical cases. The Medical Journal of Australia, 182:407–411.

Isbister GK, Sellors KV, Beckmann U, Chiew AL, Downes MA, Berling I. 2015. Catecholamine-induced cardiomyopathy resulting from life-threatening funnel-web spider envenoming. Med J Aust, 203:302–304.

Katoh K, Standley DM. 2013. MAFFT multiple sequence alignment software version 7: improvements in performance and usability. Mol Biol Evol, 30:772–780.

Kim SI, Farrell BD. 2015. Phylogeny of world stag beetles (Coleoptera: Lucanidae) reveals a Gondwanan origin of Darwin’s stag beetle. Mol.Phylogenet.Evol., 86:35–48.

Král J, Kořínková T, Krkavcová L, Musilová J, Forman M, Herrera IMÁ, Haddad CR, Vítková M, Henriques S, Vargas JGP. 2013. Evolution of karyotype, sex chromosomes, and meiosis in mygalomorph spiders (Araneae: Mygalomorphae). Biol. J. Linn. Soc., 109:377–408.

Kück P. 2009. ALICUT: a Perlscript which cuts ALISCORE identified RSS. Department of Bioinformatics, Zoologisches Forschungsmuseum A. Koenig (ZFMK), Bonn, Germany, version, 2.

Kück P, Longo GC. 2014. FASconCAT-G: extensive functions for multiple sequence alignment preparations concerning phylogenetic studies. Frontiers in Zoology, 11:81.

Kuntner M, Hamilton CA, Cheng R-C, Gregoric M, Lupse N, Lokovsek T, Lemmon E, Lemmon A, Agnarsson I, Coddington JA. 2018. Golden Orbweavers Ignore Biological Rules: Phylogenomic and Comparative Analyses Unravel a Complex Evolution of Sexual Size Dimorphism. bioRxiv:368233.

Lanfear R, Calcott B, Kainer D, Mayer C, Stamatakis A. 2014. Selecting optimal partitioning schemes for phylogenomic datasets. BMC evolutionary biology, 14:82.

Lanfear R, Frandsen PB, Wright AM, Senfeld T, Calcott B. 2016. PartitionFinder 2: new methods for selecting partitioned models of evolution for molecular and morphological phylogenetic analyses. Mol Biol Evol, 34:772–773.

Leavitt DH, Starrett J, Westphal MF, Hedin M. 2015. Multilocus sequence data reveal dozens of putative cryptic species in a radiation of endemic Californian mygalomorph spiders (Araneae, Mygalomorphae, Nemesiidae). Mol.Phylogenet.Evol., 91:56–67.

Lemmon AR, Emme SA, Lemmon EM. 2012. Anchored Hybrid Enrichment for Massively High-Throughput Phylogenomics. Syst Biol, 61:727–744.

Li J-T, Wang J-S, Nian H-H, Litvinchuk SN, Wang J, Li Y, Rao D-Q, Klaus S. 2015. Amphibians crossing the bering land bridge: evidence from holarctic treefrogs (*Hyla*, Hylidae, Anura). Mol.Phylogenet.Evol., 87:80–90.

Lüddecke T, Krehenwinkel H, Canning G, Glaw F, Longhorn SJ, Taenzler R, Wendt I, Vences M. 2018. Discovering the silk road: Nuclear and mitochondrial sequence data resolve the phylogenetic relationships among theraphosid spider subfamilies. Mol.Phylogenet.Evol., 119:63–70.

Maddison WP, Evans SC, Hamilton CA, Bond JE, Lemmon AR, Lemmon EM. 2017. A genome-wide phylogeny of jumping spiders (Araneae, Salticidae), using anchored hybrid enrichment. ZooKeys:89.

Maguilla E, Escudero M, Luceño M. 2018. Vicariance versus dispersal across Beringian land bridges to explain circumpolar distribution: A case study in plants with high dispersal potential. J Biogeography, 45:771–783.

Main B. 1957. Occurrence of the trap-door spider *Conothele malayana* (Doleschall) in Australia (Mygalomorphae: Ctenizidae). Western Australian Naturalist, 5:209–216.

Main B. 1993. Biogeographic significance of the Nullarbor cave mygalomorph spider *Troglodiplura* and its taxonomic affinities. Journal of the Royal Society of Western Australia, 76:77–85.

Main BY. 1969. A blind mygalomorph spider from a Nullarbor plain cave. Roy Soc West Australia J. 52:9–11

Main BY, Mascord R. 1969. A new genus of diplurid spider (Araneae: Mygalomorphae) from New South Wales. Journal of the Entomological Society of Australia (NSW), 6:24.

Matthews KJ, Maloney KT, Zahirovic S, Williams SE, Seton M, Mueller RD. 2016. Global plate boundary evolution and kinematics since the late Paleozoic. Global and Planetary Change, 146:226–250.

Milne R. 2006. Northern hemisphere plant disjunctions: a window on Tertiary land bridges and climate change? Annals of Botany, 98:465–472.

Mirarab S, Warnow T. 2015. ASTRAL-II: coalescent-based species tree estimation with many hundreds of taxa and thousands of genes. Bioinformatics, 31:i44–i52.

Misof B, Misof K. 2009. A Monte Carlo approach successfully identifies randomness in multiple sequence alignments: a more objective means of data exclusion. Syst Biol, 58:21–34.

Mittermeier RA, Turner WR, Larsen FW, Brooks TM, Gascon C. 2011. Global biodiversity conservation: the critical role of hotspots. Biodiversity hotspots, Springer, p. 3–22.

Montes de Oca L, D’Elía G, Pérez-Miles F. 2016. An integrative approach for species delimitation in the spider genus *Grammostola* (Theraphosidae, Mygalomorphae). Zool.Scr., 45:322–333.

Mora E, Paspati A, Decae AE, Arnedo MA. 2017. Rafting spiders or drifting islands? Origins and diversification of the endemic trap-door spiders from the Balearic Islands, Western Mediterranean. J Biogeography, 44:924–936.

Morrone JJ, Crisci JV. 1995. Historical biogeography: introduction to methods. Annu Rev Ecol Evol Systemat, 26:373–401.

Murienne J, Daniels SR, Buckley TR, Mayer G, Giribet G. 2014. A living fossil tale of Pangaean biogeography. Proceedings of the Royal Society of London B: Biological Sciences, 281:20132648.

Myers N, Mittermeier RA, Mittermeier CC, da Fonseca GA, Kent J. 2000. Biodiversity hotspots for conservation priorities. Nature, 403:853–858.

Nance RD, Murphy JB, Santosh M. 2014. The supercontinent cycle: a retrospective essay. Gondwana Research, 25:4–29.

Nentwig W. 1983. The prey of web-building spiders compared with feeding experiments (Araneae: Araneidae, Linyphiidae, Pholcidae, Agelenidae). Oecologia, 56:132–139.

Nguyen LT, Schmidt HA, von Haeseler A, Minh BQ. 2014. IQ-TREE: a fast and effective stochastic algorithm for estimating maximum-likelihood phylogenies. Mol Biol Evol, 32:268–274.

Opatova V, Arnedo MA. 2014a. From Gondwana to Europe: inferring the origins of Mediterranean *Macrothele* spiders (Araneae: Hexathelidae) and the limits of the family Hexathelidae. Invertebr Systemat, 28:361–374.

Opatova V, Arnedo MA. 2014b. Spiders on a Hot Volcanic Roof: Colonisation Pathways and Phylogeography of the Canary Islands Endemic Trap-Door Spider *Titanidiops canariensis* (Araneae, Idiopidae). PLoS ONE, 9:e115078. doi:115010.111371/journal.pone.0115078.

Opatova V, Bond JE, Arnedo MA. 2013. Ancient origins of the Mediterranean trap-door spiders of the family Ctenizidae (Araneae, Mygalomorphae). Mol Phylogenet Evol, 69:1135–1145.

Opatova V, Bond JE, Arnedo MA. 2016. Uncovering the role of the Western Mediterranean tectonics in shaping the diversity and distribution of the trap-door spider genus Ummidia (Araneae, Ctenizidae). J Biogeography, 43:1955–1966.

Ortiz D, Francke OF, Bond JE. 2018. A tangle of forms and phylogeny: Extensive morphological homoplasy and molecular clock heterogeneity in *Bonnetina* and related tarantulas. Mol.Phylogenet.Evol., 127:55–73.

Ott R, Höfer H. 2003. *Envia garciai*, a new genus and species of mygalomorph spiders (Araneae, Microstigmatidae) from Brazilian Amazonia. Iheringia. Série Zoologia, 93:373–379.

Panchen AL. 1992. Classification, evolution, and the nature of biology. Cambridge University Press.

Passanha V, Brescovit AD. 2018. On the Neotropical spider Subfamily Masteriinae (Araneae, Dipluridae). Zootaxa, 4463:1–73.

Pattengale ND, Alipour M, Bininda-Emonds ORP, Moret BME, Stamatakis A. 2010. How many bootstrap replicates are necessary? Journal of Computational Biology, 17:337–354.

Pérez-Miles F. 1994. Tarsal scopula division in Theraphosinae (Araneae, Theraphosidae): its systematic significance. J. Arachnol.:46–53.

Pérez-Miles F. 2002. The occurrence of abdominal urticating hairs during development in Theraphosinae (Araneae, Theraphosidae): phylogenetic implications. J. Arachnol., 30:316–320.

Pérez-Miles F, Lucas S, da Silva P Jr, Bertani R. 1996.

Prum RO, Berv JS, Dornburg A, Field DJ, Townsend JP, Lemmon EM, Lemmon AR. 2015. A comprehensive phylogeny of birds (Aves) using targeted next-generation DNA sequencing. Nature, 526:569.

Raven R. 1991. A revision of the mygalomorph spider family Dipluridae in New Caledonia (Araneae). Mémoires du Muséum National d’Histoire Naturelle, Paris (A), 149:87–117.

Raven RJ, Schwendinger P. 1995. Three new mygalomorph spider genera from Thailand and China (Araneae). Memoirs-Queenslad Museum, 38:623–642.

Raven RJ. 1980. The evolution and biogeography of the mygalomorph spider family Hexathelidae (Araneae, Chelicerata). J. Arachnol., 8:251–266.

Raven RJ. 1981. A Review of the Australian Genera of the Mygalomorph Spider Subfamily Diplurinae (Dipluridae: Chelicerata). Aust. J. Zool., 29:321–363.

Raven RJ. 1985. The spider infraorder Mygalomorphae (Araneae): cladistics and systematics. Bull. Amer. Mus. Nat. Hist., 182:1–180.

Raven RJ. 1994. Mygalomorph spiders of the Barychelidae in Australia and the western Pacific. Mem. Queensl. Mus., 35:291–706.

Ree RH, Smith SA. 2008. Maximum Likelihood Inference of Geographic Range Evolution by Dispersal, Local Extinction, and Cladogenesis. Syst Biol, 57:4–14.

Ríos Tamayo D. 2017. Re-description of *Trichopelma cubanum* (Theraphosidae: Ischnocolinae) and comments about the familial placement of Trichopelma. Mun. Ent. Zool., 12:194–198.

Ríos-Tamayo D, Goloboff P. 2017. *Vilchura calderoni*, a new genus and species of Euagrinae (Araneae: Mygalomorphae: Dipluridae) from Chile. Arachnology, 17:183–187.

Rix MG, Bain K, Main BY, Raven RJ, Austin AD, Cooper SJ, Harvey MS. 2017a. Systematics of the spiny trapdoor spiders of the genus *Cataxia* (Mygalomorphae: Idiopidae) from south-western Australia: documenting a threatened fauna in a sky-island landscape. The Journal of Arachnology, 45:395–423.

Rix MG, Cooper SJ, Meusemann K, Klopfstein S, Harrison SE, Harvey MS, Austin AD. 2017b. Post-Eocene climate change across continental Australia and the diversification of Australasian spiny trapdoor spiders (Idiopidae: Arbanitinae). Mol.Phylogenet.Evol., 109:302–320.

Rix MG, Raven RJ, Main BY, Harrison SE, Austin AD, Cooper SJ, Harvey MS. 2017c. The Australasian spiny trapdoor spiders of the family Idiopidae (Mygalomorphae: Arbanitinae): a relimitation and revision at the generic level. Invertebr Systemat, 31:566–634.

Sanderson MJ. 2002. Estimating absolute rates of molecular evolution and divergence times: a penalized likelihood approach. Mol Biol Evol, 19:101–109.

Sanmartín I, Enghoff H, Ronquist F. 2001. Patterns of animal dispersal, vicariance and diversification in the Holarctic. Biol. J. Linn. Soc., 73:345–390.

Sanmartín I, Ronquist F. 2004. Southern hemisphere biogeography inferred by event-based models: plant versus animal patterns. Syst Biol, 53:216–243.

Satler JD, Carstens BC, Hedin M. 2013. Multilocus Species Delimitation in a Complex of Morphologically Conserved Trapdoor Spiders (Mygalomorphae, Antrodiaetidae, *Aliatypus*). Syst Biol, 62:805–823.

Schmidt G. 2002. Gehören *Brachionopus* Pocock, 1897 und *Harpactirella* Purcell, 1902 zu den Theraphosiden. Arthropoda, 10:12–17.

Selden PA. 2002. First British Mesozoic Spider, From Cretaceous Amber Of The Isle Of Wight, Southern England. Palaeontology, 45:973–983.

Selden PA, Gall JC. 1992. A Triassic mygalomorph spider from the northern Vosges, France. Palaeontology, 35:211–235.

Selden PA, Shcherbakov DE, Dunlop JA, Eskov KY. 2014. Arachnids from the Carboniferous of Russia and Ukraine, and the Permian of Kazakhstan. Paläontologische Zeitschrift, 88:297–307.

Sharma PP, Baker CM, Cosgrove JG, Johnson JE, Oberski JT, Raven RJ, Harvey MS, Boyer SL, Giribet G. 2018. A revised dated phylogeny of scorpions: Phylogenomic support for ancient divergence of the temperate Gondwanan family Bothriuridae. Mol.Phylogenet.Evol., 122:37–45.

Shimodaira H. 2002. An approximately unbiased test of phylogenetic tree selection. Syst Biol, 51:492–508.

Shimodaira H, Hasegawa M. 2001. CONSEL: for assessing the confidence of phylogenetic tree selection. Bioinformatics, 17:1246–1247.

Shimojana M, Haupt J. 1998. Taxonomy and natural history of the funnel-web spider genus *Macrothele* (Araneae: Hexathelidae: Macrothelinae) in the Ryukyu Islands (Japan) and Taiwan. Species Diversity, 3:1–15.

Shultz JW. 1987. The origin of the spinning apparatus in spiders. Biological Reviews, 62:89–113.

Simon E. 1864. Histoire naturelle des Araignées (Aranéides). Paris, Roret edit.

Simon E. 1889. Voyage de M. E. Simon au Venezuela (Decembre 1887–Avril 1888) 4e Mémoire.. Annales de la Société Entomologique de France, Paris, p. 169–220.

Smith SA. 2010. Lagrange C++ Manual.

Smith SA, O’Meara BC. 2012. treePL: divergence time estimation using penalized likelihood for large phylogenies. Bioinformatics, 28:2689–2690.

Snazell RG, Allison R. 1989. The genus *Macrothele* Ausserer (Araneae; Hexathelidae) in Europe. Bull Br arachnol Soc, 8:65–72.

Stamatakis A. 2014. RAxML version 8: a tool for phylogenetic analysis and post-analysis of large phylogenies. Bioinformatics, 30:1312–1313.

Starrett J, Derkarabetian S, Hedin M, Bryson RW Jr, McCormack JE, Faircloth BC. 2017. High phylogenetic utility of an ultraconserved element probe set designed for Arachnida. Mol. Ecol. Res., 17:812–823.

Starrett J, Hayashi CY, Derkarabetian S, Hedin M. 2018. Cryptic elevational zonation in trapdoor spiders (Araneae, Antrodiaetidae, *Aliatypus janus* complex) from the California southern Sierra Nevada. Mol.Phylogenet.Evol., 118:403–413.

Starrett J, Hedin M, Ayoub N, Hayashi CY. 2013. Hemocyanin gene family evolution in spiders (Araneae), with implications for phylogenetic relationships and divergence times in the infraorder Mygalomorphae. Gene, 524:175–186.

Su YC, Brown RM, Chang YH, Lin CP, Tso IM. 2016. Did a Miocene–Pliocene island isolation sequence structure diversification of funnel web spiders in the Taiwan-Ryukyu Archipelago? J Biogeography, 43:991–1003.

Swofford D. 2003. PAUP*. Phylogenetic Analysis Using Parsimony (*and Other Methods), version 4.0b10.

Tiffney, Manchester. 2001. The use of geological and paleontological evidence in evaluating plant phylogeographic hypotheses in the Northern Hemisphere tertiary. International Journal of Plant Sciences, 162:S3–S17.

Toussaint EF, Bloom D, Short AE. 2017. Cretaceous West Gondwana vicariance shaped giant water scavenger beetle biogeography. J Biogeography, 44:1952–1965.

Turner SP, Longhorn SJ, Hamilton CA, Gabriel R, Pérez-Miles F, Vogler AP. 2018. Re-evaluating conservation priorities of New World tarantulas (Araneae: Theraphosidae) in a molecular framework indicates non-monophyly of the genera, *Aphonopelma* and *Brachypelma*. Systematics and Biodiversity, 16:89–107.

Veevers J. 2004. Gondwanaland from 650–500 Ma assembly through 320 Ma merger in Pangea to 185–100 Ma breakup: supercontinental tectonics via stratigraphy and radiometric dating. Earth-Science Reviews, 68:1–132.

Vollrath F, Selden P. 2007. The Role of Behavior in the Evolution of Spiders, Silks, and Webs. Annu Rev Ecol Evol Systemat, 38:819–846.

Wang B, Dunlop JA, Selden PA, Garwood RJ, Shear WA, Müller P, Lei X. 2018. Cretaceous arachnid *Chimerarachne yingi* gen. et sp. nov. illuminates spider origins. Nature ecology & evolution, 2:614.

Wheeler WC, Coddington JA, Crowley LM, Dimitrov D, Goloboff PA, Griswold CE, Hormiga G, Prendini L, Ramírez MJ, Sierwald P. 2017. The spider tree of life: phylogeny of Araneae based on target-gene analyses from an extensive taxon sampling. Cladistics, 33:574–616.

Wiley EO. 1988. Vicariance biogeography. Annu Rev Ecol Evol Systemat, 19:513–542.

Will TM, Frimmel HE. 2018. Where does a continent prefer to break up? Some lessons from the South Atlantic margins. Gondwana Research, 53:9–19.

Wood HM, González VL, Lloyd M, Coddington J, Scharff N. 2018. Next-generation museum genomics: Phylogenetic relationships among palpimanoid spiders using sequence capture techniques (Araneae: Palpimanoidea). Mol.Phylogenet.Evol., 127:907–918.

Wood HM, Matzke NJ, Gillespie RG, Griswold CE. 2012. Treating fossils as terminal taxa in divergence time estimation reveals ancient vicariance patterns in the palpimanoid spiders. Syst Biol, 62:264–284.

World Spider Catalog. 2018. World Spider Catalog (2018). World Spider Catalog. Version 19.5. Natural History Museum Bern, online at http://wsc.nmbe.ch/, accessed on {16 November 2018}. doi: 10.24436/2

Wunderlich Jh. 2011. Some fossil spiders (Araneae) in Eocene European ambers. In: Wunderlich J editor. Beiträge zur Araneologie, p. 472–538.

Xu X, Liu F, Cheng R-C, Chen J, Xu X, Zhang Z, Ono H, Pham DS, Norma-Rashid Y, Arnedo MA. 2015. Extant primitively segmented spiders have recently diversified from an ancient lineage. Proc. R. Soc. B, 282:20142486.

Young A, Flament N, Maloney K, Williams S, Matthews K, Zahirovic S, Müller RD. 2018. Global kinematics of tectonic plates and subduction zones since the late Paleozoic Era. Geoscience Frontiers. https://doi.org/10.1016/j.gsf.2018.05.011

Yu Y, Harris AJ, Blair C, He X. 2015. RASP (Reconstruct Ancestral State in Phylogenies): a tool for historical biogeography. Mol.Phylogenet.Evol., 87:46–49.

Zachos J, Pagani M, Sloan L, Thomas E, Billups K. 2001. Trends, Rhythms, and Aberrations in Global Climate 65 Ma to Present. Science, 292:686–693.

